# Comparative population pangenomes reveal unexpected complexity and fitness effects of structural variants

**DOI:** 10.1101/2025.02.11.637762

**Authors:** Scott V. Edwards, Bohao Fang, Danielle Khost, George E Kolyfetis, Rebecca G Cheek, Devon A DeRaad, Nancy Chen, John W Fitzpatrick, John E. McCormack, W. Chris Funk, Cameron K Ghalambor, Erik Garrison, Andrea Guarracino, Heng Li, Timothy B Sackton

## Abstract

Structural variants (SVs) are widespread in vertebrate genomes, yet their evolutionary dynamics remain poorly understood. Using 45 long-read de novo genome assemblies and pangenome tools, we analyze SVs within three closely related species of North American jays (*Aphelocoma*, scrub-jays) displaying a 60-fold range in effective population size. We find rapid evolution of genome architecture, including ∼100 Mb variation in genome size driven by dynamic satellite landscapes with unexpectedly long (> 10 kb) repeat units and widespread variation in gene content, influencing gene expression. SVs exhibit slightly deleterious dynamics modulated by variant length and population size, with strong evidence of adaptive fixation only in large populations. Our results demonstrate how population size shapes the distribution of SVs and the importance of pangenomes to characterizing genomic diversity.

## Main text

A fundamental challenge of genome evolution is to understand how diverse mutational mechanisms alter genome architecture and the degree to which the observed variation is adaptive, neutral, or deleterious with respect to fitness (Ohta 2002, Lynch 2007). Avian genomes are widely considered the simplest genomes among amniotes (birds, non-avian reptiles and mammals), exhibiting broad interchromosomal synteny and a conservative mode of structural evolution compared to mammals and non-avian reptiles (Ellegren 2013, Zhang et al. 2014, Bravo et al. 2021, Galbraith et al. 2021, Griffin et al. 2024). Previous comparative genomic surveys have revealed that avian genomes exhibit a depauperate repeat landscape, with a few dramatic examples of expansions of transposable elements and satellites emerging in specific lineages (Manthey et al. 2018, Stiller et al. 2019), as well as largely congruent gene content (Griffin et al. 2024), with a few noteworthy examples of gene duplication and loss providing substrates for adaptive evolution (Yuri et al. 2008, Feng et al. 2020). However, most such analyses have been conducted with short-read sequencing data and reference-based bioinformatic tools, which have well-known biases (Günther et al. 2019, Martiniano et al. 2020, Lin et al. 2024) and are less effective at capturing complex types of variation, such as structural variants (SVs): insertions, deletions, inversions and other complex multi-nucleotide variants (Eizenga et al. 2020, Li et al. 2020, Andreace et al. 2023). The biases induced by reference-based analyses, as well as the heterogeneity of laboratory and bioinformatic pipelines between studies, make comparisons of SV abundance and dynamics among species difficult. Furthermore, most studies of SV dynamics in natural populations, with a few recent exceptions (Barton et al. 2019, Fang et al. 2024), are focused on the adaptive potential of selected SVs, such as inversions (Knief et al. 2016, Funk et al. 2021, Knief et al. 2024). Thus, we still lack a comprehensive genome-wide understanding of the distribution of fitness effects across the full spectrum of SVs at the population level.

Pangenomes can capture the full spectrum of genetic variation, including SVs, among individuals of a species, or among species (Paten et al. 2017, Eizenga et al. 2020, Secomandi et al. 2025). By comparing all assemblies to one another, without designating a reference assembly, pangenome approaches avoid many of the problems associated with reference-based variant calling, especially for SVs, which can be completely missed if query sequences contain genomic regions not found in the reference. Pangenomes have been a useful framework for understanding variation in gene and repeat content and SVs among individuals, species and strains, particularly among bacteria and domesticated animals and plants (Rouli et al. 2015, Bayer et al. 2020, Rosconi et al. 2022, Leonard et al. 2023, Wang et al. 2023, Schreiber et al. 2024). Among vertebrates, humans are the best-studied species in terms of pangenomes; the Human Pangenome Consortium has shown how a draft pangenome, consisting of 47 long-read phased diploid assemblies, captures more SVs and provides better mapping rates and accuracies than protocols involving a single reference assembly (Liao et al. 2023). Pangenomes are known to be more effective at capturing SVs when built with long-read genomes assembled de novo, yet we have few examples of data sets of the size and quality of the human draft pangenome. We also know little about the effectiveness of pangenome tools among closely related species (Li et al. 2020), especially in vertebrates.

Here we present a pangenome analysis of genomic variation among 45 long-read phased assemblies of North American scrub-jays in the genus *Aphelocoma*, as well as additional related species. Scrub-jays are a model system for studies in biogeography, speciation, life history, ecology, and local adaptation, and have been the subject of decades-long studies of social behavior (Woolfenden et al. 1984, McCormack et al. 2011, Chen et al. 2016, Aguillon et al. 2017, Chen et al. 2019, DeRaad et al. 2022). We chose these birds because their phylogeny, phylogeography and relative amounts of SNP diversity are already well understood and show a wide spectrum of levels of genetic variation that provide a clear context for interpreting the evolution of SVs (Peterson 1992, Brown et al. 1995, DeRaad et al. 2022). We focus on the Florida (*A. coerulescens*, AC), Island (*A. insularis*, AI) and Woodhouse’s (A*. woodehouseii*, AW) Scrub-jays; whereas AC occurs exclusively in Florida and AI only on Santa Cruz Island, California, AW is widely distributed across western North America (Fig. 1A). Recent investigations in corvids (Corvidae), including jays, have begun to reveal complex and rapid dynamics of satellite and repeat evolution in this group, albeit without the advantages of comprehensive long-read data or pangenome tools (Weissensteiner et al. 2017, Weissensteiner et al. 2020, Peona et al. 2023). Our study focuses on comparative population genomics, wherein genetic diversity and dynamics are compared across a suite of related species to understand the drivers of interspecific variation in genomic diversity (Ellegren et al. 2016, Edwards et al. 2021). Most (67%) of the genomic resources for this work were purpose-collected for this project, allowing us to maximize data quality and reproducibility.

**Figure 1.**
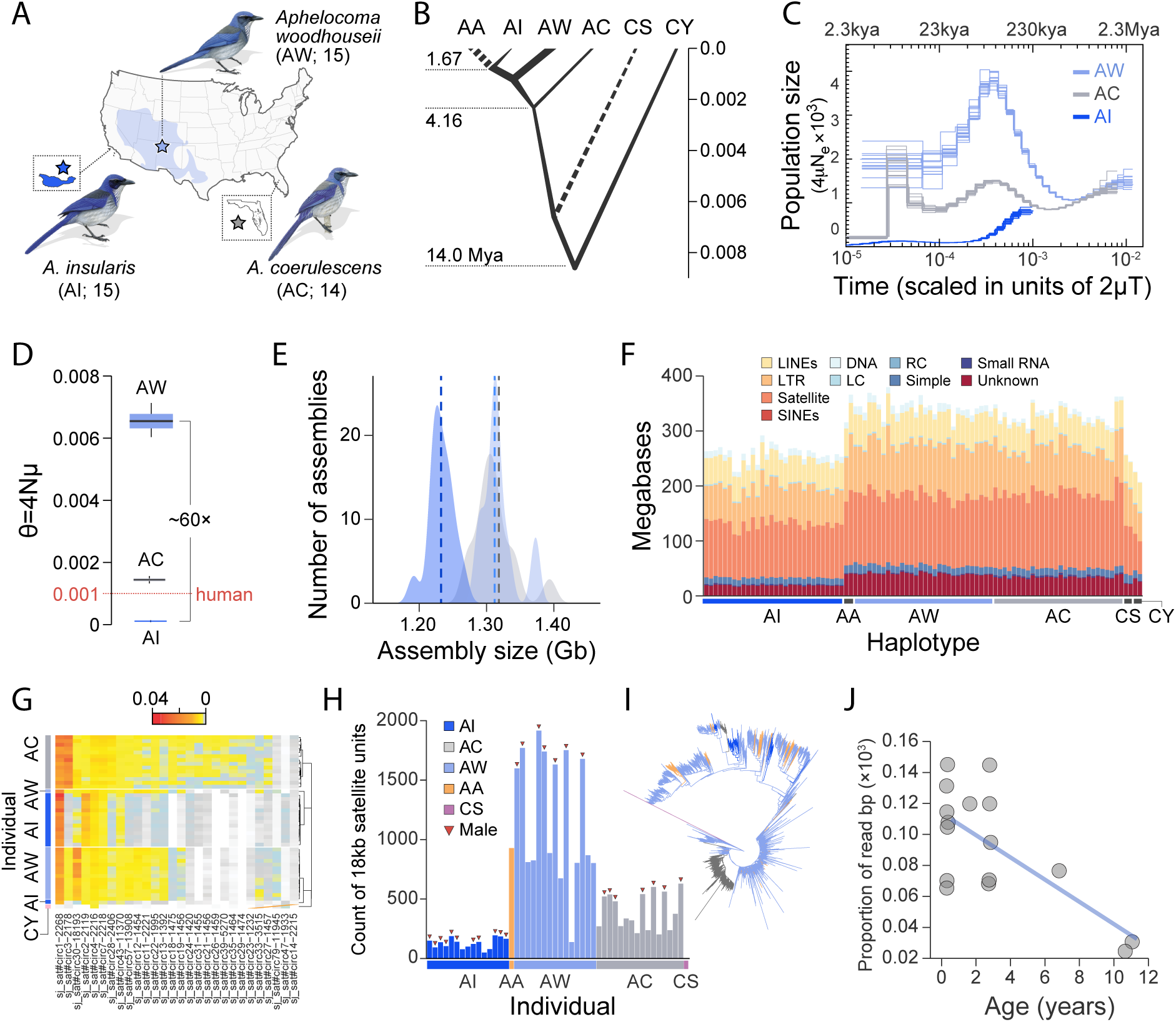
Demographic and genomic overview. **(A)** Distributions of Florida (*A. coerulescens*, AC), Island (*A. insularis*, AI), and Woodhouse’s (*A. woodehouseii*, AW) Scrub-Jays within the United States. Bird illustrations, © Lynx Edicions. (**B**) Species tree with branch lengths and widths estimated with bpp, with approximate divergence times (Mya). **(C)** PSMC-inferred demographic histories for AI, AC, and AW. **(D)** Genome-wide heterozygosity from bpp, with human shown in red as reference. (**E**) Distribution of assembly sizes (Gb) for the 90 haplotypes (45 diploid individuals). **(F)** Repeat content for the three species and two outgroups (CS=Steller’s Jay, CY=Yucatán Jay). **(G)** Heatmap of the 28 most abundant satellites across all haplotypes. **(H)** Expansion of satellite sj_sat#circ30_18193 (18.2 kb unit repeats) on the Z chr. of AW. **(I)** Neighbor-joining tree of ∼3500 proxies for ∼27,000 satellite monomers showing partial homogenization. **(J)** Telomere abundance versus age in known-age AC individuals. The regression (p = 0.000091, R^2^ = 0.68) suggests that there is a proportional decrease (β) in telomere abundance of −9.793e-06 per year.

### Geographic, genomic and demographic background

Our study is based on multiple blood and tissue samples rapidly cryo-preserved from AW in New Mexico (n=15 individuals), AI from Santa Cruz Island (n=15), AC from Florida (n=14; Fig. 1A, table S1). An individual from a related genus, the Yucatán Jay (*Cyanocorax yucatanicus*, CY, n=1) served as an outgroup for polarizing variants. To these we added other long-read jay genomes for various analyses as they became available, including one California Scrub-Jay (*A. californica*, AA (DeRaad et al. 2023)) and one Steller’s Jay (*Cyanocitta cristata*, CS (Benham et al. 2023), all with a well-known phylogeny (Fernando et al. 2017) (Fig. 1B). All populations were sampled from what can be reasonably considered single localities/regions without barriers to gene flow, and close relatives were avoided when known. We sequenced each sample with PacBio-HiFi to 29.6X – 61.4X coverage (average 43.7X; table S2) and generated 90 phased diploid assemblies with hifiasm (Cheng et al. 2021). We improved the assembly of an AW male and female with HiC (fig. S1), which served as reference genomes for purposes of annotation. The AW female reference was annotated using TOGA (Kirilenko et al. 2023) projections from chicken and zebra finch, which were collapsed into a single non-redundant annotation of high completeness (97.2% overall; 96.6% single, 0.6% duplicated). Blood samples generally yielded more highly contiguous assemblies, whose N50 without scaffolding by HiC ranged from 5.1 to 19 Mb (fig. S2, table S2). Karyotypes (2N = 80) helped clarify the number of macro- and microchromosomes (fig. S1).

To confirm and refine the known demographic background of this species complex (DeRaad et al. 2022), we focused on regions of high-mapping quality (Li et al. 2018), comprising approximately 85% of the AW reference genome (fig. S3). Individuals of each species were confirmed to be unrelated (fig. S4). PSMC (Li et al. 2011) and bpp (Rannala et al. 2017) confirmed the rank order of nucleotide diversity among *Aphelocoma* species as AW > AC > AI (Fig. 1C,D; fig. S5); AW birds have an effective population size ∼60 times that of AI birds, assuming similar mutation rates. AI birds exhibit longer runs of homozygosity (fig. S6) and nearly 70% of haplotype trees as monophyletic, as expected for a species that experienced a strong bottleneck (fig. S7, table S3). We found no evidence for gene flow between AW and AC birds (Materials, McCormack et al. 2011, DeRaad et al. 2022) (fig. S8), as expected given their wholly allopatric distributions today.

Our long-read assemblies revealed a conspicuous difference in assembly size among species: the average primary assembly sizes of AI birds (1.23 Gb) and the CY outgroup (1.19 Gb) were ∼79.6 Mb and ∼132 Mb shorter, respectively, than for AW (1.32 Gb) and AC (1.31 Gb) birds (Fig. 1E, fig. S9). Heterozygosity is known to influence the sizes of primary and locally phased haplotype assemblies due to the collapse of similar sequence into single regions when there are few differences between haplotypes, as in the AI birds (Cheng et al. 2021, Rhie et al. 2021). However, multiple lines of evidence suggest that variation in heterozygosity leading to technical artifacts is not the primary driver of assembly size in our data. The assemblies show no significant differences in size of the single copy fraction of the genome, which all fall between 955 Mb (AI) and 974 Mb (CY; fig. S10). Instead, multiple different satellites and repeat types show significant differences among species and chromosomes (Fig. 1F, fig. S11,S12). We annotated repeats using RepeatModeler2, RepeatMasker (Smit et al. 2015, Flynn et al. 2020) and Satellite Repeat Finder (Zhang et al. 2023), and compared the abundance of different repeat classes among species. We find a large and significant increase in the amount of LTRs, LINEs and total satellites in AW and AC assemblies compared to AI, CY, and CS, consistent with differences in assembly size (Fig 1F), yet some common satellites are more abundant in AI assemblies than in the other species (see below). Satellite DNA is difficult to assemble faithfully even when using HiFi reads (Huang et al. 2023, Peona et al. 2023); nonetheless, we found a strong correlation between the abundance of individual satellites per bird in reads and the abundance recovered from assemblies (fig. S13), suggesting that assembly size differences are not artifacts of heterozygosity or assembly protocol.

Despite intensive study of increasingly diverse species, including corvids (Weissensteiner et al. 2017, Peona et al. 2023), satellite DNA is still the least known component of the repetitive fraction of avian genomes. Satellite DNA was estimated to be the most common repeat type in the repetitive landscape of *Aphelocoma*, comprising between 37.1% (AW) and 41.5% (AC) of the total repeat landscape (Fig. 1F, table S4). Despite moderately low heterozygosity, AC assemblies possessed the highest total satellite abundance (mean 137.2 Mb per haplotype) compared to AW (mean 127.6 Mb) and AI assemblies (mean 98.9 Mb), much more than the CY and CS outgroups (63.1 and 88.0 Mb, respectively) (Fig. 1F). Satellite Repeat Finder (Zhang et al. 2023) detected a total of 300 distinct satellites across species, 28 of which had a per base pair abundance greater than 0.001; these core satellites had a range of unit lengths from 1232 bp to 18193 bp, with four > 10 kb (Fig. 1G). The five most common satellites across species included the 18.2 kb unit repeat satellite (sj_sat#circ30-18193) and four others with unit repeats of 2.2 - 2.3 kb; together these five satellites comprised 65% of all satellite-annotated base pairs across primary assemblies (53% of reads; fig. S13). Fourteen of the 28 satellites possessed > 80% similarity to satellites found in other corvids, suggesting long-term persistence across at least 20 MYA (Jønsson et al. 2016). However, the unit lengths of satellites detected by SRF were much longer than those detected in other corvids; two previously detected satellites in birds-of-paradise had similar unit lengths to those found in jays, but otherwise the unit lengths of jay satellites were 15 - ∼600 times longer than previously reported (table S5). Arrays of scrub-jay satellites tended not to have a higher-order internal structure like some human satellites (fig. S14). Mirroring results from other corvids (Peona et al. 2023), we found numerous examples of rapid shifts in satellite abundance between species. In both reads and assemblies, the smaller-genomed AI birds had the highest abundances of 3 of the 28 major satellites than in AW and AC, including the highly abundant sj_sat#circ1-2268 satellite (fig. S15). There were no full length copies of satellite sj_sat#circ30-18193 in CY, and only a single full length copy in CS, yet this satellite underwent a dramatic expansion in *Aphelocoma*, most pronounced on the Z chromosome of AW birds (Fig. 1H): male AW birds harbor an average of 500 more copies than do AW females. The phylogenetic relationships of ∼27,000 full length 18-kb unit repeats (Fig. 1I, fig. S16) suggest that gene conversion among satellite unit repeats has not yet fully homogenized unit-repeat clades within species, as seen in some mammals (Dover 1982, Rudd et al. 2006, Thakur et al. 2021).

We sequenced to high coverage two additional older (> 11 years) AC birds to demonstrate that telomeres show an expected decrease in length with age in the AC population, as measured from assemblies or from HiFi reads (Fig. 1J, table S6). Additionally, the AI birds have on average marginally lower telomere abundance than the other two species (fig. S17), a pattern consistent with recent theory predicting shorter telomeres in species with smaller effective population sizes as a consequence of fixation of deleterious and shorter telomere alleles (Brown et al. 2024). However, we cannot rule out that these differences are here driven by differences in average individual ages sampled across species. We also found subtle differences in GC content among species, attributable largely to differences in the abundance of specific satellite repeats (fig. S18). Overall, the data suggest a scenario in which the abundances of multiple satellites and transposable elements exploded in the ancestor of *Aphelocoma* and then were differentially reduced in AI, shifting its base composition, perhaps on its founding of Santa Cruz Island or through later drift.

### Pangenome graphs and structural variants

We used the Pangenome Graph Builder (Garrison et al. 2024) and minigraph (Li et al. 2020) to build our primary pangenome graphs (Materials) (fig. S19). PGGB first builds all-versus-all communities of related sequences; each community generally corresponds to a distinct chromosome. By contrast, minigraph builds a pangenome graph sequentially, adding sequences from each haplotype in succession. The robustness of pangenome methods to varying levels of sequence divergence is currently unknown. CY and CS were included as the outgroups in the PGGB graph and minigraph, respectively. Using alignments of over 400 genes orthologous between scrub-jays and primates, we found that the sequence divergence between AC and AW, and between AW and CY, was approximately half that between human and chimp and human and orangutan, respectively (fig. S20-21, table S7), despite similar divergence times among these species pairs (Kumar et al. 2022). The modest sequence divergence among *Aphelocoma* species reflects the expected slower substitution rate of birds compared to primates and other mammals (Green et al. 2014, Zhang et al. 2014).

Across all 90 haplotypes (including the CY outgroup) and the reference AW assembly, we input a total of 113,401,058,458 bp of assembly in 113,148 contigs into the PGGB pipeline (Fig. 2A). The main PGGB pangenome graph (Fig. 2A) incorporated 92,619,931,975 bp (81.7% of input) in 45190 contigs across 30 communities, each corresponding to an AC chromosome (Romero et al. 2024). Another 13,238,463,357 bp (11.7%) in 56,813 contigs were incorporated into 18 additional communities with AW reference sequences but without chromosomal assignment. An additional 7,319,110,588 bp (6.5%) in 6,856 contigs were incorporated into 946 additional communities without any reference assembly sequence. The node depth in a pangenome graph records the number of times a node is traversed by haplotypes in the graph (Guarracino et al. 2022). High depth nodes (in our case depth > 90 haplotypes) indicate repetitive regions that are succinctly described by haplotypes traversing a node repeatedly, whereas nodes close to depth 90 (or 45 in the case of sex chromosomes in females) indicate single- or -low-copy regions. All 90 haplotypes populated most regions in the 30 chromosome-level communities in the PGGB pangenome graph, whereas approximately 43% of chromosome 15, 37% of chromosome 2 and 11-14% of chromosomes 1, 1A, and 5, had regions of depth less than 10, suggesting inadvertent loss of these regions during graph construction (Fig. 2A). We note that several of these missing regions are flanked on one side by large inversions, which may have challenged the PGGB pipeline (Fig. 2A).

**Figure 2.**
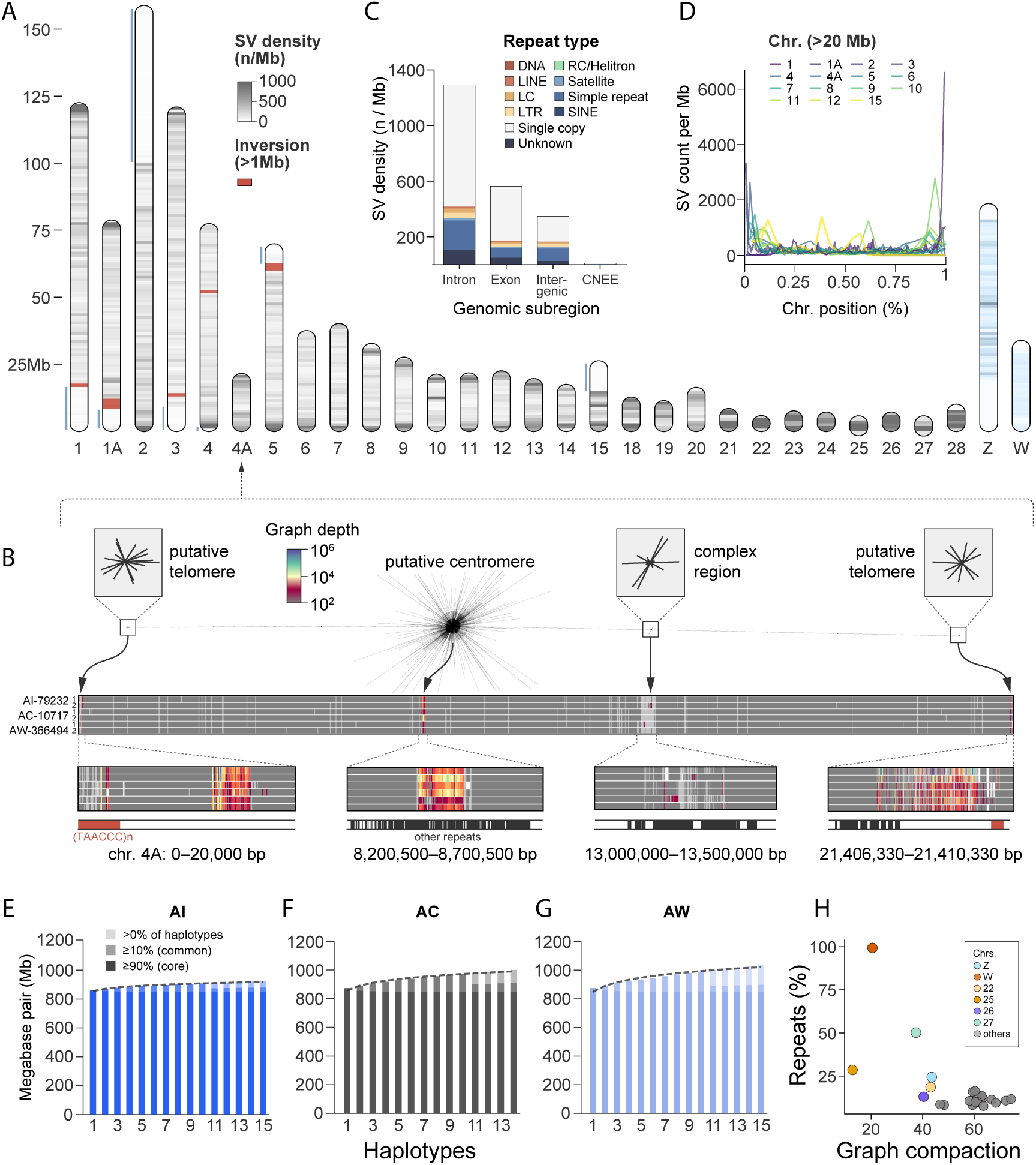
Pangenome graph captures genome-wide structural variants (SV). **(A)** Chromosome-level schematic indicating overall SV density (n/Mb) and large inversions (>1 Mb). Light blue vertical lines indicate regions that failed PGGB graph construction. **(B)** Pangenome graph depth on chromosome 4A, revealing high-depth blocks in putative telomeric regions, centromeric regions, and other complex regions. **(C)** SV density in different genomic compartments (x-axis) and overlap with major repeat categories (stacked bars). **(D)** SV counts per Mb along chromosomes. **(E–G)** PGGB pangenome growth curves show pangenome size (in Mb; y-axis) added by each sample (x-axis) to the graph for AI, AC, and AW, respectively. **(H)** Graph compaction (x-axis) versus percent repeats (y-axis) by chromosome, showing the highest compaction for chromosomes with lower repeat content.

We found that graphs of whole chromosomes exhibited a wide range of node depths that faithfully recorded major repetitive regions, telomeres and putative centromeres within chromosomes (Fig. 2B-D). Ultra-high depth regions (generally > 10,000 - > 100,000) tended to begin and end abruptly, producing block-like patterns sometimes exceeding 900 kb in length (fig. S22). High-depth regions (> 1000) of chromosomes, such as those tentatively assigned to centromeres, were generally dominated by single satellites or long-terminal repeats (fig. S23).

The highest depth regions of chromosomes 1,1A, and 3 were all dominated by sj_sat_circ146−2809, whereas such regions on other chromosomes tended to be dominated by chromosome-specific satellites. Two-dimensional graph visualizations of many of the main communities revealed putative centromeres and telomeric sequence (Fig. 2B).

To understand the dynamics of pangenomes in birds, we quantified ‘core’ sequence (present in >90% of haplotypes), versus ‘accessory’ sequence, (either haplotype-specific or found in <10 % of haplotypes), using the Panacus pipeline (Parmigiani et al. 2024) (Fig. 2E-G). The pangenome graphs of all three species implied a high percentage of the genomes as being core (AW, 79.4%; AC, 84.7%; AI, 89.3%). The accessory sequence was more variable among species: whereas 13.3% of the sequence in the AW graph occurred in only 1-2 individuals, only 4.6% of AI graph sequence was estimated to be accessory. Although some of this variation could result from variation in assembly length and quality between species, variation in the percent of accessory sequence between species is consistent with predictions from the relative amounts of diversity in the three species: AI birds on average share more sequence within species than AW and AC birds. The patterns in scrub-jays show striking similarities as well as differences from the few pangenome graphs available for other vertebrates. For example, at similar sampling intensities, the human pangenome exhibits proportionally much less core sequence (56.2%) than *Aphelocoma*, whereas a pangenome of inbred strains of chicken exhibited a percent accessory sequence (5.0%) similar to the AI pangenome (fig. S24). The lower core sequence in birds than in humans could be driven in part by technical differences because each of the haplotypes in the human pangenome was more accurate and highly resolved than the haplotypes studied here (Liao et al. 2023). However, analysis of a chicken pangenome (Rice et al. 2023) constructed with sequencing methods broadly similar to those used here, allows more confidence in the comparison. Graph compaction – the ratio of the length of the input sequences to the length of the graph (calculated as the sum of the lengths of all nodes) – was lowest for the highly repetitive W chromosome, whereas greater compaction was observed for chromosomes with less repetitive DNA (Fig. 2H).

### Structural variant diversity within and between species

To call SVs, we projected the PGGB pangenome graph to VCF format using vg deconstruct v1.40.0. The AW female reference assembly provided standardized coordinates for all haplotypes, the CY outgroup provided ancestral states, and we employed a reproducible protocol for classifying multiallelic and complex SVs (Materials). Approximately half (51.3%) of SVs were able to be polarized, but it is difficult to distinguish the underlying causes of polarization failure. Many SVs likely could not be polarized because of the smaller assembly size of CY haplotypes, but in principle some could not be polarized because of assembly gaps in the CY outgroup for particular *Aphelocoma* SVs or because of difficulties in incorporating the diverged outgroup into the graph structure. Throughout, we refer to insertions and deletions <50 bp in length as indels (INS and DEL) and those >50 bp as SVs (SV_INS and SV_DEL).

As found in genome-wide studies of other organisms, including birds (Nam et al. 2012, Barton and Zeng 2019), there is a marked bias towards deletions, most evident in the 1,916,338 indels in AW, where the DEL/INS ratio is 1.50, compared to 1.39 (n=838,106) and 1.29 (n=201,649) in AC and AI, respectively (fig. S25, table S8). The bias is less pronounced among SVs: the ratio of SV_DEL to SV_INS is 1.20 in AW (n=158,686), 1.22 in AC (n=72,896) and 1.31 in AI (n=35,442). SV indels recorded in the PGGB pangenome graph ranged in size from 50 - 91,520 bp (table S8). Across all species, the mean (1,125 bp) and median (172 bp) length of SV_INS were generally greater than SV_DEL (mean 329 bp, median 148 bp), possibly reflecting the impact of transposable elements, which overlap with 22% of SVs, similar to the percentage overlapping simple sequence repeats (21%; Fig. 2C). The number of SVs identified by other pangenome (minigraph) and reference-based (svim-asm) methods are generally fewer than those identified with PGGB, but exhibit similar trends within and across species (table S9).

There was a marked increase in SV density towards chromosome ends, compared to chromosome interiors (Fig. 2D). The density of SNPs, indels and SVs each exhibit positive correlations with recombination rate in highly repeatable decreasing rank order (SNPs: R^2^=0.22; indels: R^2^=0.12; SVs: R^2^ = 0.03) (fig. S26); as a consequence, all three types of variants exhibited the highest density on microchromosomes (fig. S27). SVs were rarest in highly conserved non-exonic regions (CNEEs), which are known to influence gene regulation and act as enhancers in birds (Sackton et al. 2019) (Fig. 2B, fig. S25). Surprisingly, we found that the density of SVs (SVs per Mb) was highest not in intergenic regions, which comprise 59% of the reference genome, but in introns, which comprise only 12.7% (Fig. 2C). Also surprising is the finding that, although the number of SVs in exons is much lower than in noncoding regions, the density per Mb is higher than even intergenic regions (521 per Mb in exons versus 328 per Mb in intergenic regions; Fig. 2C, fig. S25, table S8). The high proportion of SVs in exons and introns may signal a relationship between SV formation and germline transcription, which would disproportionately affect exons and introns.

In the PGGB graph, we detected a total of 448,012 SVs across all three *Aphelocoma* species. The rank order of the number of variants for SNPs, indels and SVs in the PGGB graph is AW > AC > AI (Fig. 3A,B). Biallelic SNPs were ∼8.5 times as common as biallelic indels, which in turn were about 20 times more common than biallelic SVs (Fig. 3C). By contrast, multiallelic SVs were ∼1.6 times as common as multiallelic SNPs, even though the average number of multiallelic SV alleles (4.18) was on par with the maximum for SNPs (Fig. 3A,B).

**Figure 3.**
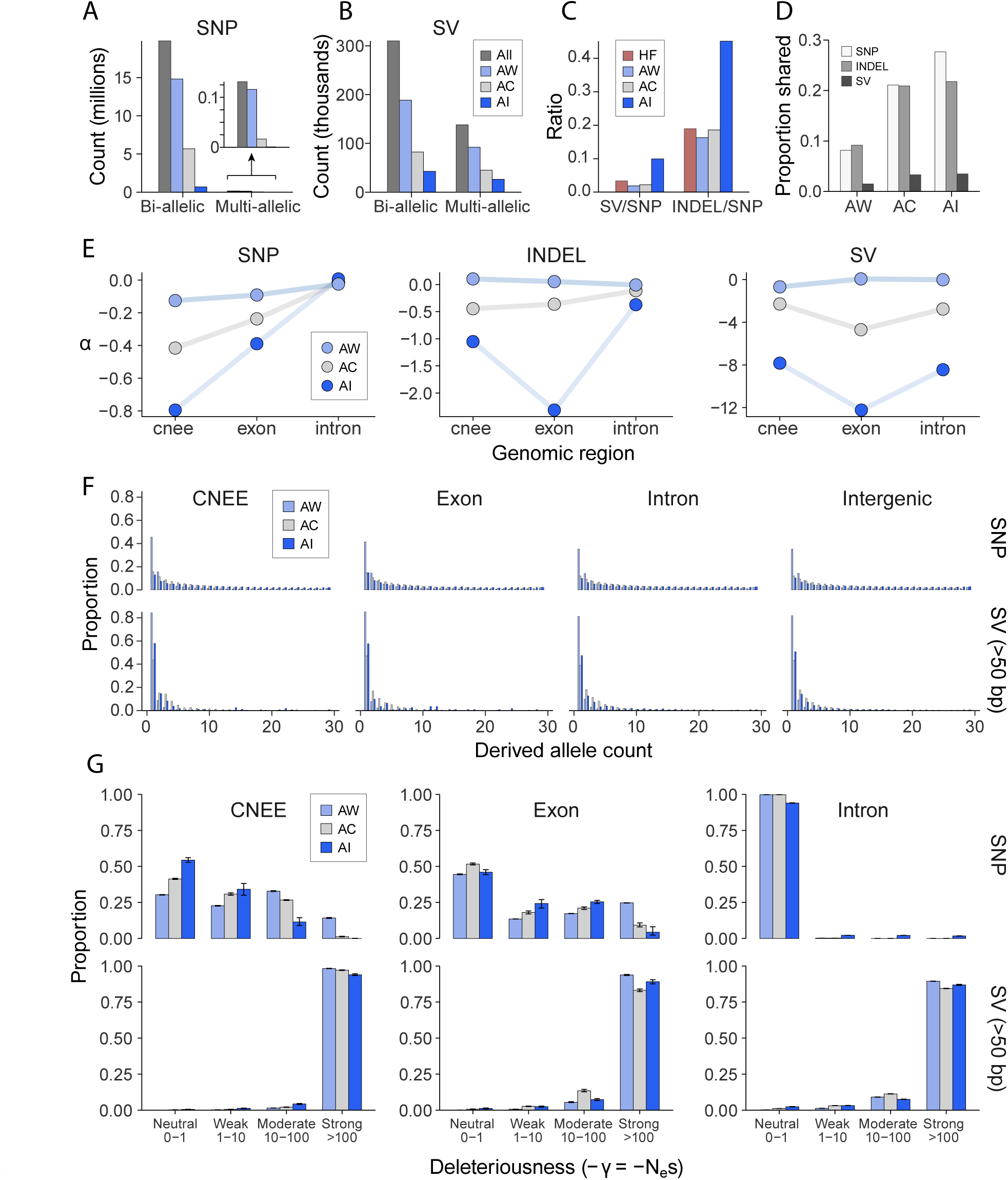
Comparative analysis of SNPs, indels, and structural variants. **(A,B)** Counts of biallelic and multi-allelic SNPs (**A**) and SVs (**B**) across species. Inset in (**A**) shows expanded scale for multi-allelic SNPs. **(C)** Ratios of indels/SNPs and SVs/SNPs for the three scrub-jays and House Finch (HF). **(D)** Proportion of variants shared among species. **(E)** Estimates of the fraction of variants fixed by selection (α) for SNPs, indels, and SVs across three genomic compartments: conserved non-exonic elements (CNEEs), exons, and introns. **(F)** Derived-allele frequency spectra in CNEEs, exons, introns, and intergenic regions for each species. **(G)** DFE in bins of population-scaled selection coefficient (γ = Nes) estimated by fastDFE, reflecting variant deleteriousness, a function of the effective population size (*N*e) and the selection coefficient (*s*). DFE was inferred from SFS of different variant types residing in different genomic compartments.

Interspecific variation in the ratio of SNPs to indels and SVs within each species can provide initial insight into the selective forces acting on SVs and indels and is robust to biases resulting from reference effects. The ratio of SVs and indels to SNPs varied from 0.019/0.163 in AW, to 0.022/0.186 in AC and up to 0.099/0.451 in AI (Fig. 3C), consistent with the hypothesis that indels and SVs are more deleterious than SNPs and more easily purged in AW and AC versus AI, assuming neutrality of SNPs. These patterns hold when considering variants only in intergenic regions, which are more likely to meet the assumption of neutrality for SNPs (AW: 0.016/0.147; AC:0.019/0.169; AI:0.087/0.369). Strikingly, in House Finches (*Haemorhous mexicanus*; HF), which have recently been surveyed for SVs using methods similar to those used here (Fang and Edwards 2024), the ratio of SVs or indels to SNPs is similar to that found in AW and AC, despite the number of SNPs and SVs being nearly twice as high in HF as in AW or AC, and highlighting the extent to which AI is an outlier. The number of SVs shared between species and species-specific SVs also reflect these trends (Fig. 3D, fig. S28). Biallelic SNPs are 5.5 times more likely to be interspecifically shared by incomplete lineage sorting than SVs (6.8 % of SNPs versus 1.2% of SVs), whereas indels and SNPs have similar proportions shared between species (7.45% indels shared), patterns that hold for individual genomic compartments (fig. S29).

### Distribution of fitness effects of structural variants

To estimate the distribution of fitness effects (DFE) of indels and SVs, we conducted a variety of tests. Classical tests of neutrality based on summary statistics that have been couched in terms of the numbers of synonymous and nonsynonymous polymorphisms and fixed differences in protein-coding genes (Smith et al. 2002, Stoletzki et al. 2011) also offer a useful logic for the analysis of SVs (Barton et al. 2018). We first adapted an estimate of *α*, the fraction of variants fixed between species by positive selection, by equating fixed and polymorphic intergenic SNPs to counts of synonymous variants, and SVs overlapping different genomic compartments (CNEEs, exons, introns and intergenic) to nonsynonymous variants (Stoletzki and Eyre-Walker 2011), taking care to filter out variants below a frequency of 15% (derived allele count of ∼4), so as not to dilute the estimate with slightly deleterious variants segregating at low frequency (Gossmann et al. 2010, Stoletzki and Eyre-Walker 2011). For categories other than intergenic SNPs, which here are used as a neutral standard, we found that *α* was overwhelmingly negative across species and genomic regions, most consistently for indels and SVs in AI birds, which were uniformly negative or 0 *α* for all genomic categories (Fig. 3E, table S10). However, a small fraction (1-6%) of insertions in intergenic regions and introns in AC birds, and between 32% and 40% of insertions, SV insertions and SV deletions across all genomic compartments in AW birds, were estimated to be fixed adaptively (Fig. 3E, fig. S30, table S10), as in the Great Tit (*Parus major*), another species with a large effective population size (Barton and Zeng 2019). Estimates of the direction of selection, which is less biased by small sample sizes (Stoletzki and Eyre-Walker 2011), reflect trends similar to *α* (table S10).

To estimate population genetic parameters of the DFE, specifically the distribution of the scaled selection coefficients of new mutations (*γ*, the product of the effective population size and selection coefficient), we used two maximum likelihood methods (Barton and Zeng 2018, Sendrowski et al. 2024), both of which incorporate the possibility of misidentification of ancestral states, as well as variation in mutation rate among site classes, both of which can compromise estimates of the DFE if ignored (Kvikstad et al. 2014, Glémin et al. 2015, Tataru et al. 2017). These methods use the site frequency spectrum of SVs and indels to estimate the DFE by comparing their SFS to those of putatively neutral SNPs, in our case intergenic SNPs (Fig. 3F). Given our generally small estimates of *α*, we constrained our ML estimates of *γ* to be 0 or negative, allowing us to assume more realistically variable mutation rates among SNP controls and SV sites. Across all genomic compartments, we found that estimates of *γ*for SVs tended to be more negative, and hence more deleterious, than for indels or SNPs (Fig. 3G; fig. S31). Estimates of the distribution of *γ* were broadly similar across species and methods, implying that the selection coefficients of variants segregating in AI birds are likely more negative than for AW and AC birds, given the smaller *N*_e_ for AI birds (Fig. 3G, fig. S31).

### Selective consequences of SV length

The length of indels and SVs could influence their selective effects. We found that the mean length of indels was longest in AI (4.94 bp) versus AW and AC (4.14 and 4.11 bp, respectively; figs. S32-S34; table S11), a trend that also held for bi- and multiallelic SVs but not for complex SVs (table S11). To explore the relationships between SV length and fitness consequences, we first examined the relationship between SV length and derived allele frequency across the three focal species. The average derived allele frequency of a class of variants is a measure of its likelihood of being neutral or adaptive, whereas variants only found at low frequency are more likely to be deleterious. We found that the number of large (> 500 bp) SVs reaching high frequency in AI birds was greater than in AW or AC birds (Fig. 4A-C, fig. S35), consistent with drift driving frequencies of large and potentially deleterious SVs upwards. Using fastDFE (Sendrowski and Bataillon 2024), we also found a detectable and predictable increase in estimated deleteriousness with length in base pairs of indels and SVs (fig. S32). Only the indel length class of 1-5 bp had an appreciable proportion of sites with *γ* = 0. As the length of indels increased to ∼ 30 – 40 bp, the estimates of *γ* increased accordingly, until they leveled out above this length. As a result, estimated selection coefficients of indels (< 50 bp) varied strongly with length, whereas SVs as a group yielded estimates of *γ* that were less influenced by length. These patterns held broadly across CNEEs, exons and introns, and are also evident in length differences for SVs in heterozygotes and homozygotes, especially with SVs > 1 kb (fig. S36).

**Figure 4.**
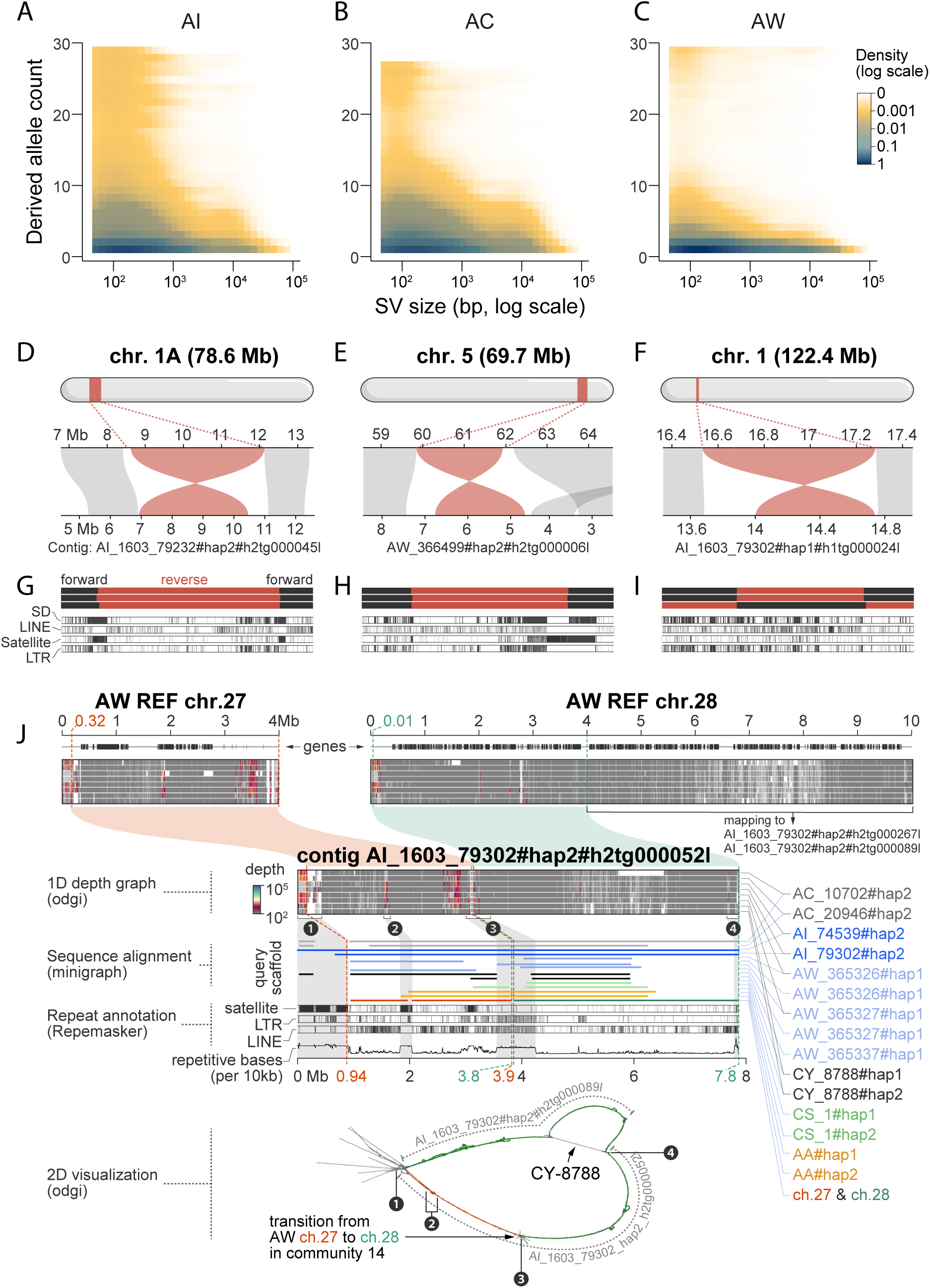
Genomic complexity revealed by pangenome tools. **(A–C)** Two-dimensional density plots (log scale) showing derived allele frequency (y-axis) versus SV length (x-axis) for three Scrub-Jays. **(D–F)** The largest three large inversions detected on chrs. 1A, 5, and 1, respectively, with their approximate positions (Mb) of inversion breakpoints. **(G–I)** Close-up views of flips in strand across the inversion (visualized with *odgi viz* on PGGB graph) and repeats (segmental duplications [SDs], satellite DNA, long interspersed nuclear elements [LINEs], and long terminal repeats [LTRs]) near putative inversion breakpoint regions. **(J)** A chromosomal fission detected in AW on chrs. 27 and 28. Upper panels show a 1D view of pangenome graph depth, repeat annotation, and gene tracks; lower panel (“bird” shape) is a 2D visualization of the PGGB graph. Each line indicates single contigs for AI, AC, AW and CY outgroup, confirming lack of AW contigs spanning the two reference chromosomes and mapping at least 250 kb to the same strand of the long AI haplotype spanning AW chrs. 27 and 28 (contig AI_1603_79302#hap2#h2tg000052l).

### Inversions and chromosome evolution

Inversions represent an important class of SVs and have been implicated in several polymorphic phenotypes and adaptive traits in birds (Lamichhaney et al. 2015, Funk et al. 2021, Knief et al. 2024, Loveland et al. 2025). To quantify the landscape of inversions in *Aphelocoma*, we used four pangenome- and reference-based methods with different sensitivities and abilities to detect inversions of different lengths: PGGB (Eizenga et al. 2021), minigraph2 (Li et al. 2020), SyRi (Goel et al. 2019) and svim-asm (Heller et al. 2021). All of these methods are best at detecting small to moderately-sized inversions, so we supplemented these approaches with genome-scanning methods, such as sliding window PCAs and multidimensional scaling (MDS), which are better at detecting large to very large inversions (Li et al. 2019, Harringmeyer et al. 2022). Across four methods employing mapping or pangenome graphs, we found a total of 382 inversions varying in length from 50 bp to 3.7 Mb (fig. S37). Ninety-five of these inversions were detectable by at least three of the four detection methods (Fig. 4D-I, fig. S37). Diverse repeats were enriched in regions flanking inversions (fig. S38) and in many cases inversions were shared among species, either ancestrally or arising recurrently within species (fig. S39). Overall, 249 genes occurred inside the 95 high-confidence inversions, but no functional enrichment of this genic subset was evident. Given our sample sizes within species, our power to detect larger inversions via genome scanning approaches is limited. Still, we discovered four larger inversions (> 1 Mb: 3.5 Mb - chr4A in AC, and 3.2 Mb - chr5 and 1.5 Mb - chr10 and 1.2 Mb – chr15 in AW) that are confirmed by multiple lines of evidence, including MDS, 1D pangenome graph strand visualizations and PCA (figs. S40-S42). When we include these larger inversions, one GO term, “kinesin complex”, is significantly enriched by 7 relevant genes. Inversion lengths did not exhibit a clear difference when in the homozygous versus heterozygous state (fig. S43). Nor within each species was there a clear relationship between inversion length and derived allele frequency, except in AI birds (fig. S44).

Minigraph, in combination with the other tools (Materials), revealed unexpected complexity in the form of a chromosomal fission in AW haplotypes (Fig. 4J, table S12). A ∼1 Mb segment of multiple haplotypes in AI, AC, AA, CS and CY, but no AW haplotypes, mapped to both AW reference chromosomes 27 (∼3.93 Mb) and 28 (∼9.94 Mb). This fission event harbors 135 genes and is flanked by highly complex repeat and satellite landscapes and higher pangenome graph depth (Fig. 4J). Using similar criteria, the PGGB pangenome graph and associated visualizations confirmed 30 of 34 AI and AC haplotypes spanning the chromosomal fission identified by minigraph, as well as two additional haplotypes each in AC and CY birds (fig. S45). Our ability to genotype all birds for this putative fission event is limited to those haplotypes possessing single contigs spanning both AW reference chromosomes. We therefore provisionally suggest that the fused condition is ancestral in CY and CS and that the fission is derived exclusively within AW haplotypes.

### Copy number variants of genes and gene expression

An important component of the SV landscape are copy number variants (CNVs) of genes. We used miniprot (Li 2023) and pangene (Li et al. 2024) to detect CNVs, estimate gene copy numbers per haplotype and construct genome-wide pangene gene graphs for 96 haplotypes and 14,112 well-annotated genes (13,515 autosomal) across ingroups and outgroups (Fig. 5A-C). Pangene aligns amino acid sequences to DNA contigs in the genome and requires well-annotated and highly complete haplotype assemblies. Moderately fragmented assemblies such as ours, as well as the difficulty of aligning non-contiguous protein sequences to DNA, increase the likelihood of false negatives, whereas false positives are less likely. Still, the overall trends in the dynamics of CNVs within and between species were robust to various filtering strategies. We found hundreds of cases of gene copy number variation (Fig. 5A-C, table S13, S14). Surprisingly, and in contrast to the distribution of SVs within species, AI birds harbored the most genes experiencing CNVs, over twice as many as found in AC and AW across filtering strategies (Fig. 5D,E). For biallelic CNVs found in at least two haplotypes, we detected 2905 in AI, 1170 in AC and 892 in AW. Approximately 83.9% of CNVs in AI were AI-specific, whereas this number was much lower in AC (37.3%) and AW (19.5%) (table S14). Across species, AI birds experienced the greatest number of haplotypes homozygous for inferred deletion or truncation of genes, about 30-70% more than AC and AW (Fig. 5D,F, table S14). We have no reason to suspect that counts of deleted genes in AI birds should be biased upward, conceivably, for example, due to fragmented genome assemblies. AI birds had among the highest quality de novo assemblies in our sample and moreover were low in heterozygosity, both of which should aid detection of genes by pangene; if assembly quality influenced counts of gene deletions, we would expect AW birds to have the highest number of gene deletions because their assembly quality was lowest. Across loci within species, deletions of genes were between 2 and 80 times more common than multiplications of genes (Fig. 5F, G). In contrast to gene deletions, gene multiplications were less common in AI (18 genes) than in AC (157) and AW (58) (Fig. 5G, fig. S46). Four genes (*FAM179A, GSTA3, MEPE, RDH16*), all characterized by multiplications, exhibited significant differences in copy number between species after correction for multiple tests (fig. S47). *FAM179A* varied from a mean of 3.9 in AI (95% c. i.) [3.3 – 4.6] to 10.5 in AC [9.5 – 11.6]; copy numbers on the two haplotypes of outgroups CY (10,15) and CS (17,18) were even higher. Nearly all autosomal CNVs were in Hardy-Weinberg equilibrium; as expected if CNVs were deleterious, the average frequency of homozygous deletions across genes within species was low, never exceeding 21% and most cases, depending on filtering strategy, never exceeding 14% (table S14).

**Figure 5.**
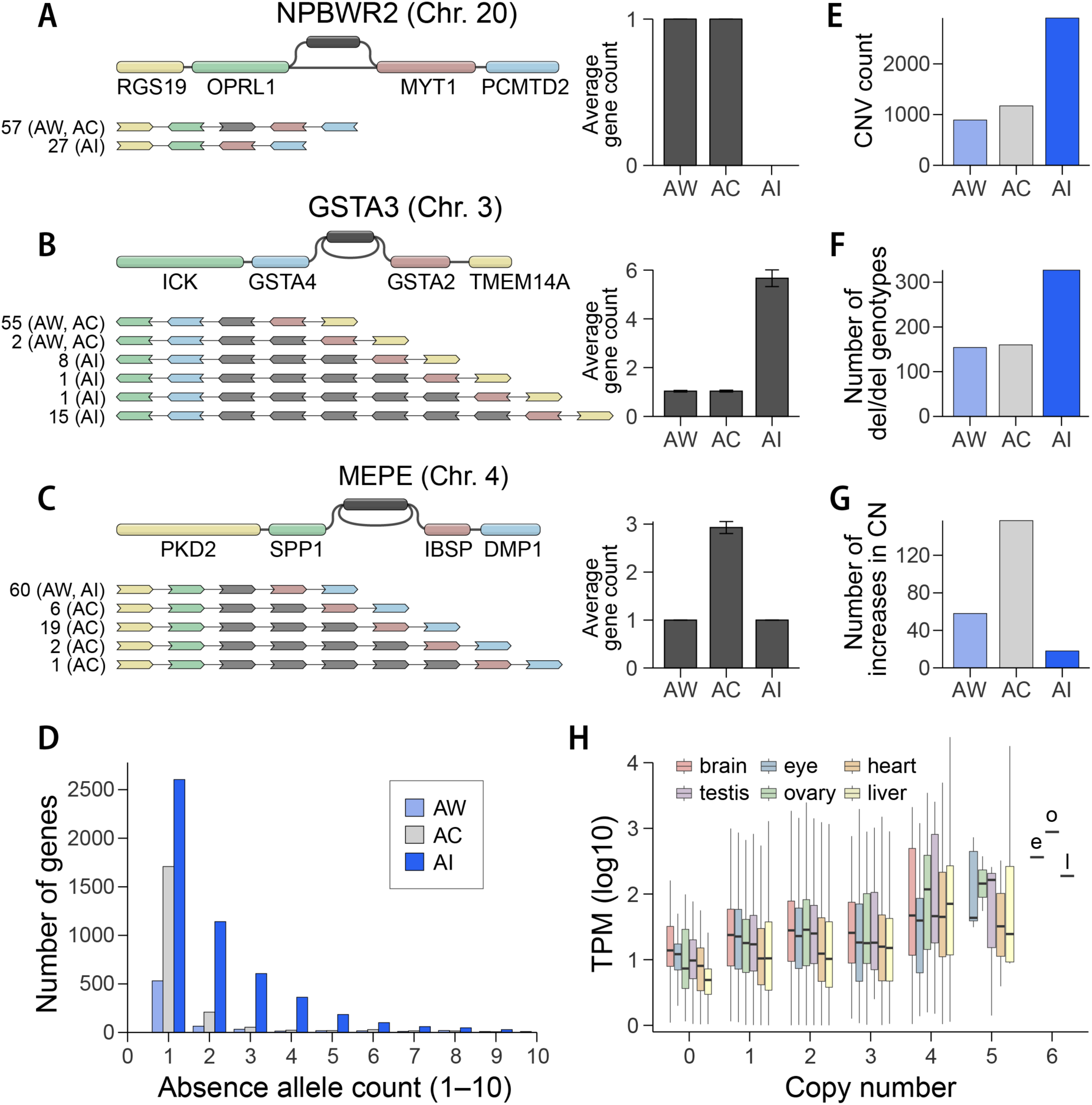
Copy number variation and gene expression. **(A–C)** Three examples of gene presence–absence variation (PAV; **A**) and copy number variation (CNV; **B**, **C**) within the pangenome gene graph constructed using Pangene, with bar plots of average copy number across species (right). **(D)** Number of genes (y-axis) for which a given “absence allele count” (x-axis) was detected per species. **(E)** Total number of CNVs per species. **(F)** Number of genes with homozygous deletions per species. **(G)** Genes with copy number increases (multiplications) for each species. **(H)** Log10-transformed transcripts per million (TPM) values in AW across tissues (brain, eye, heart, testes, gonad, liver) by gene copy number, indicating that elevated expression is associated with higher gene copy number (p = 0.048).

To detect the effect of gene copy number on gene expression, we sequenced transcriptomes of 5 tissues across each of the 15 AW birds (brain, eye, liver, heart, and gonad) and blood for each of the 14 AC birds, for a total of 62 transcriptomes. We quantified gene expression across 15,870 genes, of which 10,671 both had expression levels greater than 1 transcript per million (TPM) in at least one tissue in AW birds and were included in the pangene analysis (fewer genes – 7,581 – had TPM >1 in AC birds, as expected given our use of whole blood). The number of genes for which we detected expression (TPM > 1) was highest for gonads (14,581 genes) and lowest for liver (11,715 genes), with similar trends in average TPM. Our study design and sample sizes did not allow us to test for an effect of copy number on TPM for individual genes, or to test for associations between SVs and gene expression. However, a linear mixed model fit to autosomal genes with TPM as the response variable, copy number as the independent variable and gene and tissue as random effects, revealed a significant effect of AW gene copy number on TPM within and across tissues (p = 0.048; Fig. 5H, (Materials)).

## Discussion

Comparative population genomics is an emerging field striving to understand the determinants of genomic diversity among species, but thus far has focused primarily on SNP variation (Ellegren and Galtier 2016, Edwards et al. 2021). Studies of polymorphisms of indels and SVs are growing in number, but rarely has it been possible to compare multiple species across the full spectrum of SNPs, indels and structural variants, including inversions. Here we used long-read sequencing at the population level among multiple species as well as novel pangenome computational tools to understand the dynamics of structural variation among three species of birds with drastically different effective population sizes. To our knowledge, the only other similarly sized pangenome study is the human pangenome, currently consisting of 45 high-quality long-read assemblies (Liao et al. 2023). However, the assemblies comprising the human pangenome studies, although of even higher quality than those presented here, are thus far from a single species, and one of relatively low genetic diversity (Fig. 1D). Our pangenome graphs comprise three species roughly half as diverged as human to chimp and gorilla, and include (in the PGGB graph) an outgroup about half as divergent as orangutan is from human. The complexity of the human and avian graphs is necessarily driven by substantial differences in genome size (Liao et al. 2023): the birds studied here have genomes roughly 42% that of the human genome.

Our survey of structural variation in *Aphelocoma* jays revealed a pervasive influence of effective population size on variation in diversity and dynamics of structural variation between species. Estimates of the fraction of SVs driven to fixation by natural selection were generally low or zero in AI and AC, suggesting that fixations occur primarily by drift in these species; AW was the only species for which estimates suggested a substantial fraction (∼30-40%) of SVs appear to have been fixed by selection. The distribution of inversions, which thus far have been studied in small numbers per species and biased with an eye towards demonstrating adaptive significance, uniformly suggests that they tend to be deleterious and that adaptive inversions might be rare exceptions to this overall trend (Fang and Edwards 2024, Loveland et al. 2025). Ours and other long-read genome assemblies of birds yielding exceptionally high fractions of repetitive DNA (Manthey et al. 2018, Benham et al. 2024) have begun to revise our understanding of the prevalence of repetitive DNA in avian genomes. Our combination of approaches suggests a much richer satellite landscape and has the advantage of uniform lab and bioinformatic approaches applied to each species. Future research should emphasize standardization and benchmarking of approaches aimed at generating consistent curation and databasing of avian satellites.

The dramatic differences in assembly size among the three species studied here was unexpected. Rapid shifts in genome size are known for diverse lineages of animals and plants (Naville et al. 2019, Blommaert 2020, Adams et al. 2023) and are most often attributed to the proliferation or deletion of transposable elements. Population genetic theory suggests that species experiencing long-term reductions in population size should experience increases in genome size due to the accumulation of deleterious mutations, including transposable elements (Lynch 2007, Lynch et al. 2011); here, unexpectedly, the relatively recent bottleneck experienced by AI birds during or after the founding of Santa Cruz Island is associated with a decrease in assembly size. The decreases in abundance of some satellites and telomeres in AI relative to AW and AC suggests that, in the short term, bottleneck-associated decreases of some types of repeats may be expected. The pangene analyses also revealed an unexpected dynamism in gene copy number evolution, suggesting both the effects of drift and potential substrates for adaptive evolution, as well as influence on gene expression. The patterns of CNV within and among species suggest that gene deletions are generally deleterious and are fixed at a higher rate in species with small *N*_e_, but are selectively eliminated more readily in species with large population sizes (Lynch et al. 2000). However, the number of genes exhibiting duplicated or mutiplicated variants was highest in AW birds, implying, by contrast, that gene multiplications are on average potentially neutral or adaptive, surviving as polymorphisms longer in species with large *N*_e_.

A major challenge with the analysis of SVs is the determination of ancestral states, especially in cases of multi-allelic and complex SVs, as well as their representation in VCF files (Barton and Zeng 2019). The challenges of incorporating divergent outgroups into pangenome graphs points to a key methodological frontier: the need for tools that can simultaneously handle both recent and ancient divergence in a single graph structure. A promising future approach involves leveraging the implicit pangenome graph model - partitioning alignments based on their implied graph structure, constructing separate graphs for each partition, and then reconnecting these partitions into a comprehensive graph. Newer machine-learning approaches to SV characterization (Popic et al. 2023, Zheng et al. 2023), standardization and transparency of SV calling pipelines (as with SNPs (Mirchandani et al. 2024), as well as greater incorporation of phylogenetic perspectives into bioinformatic summaries of SVs and other variants (Digiacomo et al. 2022) may improve reliability, repeatability and comparability among studies.

## Acknowledgements

We thank Gregg and Donna Schmitt, Flavia T. Garcia, Kathrin Näpflin, Gunnar Kramer, and Subir Shakya for assistance with fieldwork in New Mexico and the Yucatán and Tori Bakley, Reed Bowman, and other Archbold staff for collection of Florida samples. We thank O. Shevchenko, E. Bernberg, and B. Kingham from the University of Delaware Sequencing & Genotyping Center for their assistance with PacBio sequencing. J. Trimble and K. Eldridge of the MCZ Department of Ornithology facilitated access to specimens. N. Mejia isolated the RNA samples. M. Houck and A. Misuraca performed karyotype analysis. M. Hiller provided the human-chicken TOGA alignment and E. Osipova produced the primate and jay alignments and the TOGA flowchart. A. Suh and V. Peona provided access to corvid satellite sequences. We thank C. Ané, G. David and J. Höglund for helpful discussion, and K. Lopez and T. Pegan for help with recombination and inversion analyses, respectively.

## Funding

SVE acknowledges support from Harvard University and from National Institutes of Health (NIH) grant R01HG011485; E.G. acknowledges support from NIH/NIDA U01DA047638, NIH/NIGMS R01GM123489, and NSF PPoSS Award 2118709. NC acknowledges NIH grant 1R35GM133412.

## Author Contributions

SVE conceived of the project, provided funding, conducted fieldwork, organized samples for sequencing, analyzed data and wrote the manuscript. BF conducted fieldwork, analyzed data, assisted with project design and writing and produced all main figures and many supplemental figures. TBS, DK, DAD analyzed data, assisted with project design and with writing. GEK analyzed data. RGC and NC provided samples and assisted with writing. HL and AG analyzed data, provided project direction, and edited the paper. JEM, WCF, CKG, EG and JWF assisted with interpretation, project design and editing the manuscript.

## Competing interests

## Data and materials availability

Sequence data for this work has been deposited in NCBI under Umbrella BioProject PRJNA1206191, the Scrub-Jay (*Aphelocoma*) Pangenome Project. Within this project, BioProject PRJNA1204306 contains the PacBio HiFi reads (with samples SAMN46016487 - SAMN46016457), whereas BioProjects PRJNA1204814 - PRJNA1204903 contain the haplotype assemblies. All scripts can be found at https://github.com/harvardinformatics/scrub-jay-genomics. Additional resources, including files necessary to set up the pangene results on a web server, are available to reviewers on Dryad.

## Supplementary Materials

### Materials and Methods

### Field methods

#### Field Methods for Island Scrub-Jays (AI, *Aphelocoma insularis*)

The island scrub-jay is restricted to Santa Cruz Island, which is 32 km from the mainland of southern California, USA. The census population size is estimated to be between 1200-3000 individuals (Sillett et al. 2012), while the effective population size (Ne) is estimated to be 346.8 with a 95% confidence interval of 327–368 based on a few thousand SNP loci (Cheek et al. 2022). The 250 km2 island has a Mediterranean climate characterized by cool, wet winters and hot dry summers. The vegetation is composed of a mosaic of habitat dominated by island scrub oak and includes coastal sage scrub, oak woodlands, and oak chaparral (Junak 1995, Fischer et al. 2009). Juvenile and adult male and female island scrub-jays were captured using either mist nets or box traps from three large study plots from 2009 to 2011 (see Caldwell (2013) for detailed descriptions of the study plots). Morphological measurements were collected from each captured individual using digital calipers to record (to ±0.01 mm): bill length measured from the anterior end of the nares to the tip of the bill; bill depth, measured at the anterior end of the nares; and tarsus length. Wing chord and tail length were measured with a ruler (to ±0.5 mm). We marked jays with a unique combination of up to 5 colored plastic leg bands and 1 numbered U.S. Geological Survey band. Whole blood samples were extracted from the brachial vein and preserved in Queen’s lysis buffer (Seutin et al. 1991). All work with living birds was approved by the Institutional Animal Care and Use Committees at Colorado State University (IACUC: #887) and the Smithsonian Institution.

Neutral population genetic analyses have shown that island scrub-jays exhibit isolation-by-distance across the east-west axis of the island which is likely driven by limited dispersal (Langin et al. 2015, Cheek et al. 2022). Our criteria for selecting individuals for sequencing were that they be unrelated, have multiple blood samples taken between 2009 and 2011, and show limited spatial genetic structure. Based on these criteria, we selected eleven male and four female island scrub-jays for PacBio HiFi sequencing from our two study plots located in the central valley of Santa Cruz Island. We confirmed that all individuals were less than full siblings based on a kinship matrix calculated using the Genome-wide Efficient Mixed Model Association software toolkit (Zhou et al. 2012, Cheek et al. 2022).

#### Field methods for Woodhouse’s Scrub Jays (AW, *Aphelocoma woodhouseii*)

Woodhouse’s Scrub-Jay specimens were collected under permit 3704 from the New Mexico Department of Game and Fish and Federal US Fish & Wildlife Permit Number MB155188-0. Field collection took place in October 2018 and June 2021 northwestern New Mexico in the vicinity of Turley, Navajo Dam City, Cedar Hill and Blanco. Specimens for karyotyping were collected in May 2023. Specimens were collected by mist-net and shotgun. Birds were targeted for collection randomly upon encounter in the field. After sacrifice, tissues were immersed and minced in RNAlater in individual nunc tubes and stored at room temperature for 24 hours, after which they were immersed in liquid nitrogen or vapor for transport back to the Museum of Comparative Zoology (MCZ) at Harvard University. The *Cyanocorax yucatanicus* sample from Yucatán, Mexico, was processed similarly to the AW samples. All specimens were prepared as vouchers and are retained in the MCZ. Full details on each specimen can be found in Table S1 and at MCZbase (https://mczbase.mcz.harvard.edu/).

#### Field methods for Florida Scrub-Jays (AC, *Aphelocoma coerulescens*)

The Florida Scrub-Jay is restricted to the xeric oak scrub habitats of Florida (Woolfenden and Fitzpatrick 1984). A population of individually-banded Florida Scrub-Jays has been monitored at Archbold Biological Station in Venus, FL since 1969, providing detailed information on individual life histories (Woolfenden and Fitzpatrick 1984). Florida Scrub-Jays have very low rates of extra-pair paternity (Quinn et al. 1999, Townsend et al. 2011) allowing us to reconstruct fairly accurate pedigrees from field observations of breeding behavior, though we also confirmed pedigree relationships with genetic data for >15 years (Chen et al. 2016). For this project, we sampled mostly younger individuals (< 3 years of age) out of convenience. We sampled birds with no known close relationships to each other (no parent-offspring, full sibling, or half sibling relationships). To minimize impacts on our study population, we focused on individuals who also needed a band replaced or were easily trappable. We captured individuals using Potter traps and collected whole blood from the brachial vein. Blood samples were stored in RNAlater at room temperature for 24 hours before transferring to a −80 freezer and shipped on dry ice. All work was approved by the Institutional Animal Care and Use Committees at Cornell University (IACUC 2010-0015) and Archbold Biological Station (AUP-006-R), with permits from the US Fish and Wildlife Service (TE824723-8, TE-117769), the US Geological Survey (banding permits 07732, 23098), and the Florida Fish and Wildlife Conservation Commission (LSSC-10-00205).

### Laboratory Methods

#### DNA isolation and PacBio HiFi sequencing

All samples were sequenced at the Delaware Biotechnology Institute DNA Sequencing and Genotyping Center in Newark, Delaware. For AI or AC blood samples, 100ul of sample was pelleted and the pellet was processed with Qiagen’s MagAttract HMW DNA extraction kit. Genomic DNA was isolated from the pellets using Qiagen’s MagAttract HMW DNA extraction kit. Lysis was shortened to 30 minutes at 56C° and all shaking steps were replaced with rotation. Elution was performed overnight at room temperature. DNA isolation from the AW tissue samples was performed on cryo-pulverized material following the Qiagen MagAttract kit instructions. Lysis was shortened as much as possible until there was no visible tissue remaining (typically 30-60 minutes) and was conducted at 56C°. All shaking steps were converted to rotation and samples were eluted overnight at room temperature. Genmic DNA quality was assessed with a Femto instrument (Agilent). All samples were sheared using a Megaruptor 3 (Diagenode) instrument with target size 14-15 kb. All libraries were prepared using the PacBio SMRT Bell prep kit v3 (Pacbio). Sample libraries were size-selected for 6-8 kb fragments using a Blue Pippin instrument (Sage). Each sample was sequenced on 2 flow cells of a Sequel IIe instrument using sequencing kit v2 and Sequel II binding kit 2.2 and 30 hours of movie was recorded. PacBio HiFi reads were downloaded from the Delaware Center web site as BAM files for further processing.

#### HiC sequencing

HiC sequencing of the AW reference individual (number MCZ Orn 365336, female) and AW male individual MCZ Orn 365338 was performed by Arima Genomics in San Diego, CA. 359,457,888 read pairs were generated on an Illumina MiniSeq sequencer from AW 365336 and 352,068,517 were generated from MCZ 365338. Approximately 250,000 reads were generated from human control samples to evaluate quality of the library, which suggested 58.4-59.5% long-range cis interactions. Paired-end sequences were downloaded as fastq.gz files for genome assembly.

#### Karyotyping

Ten *A. woodhouseii* were collected in northwestern New Mexico in May 2023 and whole eyes and tracheal tissue were extracted with sterile dissecting equipment by immersing them in fresh medium (alpha MEM + 10% FBS, + 1% Glutamine/Pen-strep, + 1% fungizone) in biopsy vials. Tissues were shipped at room temperature overnight, or in some cases after 24 hours in at 4° C in a refrigerator, to the Frozen Zoo, Reproductive Sciences and Conservation Science, San Diego Zoo for processing. All samples were analyzed with standard (non-banded) Giemsa staining. C- and G-banding was applied to one individual (female Lab#23907) sample. An example karyotype of is presented in figure S1, and was found to have a diploid number of 80. Full details of the karyotype analysis will be published elsewhere.

#### RNA isolation and sequencing

RNA was isolated from approximately 12 μl whole blood from *A. coerulescens* samples using Qiagen kits and the same methods as in (Mejia et al. 2024). Cells were separated from RNAlater in the tube by centrifugation. (20,800 × g). Cells were homogenized with Qiazol and zirconia/silica one mm beads (BioSpec Products) on a TissueLyser LT (Qiagen, Hilden, Germany) in the Harvard Bauer Core Facility. RNA was isolated from homogenized cells using the RNeasy Plus Universal mini kit protocol (Qiagen, Hilden, Germany). Libraries were prepared with KAPA mRNA Hyperprep kits by staff in the Bauer Core and sequenced on an Illumina NOVASeq SP platform with paired end reads of 150 bp length, yielding between 20 and 30 million reads per sample.

For tissues from *A. woodhouseii*, RNA was isolated from sample volumes of approximately 2-3 mm^2^. Isolation followed instructions in the RNeasy Plus Universal mini kit protocol after homogenization with silica beads and the TissueLyser. Library preparation and sequencing was performed as for blood above.

### Bioinformatics

#### Genome assembly

To automate assembly for our 46 individuals, we organized our workflow as a Snakemake pipeline available here: https://github.com/harvardinformatics/scrub-jay-genomics. To summarize, we trimmed any remaining adapters from our reads using `Cutadapt` (Martin 2011) and assembled using `Hifiasm` (Cheng et al. 2021) using the option to partially phase each sample, which generates a primary assembly plus two haplotype assemblies per individual. For the samples for which we had HiC data (one male and one female *A. woodhouseii* and our outgroup *C. yucatanicus*) we integrated the reads into the Hifiasm assembly to better phase the output. We converted the GFA output from Hifiasm to FASTA and re-named the contigs to match the ‘PanSpec’ naming convention to facilitate downstream analysis.

For our reference individuals, we improved contiguity by scaffolding with paired-end HiC data. We first filtered the HiC reads using scripts from the Arima and Esrice pipelines (as modified here: https://github.com/harvardinformatics/scrub-jay-genomics), then mapped each read pair against each draft genome individually using `BWA`. We then combined the individually mapped read pairs together and removed duplicates using `samtools` and converted the output to BED format using `bedtools` (Quinlan et al. 2010). The HiC BED file was then used for scaffolding with `Yet Another HiC Scaffolding tool (YaHS)` (Zhou et al. 2023). We visualized the HiC alignment output from `YaHS` using `JUICER` (Durand et al. 2016).

Finally, for our reference *A. woodhouseii* female, to further improve contiguity and to assign chromosome names we performed reference-based scaffolding using `RagTag` (Alonge et al. 2022c) against a draft version of the Florida scrub jay genome (Romero et al. 2024). Because the Florida scrub jay assembly was generated from a male bird, we performed a second reference-based scaffolding against the Hawaiian crow genome to scaffold the W chromosome (Sutton et al. 2018). Two contigs that were W-associated in the Hawaiian crow-scaffolded version were instead scaffolded as part of ch5 and chZ in the Florida scrub jay-scaffolded assembly; we manually trimmed those contigs off the W, then added the W scaffold to the Florida scrub jay-scaffolded genome to obtain our final assembly. In our final reference assembly (aphWoo1), chromosomes are named based on homology to the Florida scrub jay reference (Romero et al. 2024).

#### Genomescope

We applied Genomescope (Ranallo-Benavidez et al. 2020) to the individual fastq files to estimate heterozygosity and genome size. Jellyfish (Marçais et al. 2011) was used to count k-mers, using default parameters.

#### Dipcall

We used the program dipcall (Li et al. 2018) to estimate heterozygous positions and confident bases in each individual, and to call small variants and long INDELs between haplotype assemblies of each individual. Dipcall works by aligning a diploid assembly (in this case, the two haplotypes from a particular individual) to a reference genome (in this case, aphWoo1) with minimap2 (Li 2018), and then identifying heterozygous positions (where one haplotype carries a non-reference allele). Confident bases are estimated based on alignment coverage, where confident bases are defined as bases that are covered by one >=50kb alignment with mapQ>=5 from each parent and not covered by other >=10kb alignments in each parent. We built the variant call makefile with “make -j2” followed by “dipcall.kit/k8 and dipcall-aux.js vcfpair”. The output comprised a phased VCF and a BED file of confident regions which was then filtered with grep to remove sex chromosomes. Per-base heterozygosity for each individual was calculated as the number of heterozygous positions identified by dipcall, divided by the number of confident bases identified by dipcall.

#### PSMC and demographic analysis

To infer population size history with PSMC (Li and Durbin 2011), we generated a diploid consensus sequence by 1) applying “seqtk mutfa” to the reference fasta assembly, modified with the Dipcall VCF (processed by “vcf2snp.pl”); 2) masked using the BED file using “seqtk seq - cM”, and 3) converted to PSMC input format with “fq2psmcfa”. We then ran PSMC using the command “psmc -N25 -t15 -r5 -p “4+25*2+4+6” -o $SAMPLE.psmc $SAMPLE.psmcfa” and visualized the results with “psmc_plot.pl”. No bootstrapping was performed. Each line in the PSMC plot in Fig. 1C represents a different individual.

#### Tests for gene flow between *A. coerulescens* and *A. woodhouseii*

Gene flow between species could influence the distribution of SNPs and SVs and could violate models for estimating the distribution of fitness effects of variants, which usually assume single, isolated and panmictic populations. To obtain a set of SNPs that overlapped between ingroup and outgroup and would satisfy the topology (((P1,P2),P3),Outgroup) that is required by Dsuite (Malinsky et al.), we used bcftools (Danecek et al. 2021) to intersect two vcf files, one containing all ingroup scrub-jay sample genotypes and one containing genotypes for the outgroup CY sample. We then merged the resulting intersected vcfs using bcftools (Danecek et al. 2021) and thinned to 1 SNP per Kb using vcftools (Danecek et al. 2011) to generate a vcf with approximately 1 million SNP genotypes for the 45 samples (44 ingroup AI, AW, and AC) plus 1 Outgroup CY). We then removed the W and Z chromosome SNPs from that vcf and filtered to only bi-allelic genotypes (a Dsuite requirement).

Using this vcf as input for Dsuite, and treating missing sites as missing, we at first found a signature of significant gene flow between Florida and Woodhouse’s Scrub-Jays, but with only a slight deviation from the null (D = 0.05). To test whether this signature would remain if the dataset was further filtered for linkage (to reduce the likely-overpowered 1M SNP analysis), we further filtered the dataset to one SNP per 50Kb, and found that the signature of significant gene flow disappeared. Regardless, the direction of the excess allele sharing remained between Woodhouse’s and Florida Scrub-Jays. We believe that this consistent signal, that AI birds share few derived alleles with AC birds in all analyses, can be attributed to the fact that rates of lineage sorting were stronger in the AI lineage, which has undergone genetic bottlenecks and experienced repeated purging of broadly shared variants because of its evolutionary history on Santa Cruz Island. These demographic processes are likely strong enough to create a subtle deviation from the null expectation of D = 0 (Koppetsch et al. 2024), which is considered significant when we use a large SNP dataset (∼1M SNPs), but disappears in datasets that are linkage-thinned in closer accordance to a realistic estimation of the number of independent linkage blocks present in the genome. We therefore attribute the significant ABBA/BABA results to unaccounted-for demographic processes in the AI birds and conclude that there is no significant gene flow between AW and AC in this data set. This conclusion accords with previous findings in this system (McCormack et al. 2011, DeRaad et al. 2022).

#### Bayesian Phylogeography and Phylogenetics

A bed file containing high-confidence regions generated by dipcall was parsed to identify 1-kb autosomal loci that did not occur within 10kb of each other. This exercise generated ∼2500 loci, which were retrieved from each haplotype using command-line blast (Camacho et al. 2009). We attempted to extract the loci directly from the pangenome graph (Guarracino et al. 2021) but at the time of this work the odgi extract function was not working properly, and hence we used blast. The top-scoring and most complete blast hit was extracted from each haplotype. Loci were collated and aligned again using mafft v7.490 (Katoh et al. 2013) and then formatted for input into bpp v4.7.0_linux_x86_64 (Flouri et al. 2018), assigning each haplotype to its appropriate species. Before running bpp, the genealogy of each aligned locus was estimated using maximum likelihood using iqtree (Nguyen et al. 2015) to detect possible aberrant orthologs or misalignments. Rates of monophyly of alleles within each species was counted using functions in the R package ape (Paradis et al. 2004); the substitution model for each locus was estimated using ModelFinder (Kalyaanamoorthy et al. 2017).

In bpp, Model A00 was used, in which the species tree topology is provided and the program estimates branch lengths (in units of substitutions per site, μ*τ*, where μ=mutation rate and *τ*=divergence time in generations) and effective population sizes of extant and ancestral species (*θ*=4Nμ, where N is the effective population size). The results of the Markov Chain Monte Carlo (MCMC) run were summarized in Tracer v.1.7.1 (Rambaut et al. 2018). A substitution model for each locus (JC69, HKY, TN93) was assigned using the model closest to the results from ModelFinder.

#### Gene annotation

To generate a chain file for liftover, we aligned the final scaffolded *A. woodhouseii* reference genome (aphWoo1) against the *Gallus gallus* version 7b genome from NCBI (https://www.ncbi.nlm.nih.gov/bioproject/PRJNA660757/) using LASTZ (Harris 2007), implemented as a Nextflow workflow here: https://github.com/hillerlab/make_lastz_chains/tree/main. We first soft-masked repeats of the *A. woodhouseii* genome, as required for LASTZ alignment. Before performing liftover, we filtered the 80,621 chicken transcripts to remove noncoding RNA, resulting in 41,653 input transcripts. We then used the filtered transcripts and chain alignment file to annotate the *A. woodhouseii* reference using TOGA (Kirilenko et al. 2023). For the query annotation, we removed any transcripts shorter than ten amino acids in length, and filtered to only retain Intact (I), Partially Intact (PI) or Uncertain Loss (UL) projections, resulting in a total of 41,563 transcripts annotated in *A. woodhouseii*.

To ensure our annotation was as complete as possible, we also aligned *A. woodhouseii* against the zebra finch genome v1.4 (https://www.ncbi.nlm.nih.gov/datasets/taxonomy/59729/) (Warren et al. 2010), filtering the input transcripts similar to chicken (31,017 input transcripts with 48,058 isoforms). After performing liftover with TOGA we filtered the query annotation as above, resulting in a total of 42,730 annotated transcripts.

To produce our final combined, non-redundant annotation, we used `gffcompare` v0.12.6 (Pertea et al. 2020) (https://github.com/gpertea/gffcompare) to merge annotations, using the zebra finch as the reference species and the -D option to remove duplicate intron chains. For the merged GTF file, we assigned unique gene labels to each transcript using the orthology relations generated by TOGA, with labels determined by the type of orthology relationship identified by TOGA. For orthologs identified as ‘one2one’ (i.e. single copy in both reference and query species), we assigned the same gene ID as the reference species (with priority given to the zebra finch reference); for ‘one2many’ orthologs (i.e. genes duplicated in *A. woodhouseii*), we append a number to each copy of the gene. Orthologs with a ‘many2one’ relationship (i.e. multicopy in the reference species but single in *A. woodhouseii*) we append a “-like” suffix, and lastly for ‘many2many’ orthologs (multicopy in both reference and query), we append a “-like-NUM” suffix for each copy. In total, after merging we identified 34,669 transcripts across 15,966 genes. We ran BUSCO (Tegenfeldt et al. 2025) using the aves_odb10 dataset and obtained a score of 99.3%, indicating our annotation is highly complete.

#### PGGB Pangenome graph

To construct our pangenome graph, we combined the two haplotype assemblies generated by `hifiasm` for each of our 44 individuals across the three species, plus the single CY outgroup and our reference *A. woodhouseii* individual for a total of 91 haplotypes, which comprises >113 Gbase across ∼113k contigs. Because constructing an all-by-all graph from such a large dataset would be far too computationally intensive given our available computing resources, we first clustered our sequences into “communities” of related sequences, which we will then build into graphs individually. To partition into communities, we first performed an initial approximate all-by-all alignment of the sequences using `wfmash` (Guarracino et al. 2023; https://github.com/waveygang/wfmash), with the parameter for minimum percent identity set as `-p 94` based on expected divergence between the species and window size as `-w 1024` to improve runtime. In total, `wfmash` partitioned the combined haplotypes into 994 communities. We split these communities into three classes. Thirty communities correspond to chromosomes, as identified by chromosomal scaffolds from the reference individual, which we refer to as “chromosomal communities”. These chromosomal communities together contain 92.6 Gbase of sequence (i.e. ∼82% of the input sequence). Another 18 communities contain reference scaffolds that were not placed into chromosomal scaffolds, which we refer to as “unplaced reference communities”; these communities contain 13.2 Gbase of sequence, ∼11% of the total. The remaining 946 communities we refer to as “non-reference communities,” which have zero reference sequence in them and comprise only 7.3 Gbase (∼6.5% of the total). Most of these non-reference communities are quite small, with only a few contigs from a handful of individual haplotypes per community, and likely represent either spurious alignments between repeats or misassemblies or rare structural variants present in only a few assemblies. Thus, we chose to omit them when constructing the graph.

For each of the chromosomal and unplaced reference communities, we constructed a pangenome graph of the sequences using `PanGenome Graph Builder (pggb)` (Garrison et al. 2024; https://github.com/pangenome/pggb) using a minimum percent identity of -p 94 and a graph segment length of -s 100000. This produced a graph file in GFA format for each community containing indels, structural variants and single nucleotide polymorphisms. Due to computational constraints, we were unable to construct 4 of the 48 input communities: three unplaced reference communities and one corresponding to chromosome 17. These communities likely represent highly convoluted graphs that cannot be resolved by pggb. In the end we were able to obtain graphs for 30 of the 34 currently known chromosomes for the high-quality AC reference assembly (Romero et al. 2024).

In addition, we found segments of chromosomes 1, 1A, 2, 3, 5, 15, often at the ends of chromosomes and flanked by large inversions, with aberrantly low graph depth. Investigation suggested that these segments represent regions of the reference genome where our initial community construction algorithm failed to correctly place haplotype sequence into the reference community, potentially because of complex repeat structures or rearrangements. In order to account for this in downstream analysis, we produced a bed file with regions of low graph depth indicated, and used this bed file as a filter for all analyses.

#### Structural variant calling – PGGB

We use the paths in the graphs constructed by PGGB to call variants (Supp fig). For each community alignment, we take the GFA output and break it down into VCF file of individual variants using `vg deconstruct` v1.40.0 (Garrison et al. 2018; https://github.com/vgteam/vg), using the *A. woodhouseii* female reference (aphWoo1) as our base coordinates. To deal with overlapping and nested alleles in the graph, we ran vcfbub (Garrison et al. 2022) on the deconstructed VCF files to keep only the top-level variant sites (i.e. “snarls”) in the pangenome graph that are less than 100kb in size. We then used `vcfwave` (Garrison et al. 2022) to re-align the reference and alternate alleles to the genome; this process splits nested alleles into individual entries and identifies inversions (>1kb in size). Next we combined the VCF files from each community together using `bcftools concat` (Danecek et al. 2021; https://samtools.github.io/bcftools/bcftools.html) before concatenating, for any community with individuals missing (e.g. male samples for the W chromosome), we added in sample columns to those VCFs as missing data using `bcftools query`. For final cleanup, we used `bcftools +fixploidy` to set the same allele number for every site, `bcftools +fill-tags` to add allele count and allele frequency tags for each site and `bcftools norm` to split multiallelic records into biallelic.

Determination of ancestral states is critical to the designation of SVs as insertions, deletions or the SV inversion status. To determine ancestral states in our study, we used an outgroup that was approximately 12 MYA diverged from the ingroup species, and one that we learned in the process had a smaller assembly size than did the three ingroup species, resulting in a portion of SVs that could not be polarized, in part because the degree of divergence between the ingroup and outgroup species likely poses challenges to alignment and graph construction.

The use of a 12 MYA diverged outgroup struck a balance between being relatively closely related to the ingroup, thereby ensuring confidence in ancestral states, yet being distant enough to the ingroup so as not to engage in allele sharing, whether by incomplete lineage sorting or hybridization. Additionally, multiallelic SVs proved challenging to count. What might be considered one multiallelic SV in one pipeline could be considered two or more biallelic SVs another pipeline. Simple deconvolution of multi-allelic SVs into biallelic SVs is a solution, but may result in arbitrarily large numbers of SVs that may still be difficult to compare between species.

#### Structural variant calling – Minigraph and other methods

We ran minigraph v. 0.20-r559 on a total of 92 haplotypes (AI, AC, AW and CS) and the AW reference. We used the -j option (-j.05) to accommodate larger sequence divergence between species. The graph (in .gfa format) is available on Github. To call SVs, each haplotype is mapped back to the graph to produce a VCF-like file (*.gaf). The 93 *.gaf files are then merged to create a final VCF. The VCF contains insertions, deletions and inversions relative to the AW reference. In the bed file generated by mingraph, when a bubble involves an inversion, column number 6 indicates a value of 1.

To run the SyRi pipeline, we generated 45 pseudo-chromosome-level assemblies using RagTag (Alonge et al. 2022a), a scaffolding tool that maps query scaffolds to a high-quality reference genome. The assemblies were aligned to the reference using minimap2 with the preset –eqx option, and the output was sorted into BAM files with samtools. Next, SyRI was executed on each BAM file to detect structural rearrangements. The inversions were extracted from the SyRI output using awk (awk ‘$11==“INV“’).

#### Recombination

We used RELERNN (Adrion et al. 2020) to estimate per-base-pair recombination rates, only for the AW genomes, which had the most information and variability for estimating recombination rates. We used VCF files parsed by species and by chromosome and for biallelic sites with no missing data as input for RELERNN. We first used the SIMULATE function to train the algorithm, producing 13,000 training data sets, 2000 validation sets and 100 for testing. We predicted recombination rates (rho) for each chromosome and conducted bias correction using the BSCORRECT module, using a mutation rate of 2.2e-9 for noncoding DNA (Nam et al. 2010) and an upper limit for rho of 5.17. Recombination rates were plotted on a 1-Mb sliding window and averages per chromosome. Correlations of SNP and SV density with recombination were conducted by counting the number variants per Mb from the PGGB VCF and plotting with recombination.

#### Kinship and runs of homozygosity

We used PLINK2 (Chang et al. 2015) to estimate runs of homozygosity by the method of and kinship coefficients using KING (Manichaikul et al. 2010). The vcf file projected from the combined PGGB pangenome graph was parsed for biallelic autosomal SNPs. A table of relatedness was generated by PLINK2 within each species separately. Runs of homozygosity were estimated using default parameters.

#### RNAseq analysis

To evaluate transcript expression levels, we quantified transcript abundance for our sixty two samples from a variety of tissue types using `kallisto` (Bray et al. 2016). We indexed our merged TOGA transcriptome with `kallisto index` and used `kallisto quant` with 30 bootstrap replicates to estimate transcripts per million (TPM) and expected counts for each transcript in each sample.

#### Repeat annotation

RepatModeler2 (Flynn et al. 2020) was run on the AW reference genome to generate a de novo repeat library. We also ran Satellite Repeat Finder (SRF; Zhang et al. 2023) on the reference assembly as well as on each haplotype assembly and individual sets of unassembled PacBio HiFi reads. The SRF satellites were concatenated to the RepeatModeler library to obtain the full library used or annotation. In downstream analyses, satellites identified by RepeatModeler2 were removed from the library so as not to compete with the SRF satellites, which were generally longer and more diverse.

RepeatMasker was used to annotate each primary assembly, including the AW reference, as well as each haplotype assembly. For all analyses the out file produced by RepeatMasker was converted to *.bed format using rmsk2bed function of the bedops package (Neph et al. 2012) for downstream processing with custom R scripts.

#### Analysis of satellite landscapes

We used minimap2 to map the 300 satellites detected by SRF to PacBio HiFi reads. These outputs were then parsed with xx tools to produce *.len files consisting of columns designating the satellite; the total number of base pairs covered by the satellite in a given fastq.gz file; a rough measure of sequence divergence among the satellite copies; the proportion of read base pairs comprised of the satellite, excluding reads or contigs in which only one copy of the satellite repeat unit is found; and finally the number of base pairs comprised by the satellite including all reads or contigs in which only any number of copies of the satellite repeat unit is found. The *.len tables from individual birds were concatenated and the resulting tables parsed to examine the distribution of satellites across individuals and species.

Inspection of srf outputs and mapping of satellites by minimap2 indicated that satellite sj_sat_circ30_18193, which has an ∼18kb unit repeat, varied drastically between species and sexes. We mapped the distribution of this repeat to each haplotype using minimap2 and the command: minimap2 -c -N1000000 -f1000 -r100,100 18kb.fa ctg.fa > srf-aln.paf. The resulting .paf files were parsed for satellite unit copies that were at least 18 kb. There were 27,083 full-length unit copies of this satellite across AI, AA, AW, AC, CY, and CS haplotypes. These ∼27,000 units were extracted from haplotype assemblies using a bed file and seqtk sample. They were then aligned with mafft, using the option –adjustdirectionalityaccurately. To reduce the number of sequences to be analysed phylogenetically, we used cd-hit (Fu et al. 2012)to isolate 3,500 unit sequences that were less than 0.999 similar to one another. We used the command line:

`cd-hit -i all_sj_haps_18kb_cat_pos_neg.fa -o all_sj_haps_18kb_cat_pos_neg_cdhit -T 4 -c 0.999 -M 100000 -n 5 -d 0`

These 3,500 sequences were analyzed with IQ-Tree (Minh et al. 2020) with a GTR model [GTR+F+R9] determined by ModelFinder (Kalyaanamoorthy et al. 2017). The tree was depicted using functions in the R packages ape (Paradis et al. 2004) and ggtree (Yu et al. 2017).

#### Telomeres

Telomeres were initially quantified using Satellite Repeat Finder and RepeatMasker. One of the ∼300 satellites identified by SRF, sj_sat_circ86-113, corresponded to the canonical vertebrate telomere motif (TTAGGG)n, and was used either as baits in command-line blast searches (Camacho et al. 2009) or quantified using SRF satellite distributions. During the analysis phase of this project, seqtk telo was added to the seqtk set of scripts (Li 2013) and subsequently was used for all telomere analyses. Seqtk telo counts telomeric sequences occurring at the ends of contigs or scaffolds. It currently ignores telomeric sequence not occurring at the ends of reads, contigs or scaffolds. Telomere abundances were calculated as the fraction of telomeric sequence divided by the total base pairs in the target, whether HiFi reads, haplotypes or primary assemblies.

To study the effect of individual age on telomere abundance, we sequenced with PacBio HiFi two older birds that were known to be at least 11 years old so as to increase the age range of birds in this study. Each bird was first sequenced to moderate (∼10X) coverage with PacBio Sequel II technology, then followed with a single flow cell each on a PacBio Revio machine. One AC bird (AC_49710) was found as an immigrant to the Archbold Field Station site on 9/19/2012 and was subsequently banded on 4/11/2013. Blood was sampled from this individual on 5/10/2023, making it at least 11.06 years old at the time of sampling. Another AC bird, AC_49358, was known to have hatched on 4/24/2012 and was sampled for blood on 5/9/2023, making it 10.65 years old at the time of blood sampling.

#### Panacus plots

We estimated the pangenome growth curve and core size using panacus (Parmigiani et al. 2024) on PGGB .gfa file. To compare growth curves among scrub jays, humans, and chickens, we obtained a human pangenome graph (PGGB GFA; https://github.com/human-pangenomics/hpp_pangenome_resources; Liao et al. 2023) containing 44 samples from four populations (Punjabi from Lahore, Pakistan [PJL], East Asian [EAS], Admixed American [AMR]), as well as a chicken pangenome (PGGB GFA; https://zenodo.org/records/10018222; Rice et al. 2023) containing 18 samples. We then performed panacus analyses on both the human and chicken pangenomes.

#### Constructing site frequency spectra for SNPs and SVs

We wrote a custom script (available at https://github.com/harvardinformatics/scrub-jay-genomics) to parse the PGGB VCF file into a tabular format facilitating plotting and tabulating. This table (available in Supplementary Data), in bed format with each row consisting of a single variant (SNP, indel or SV), recorded the overlap of each SV with annotations, such as CNEEs, exons, introns or intergenic regions, as well as overlaps with RepeatMasker repeats. The table also recorded the ancestral and alternative alleles as indicated in the outgroup (CY), as well as the type of variant (DEL, DEL_COMPLEX, INDEL_COMPLEX, INS, INS_Complex, SNP, SV_Complex, SVDEL, SVDEL_Complex, SVINS, SVINS_Complex). There are columns recording whether or not the variant represents an inversion; whether the variant is polarized; the ancestral and derived allele(s); the base and alternate allele length(s); maximum length of variants; and the allele count. Additionally, the derived allele count (DAC) was recorded for each species (AW, AC and AI), as well as the number of haplotypes per species missing for a given variant, and the total number of alleles for the variant. We constructed SFSs for various classes of variation from this table. To compute unfolded (polarized) SFS, we first filtered to retain only biallelic variants (allele number = 2) that are polarized (polarized = TRUE). For most analysis, we also filter to only retain sites with no missing data. We then summarized counts of sites in each frequency bin (derived allele count, summarized separately for each of AC, AI, and AW) for SNPs (allele length = 1), INDELs (allele length > 1 and < 50), and SVs (allele length >= 50).

#### Estimating the Distribution of Fitness Effects

We employed the maximum-likelihood programs fastDFE (Sendrowski and Bataillon 2024) and anavar (Barton and Zeng 2018) to estimate the distribution of fitness effects (DFEs) for SNPs, INDELs, and SVs, following the methods detailed in Fang & Edwards (2024) (Fang and Edwards 2024). Both methods use the site frequency spectrum (SFS) to infer the population-scaled mutation rate (θ = 4Nₑμ, where Nₑ is the effective population size and μ is the per-site, per-generation mutation rate) and to fit shape and scale parameters of a gamma distribution for the population-scaled selection coefficients (γ = 4Nₑs, with s being the selection coefficient). These approaches also control for demographic factors and polarization errors, following the method outlined by Eyre-Walker et al. (2006) (Eyre-Walker et al. 2006).

We generated the unfolded SFS from the PGGB VCF for each species, targeting variants grouped by size class (SNPs, INDELs, SVs) and genomic region (CNEE, exon, intron) as described above. Only polarized, bi-allelic variants with complete data were included in the SFS and subsequent DFE analyses. In fastDFE, we fitted the SFS to the ‘GammaExpParametrization’ model; in anavar, we fitted the SFS to ‘neutralSNP_vs_selectiveSNP’ for SNP datasets and ‘neutralINDEL_vs_selectedINDEL’ for INDEL and SV datasets, applying a continuous gamma distribution. Both programs used a putatively neutral reference SFS derived from intergenic variants to account for demography and polarization errors.

We report the gamma distributions as the proportion of variants within four bins of scaled selection coefficients (γ): neutral (0 ≤ −Nₑs ≤ 1), weak (1 < −Nₑs ≤ 10), moderate (10 < −Nₑs ≤ 100), and strong (−Nₑs > 100). In anavar, these bins are the default output; in fastDFE, we specified them as c(−Inf, −100, −10, −1, 0). All estimates of γ are negative, reflecting the deleterious nature of the mutations, based on the assumption that beneficial mutations are typically rare and subject to strong positive selection, leading to their rapid fixation in the population and minimal contribution to DFE (Booker 2020, Robinson et al. 2023) We derived 95% confidence intervals via parametric bootstrapping in fastDFE (using the inf.bootstrap function) and by gene permutation in anavar. Finally, using the same procedures, we also inferred DFEs for variants subdivided into size classes from 1–100 bp.

#### Detecting and quantifying inversions

We used (PGGB (Eizenga et al. 2021), minigraph2 (Li et al. 2020), SyRi (Goel et al. 2019) and svim-asm (Heller and Vingron 2021)) to identify inversions across the three core *Aphelocoma* species. We also used lostruct (Li and Ralph 2019) to identify large inversions. The PGGB vcf flags inversions when using recent versions of vcfwave (Garrison et al. 2022), as of July 2023. The decomposed vcf produces a comment in the information column, “INV=YES”, which was used to count inversions and retrieve their coordinates. Similarly, minigraph2 also lists inversions in the outputted vcf file. SyRi uses whole-genome alignment between a reference and a query genome to detect inversions. We broadly followed methods in (Fang and Edwards 2024).

To produce chromosome-scale inputs for SyRI, we first scaffolded our haplotype assemblies with RagTag (Alonge et al. 2022b); this will in principle normalize inversions between the haplotype and the reference when no contig in the haplotype assembly spans a breakpoint, but in these cases we have limited power to detect inversions in the first place. These scaffolded haplotype assemblies were aligned to the AW reference using minimap2 with the preset –eqx option, and the output was sorted into BAM files with samtools (Li et al. 2009). Next, SyRI was applied to each BAM file to detect structural rearrangements, including inversions. The inversions were extracted from the SyRI output using awk (awk ‘$11==“INV“’).

Svim-asm works with minimap to detect inversions and other SVs. We first mapped each haplotype to whichever reference was being used using minimap2; we used the AW reference for most analyses, but also used the CY outgroup and the recently published genome of *Cyanocitta stelleri* (Steller’s Jay; Benham et al. 2023). The bam files produced by minimap2 were used as input for svim-asm in diploid mode; each run used the two bam files produced by the two haplotypes of each individual. The minimum variant size was set to 50 bp. Options --query_names and --symbolic_alleles were used to aid in vcf interpretation. Vcf files were parsed and inversions flagged as incomplete were retained and included in the final counts.

Lostruct uses multidimensional scaling (MDS) to infer subgroupings of haplotypes or individuals that diverge in their snp profiles, which may indicate an inversion. The main PGGB VCF was parsed by chromosome and by species, retaining only biallelic snps with no missing data. VCF files were loaded into R using the ‘read_vcf’ function and eigenvalues were calculated for sliding windows of 5,000 or 50,000 bp, depending on the SNP density in the VCF. Principal components were calculated with for each haplotype using ‘pc_dist’. These PCs were then plotted on a 2-dimensional plot using the ‘cmdscale’ function and saved. The coordinates of the resulting MDS plot were converted to genomic coordinates to aid in inversion location.

The resulting MDS plots each chromosome/species were visually inspected for abrupt shifts in the value of MDS1 that could indicate an inversion. Breakpoints in the MDS plots were estimated using the Bayesian breakpoint analysis tool mcp (Lindeløv 2020). MDS plots were visually inspected and the posterior distribution of the breakpoint(s) was manually shifted using strong priors when necessary. (MCP alone often placed breakpoints earlier in the chromosome scan than expected given the major shift and number of potential shifts in MDS values across the chromosome. The mean position of each breakpoint was recorded and converted to a bed file for each chromosome.

The bed files were used to calculate principal components for each putative inversion genotype for each individual using SNPRelate (Zheng et al.). Heterozygosity for each putative inversion genotype was estimated using VCFtools (vcftools --het.; Danecek et al. 2011).

#### Pangene

Pangene (Li et al. 2024) aligns protein sequences to genome assemblies to produce a graph of coding regions for a collection of genomes. We aligned annotated protein sequences from TOGA to each haplotype using miniprot (Li 2023), which creates a protein .paf file. This .paf file was used as input for pangene, which produces a .gfa file. The pangene-1.1-bin-sj.tar.bz2 file, shared on Dryad, contains instructions and data for building a local version of an interactive tool to explore the pangene results.

### Supplementary Text

#### Phylogenetic considerations and statistical testing

Throughout, we did not use phylogeny when conducting statistical tests of various traits between species (repeat abundances, assembly size, telomere abundances, etc), primarily because our data structure is very different from what is typically used in such tests. Current phylogenetic models, such as phylolm (Ho et al. 2014), are tailored to phylogenies containing many species, each represented by a single individual or allele. By contrast, the data analyzed here in most cases consists of only 4 species (AI, AW, AC and CY), each represented by 1-15 individuals (2-30 alleles per species). Although some have argued that this data structure can be accommodated by using a star phylogeny within species, in practice this approach often breaks existing packages, including phylolm, yielding errors. Additionally, which tree and which branch lengths best describes a tree of a few species and many individuals is not clear; presumably some summary tree could be used, but given the extensive variation in gene trees across the genome, and the prevalance of incomplete lineage sorting (in which alleles within species coalesce deeper than between species), choosing which tree to represent the data set was problematic. We therefore used standard statistical testing throughout in lieu of phylogenetic models more appropriate for our data.

#### Calibrating *Aphelocoma* divergences against primates

To place sequence divergence in *Aphelocoma* in the context of commonly studied primate species, we compared divergence in *Aphelocoma* and the CY outgroup directly to those of primates across 415 orthologous genes, using TOGA (Kirilenko et al. 2023) and mafft (Katoh and Standley 2013). We calculated distances using the ‘dist.dna’ function in the R package ape (Paradis et al. 2004). We found that mean divergence across genes between AC and the CY outgroup (mean 0.0137 substitutions per site) was 55.5% of that between orangutan and human (mean 0.0247); similarly, the common ancestor of AC and AW (mean across genes, 0.00493) was only 54.4% as deep as the ancestor of human and chimpanzee (mean 0.00907). These results are summarized in (Supplementary). We retrieved absolute divergence times for *Aphelocoma* and outgroups from TimeTree5 (Kumar et al. 2022). Absolute divergence times of the common ancestor of AC and AW (∼5.2 MYA) and with the CY outgroup (∼12 MYA) were similar to those of human and chimp (∼6.4 MYA), and orangutan (∼15 MYA), respectively (Kumar et al. 2022).

#### Sensitivity of the PGGB vcf projection to different reference assemblies

Before estimating the average fitness effects of SVs and indels, we first confirmed that our method for projecting variation from the PGGB pangenome graph to VCF format, which requires designating both a reference haplotype and an outgroup, was not strongly biased towards or against calling fixed and polymorphic SVs in the reference or other species. Comparisons of counts of fixed and polymorphic SNPs and SVs across the 50 million bp of chromosome 7 suggest that the ratios of fixed and polymorphic variant counts derived from the PGGB pangenome graph, which used the AW HiC assembly as a reference but the CY bird as an outgroup, were comparable to those using the CY bird as both reference and outgroup (Supplement). We therefore ended up using the AW reference to obtain SV counts, while still using the CY as an outgroup to designate ancestral states. By using the AW reference to obtain SV counts, we are able to capture substantially more SVs in our analyses of selection, because its assembly is 11% larger than that of CY.

#### Characterizing a chromosomal fission in *A. woodhouseii*

Inspecting the .paf output files from minigraph, and using the paftools.js misjoin tool, we found that many contigs from AI and AC, but not from AW, mapped to both chromosome 27 and 28 of the AW reference, with the same strand of the query of single contigs mapping to parts of both AW ch27 and ch28. We found 21 AC haplotypes (21/28 = 75%) and 10 AI haplotypes (10/30 = 33%) that spanned the two AW chromosomes. We could find 9 AC birds (AC_1713_89780, AC_1873_10702, AC_1873_10717, AC_1873_10752,AC_1873_10831, AC_1873_20930, AC_1873_20934, AC_1873_20946,and AC_1873_20970) in which individual contigs of both haplotypes spanned the two AW chromosomes, but only 1 AI bird (AI_1833_00687).

A parsimonious explanation for these patterns is that a chromosomal fission is present in the AW reference that is not present in at least some AI and AC birds. Because the minigraph misjoin tool requires full contigs spanning the regions, we reasoned that the absence of contigs spanning the two AW chromosomes in some birds could be due to fragmented assemblies in this region. However, we confirmed the overall patterns found with minigraph by interrogation of community 14, which includes both AW chromosome 27 and 28, possibly due to a ‘telomere kiss’ – the joining of ends of chromosomes into a loop because of their sequence similarity (Fig. 4J). We remade the pangenome graph of community 14 using more stringent parameters to improve alignment quality: pggb -p 95 -s 10k or -p 94 -s 10k. We then used odgi layout

We then used odgi draw to produce a two-dimensional layout of the graph, coloring specific segments to learn more about which contigs in the community spanned which regions of the layout. In particular we identified specific contigs in AI birds that spanned the AW ch27/28 junction:

`odgi draw -i allbird_community.14.final.*.smooth.final.og -c allbird_community.14.final.*.smooth.final.og.lay -p colored-layout.png -b ../allbird_community.14.bed`

Alignments of at least 250kps against both chr27 and chr28 were deemed evidence of lack of a chromosomal fission in these individuals. The odgi draw analyses complemented those of minigraph. Odgi draw could not detect evidence for contigs in AI_1603_79174#hap1 that span AW ch27/28, but it detected four additional instances in AC_1873_20908#hap2, AC_1873_20954#hap1, CY_8788#hap1, CY_8788#hap2.

Further interrogation with minigraph found evidence for contigs spanning the AW ch27/28 reference in both CY haplotypes (contigs CY_8788#1#h1tg000184l and CY_8788#1#h1tg000184l) and one CS haplotype (CS_1#1#h1tg000047l), as well as one haplotype of CS and both haplotypes of AA. We further used odgi draw to better define the coordinates of contigs spanning the putative fission and repetitive regions in the community 14 graph. We established in the community.14 layout that the “bird’s neck” arises due to sequences present in only in the two CY_8788, whereas the region where the haplotypes switch between AW ch27/28 chromosomes is on the bottom.

No AW contigs were found to span both AW reference ch27 and ch28 chromosomes, suggesting that the fission is fixed or nearly fixed in AW. The frequency of the fused condition is uncertain in the remaining species, but our provisional hypothesis is that a chromosomal fission of ch27 and ch28 is derived within AW, whereas the other species retain the ancestral, fused condition.

#### Gene expression and gene copy number

Because the pangene results were constructed based on the best-mapping protein isoform for each gene, and the kallisto results were generated on all transcripts, we first normalized the kalliso and pangene results to create a single estimate of copy number variation and gene expression for each annotated gene. We then verified with PCA that samples correctly cluster by species and tissue in expression space, and checked for aberrant outlier samples. Because we lack power for a per-gene analysis due to the limited sample size, we decided to quantify the relationship between copy number and expression in aggregate using mixed effect linear models, filtered to examine only genes with copy number variation. We first fit a series of models to each tissue with expression (log(TPM)) as the response variable, a fixed effect of count as the main predictor, a random effect of gene (to allow each gene to have its own intercept), and optionally a main effect of sex. We then fit an aggregate model that pools across tissue and sex, revealing a significant main effect of gene copy number on log10(TPM) (p = 0.0476). The model estimated that log10(TPM) increases by 1.5e-02 with each gene copy. Some individual tissues also revealed positive relationships between gene copy number and log10(TPM): female gonad, brain, heart (with sex as covariate), but not male gonad or eye. Full analysis details, including exploratory data analysis and quality control checks, are available as an R markdown document at https://github.com/harvardinformatics/scrub-jay-genomics.

There is some noise in either in our identification of gene deletions or in our estimates of gene expression, or both: we find a small number (n=30) of genes exhibiting TPM > 1 (log10 TPM > 0) in at least one tissue in AW individuals putatively homozygous for a given gene deletion, a discrepancy that may arise due to mismapping of RNA-seq reads, which is particularly likely when mapping to genes with copy number variants.

### List of Supplementary Tables (in Excel file)

table S1. Sample details

table S2. PacBio HiFi read and hifiasm assembly statistics

table S3. Statistics of gene trees from 2489 1-kb dipcall regions used in bpp analysis

table S4. RepeatMasker results per haplotype

table S5. Comparison of scrub-jay satellites detected with srf and other corvid satellites

table S6. Ages of individuals (if known) and telomere abundances in reads and haplotype assemblies

table S7. Comparison of divergences of scrub-jays and primates at 415 orthologous genes

table S8. Counts of SNPs, indels and SVs from PGGB VCF file

table S9. Summary of SVs detected with minigraph and svim-asm

table S10. Estimates of alpha and direction of selection on indels and SVs.

table S11. Length statistics of SNPs, indels and SVs from the PGGB VCF

table S12. Minigraph mapping evidence for a chromosomal fission in AW REF ch27 and ch28

table S13. Results of pangene analysis of all haplotypes

table S14. Effect of filtering on pangene summary statistics

### Supplementary Figures

**Fig. S1.**
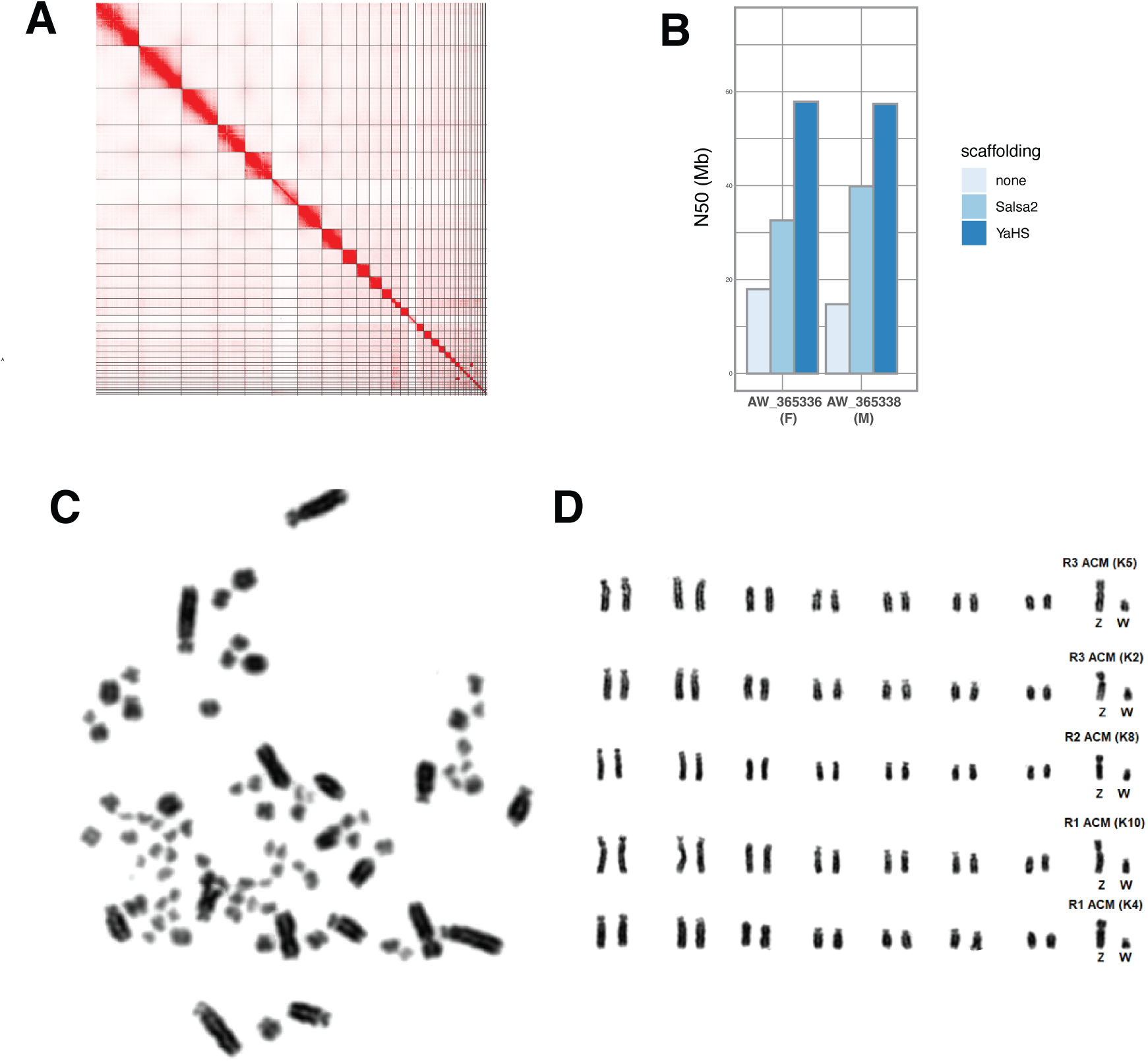
HiC analysis of reference genomes and karyotypes. **A**, Juicer (Durand et al. 2016) plot of putative chromosomes of the AW reference. **B**, Improvement of contiguity (N50) of three PacBio HiFi assemblies with additional HiC data. The AW (F) reference was used throughout the study. **C**, Female (MCZ 366861) metaphase chromosome spread. The spread suggests a diploid number of 2N=80. **D**, Replicate female (also MCZ 366861) metaphase spreads, organized by largest 7 autosomes and sex chromosomes.

**Fig. S2.**
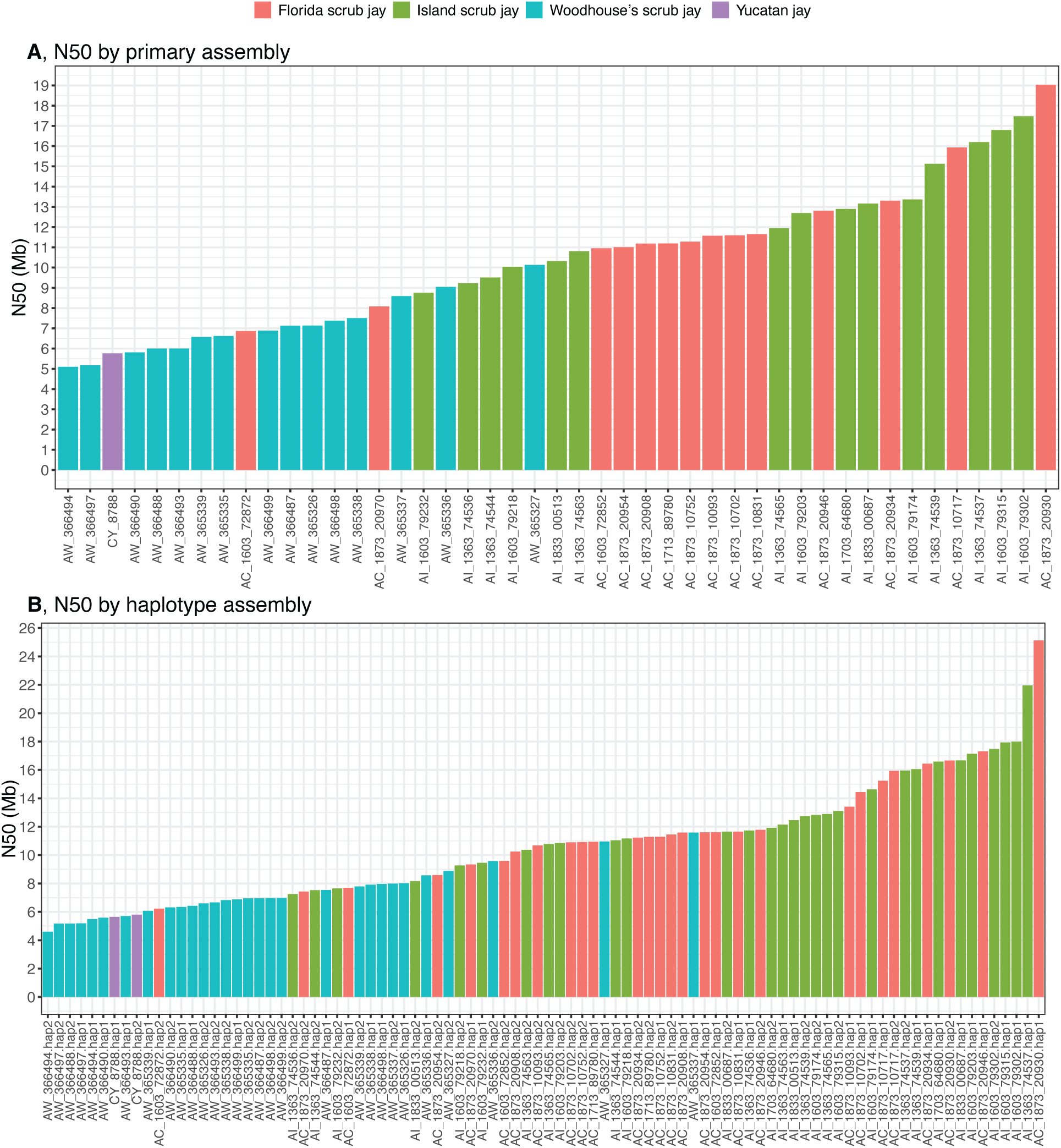
Assembly quality by haplotype and individual. N50 of **A)** primary assemblies and **B)** haplotype assemblies produced by hifiasm (Cheng et al. 2021).

**Fig. S3.**
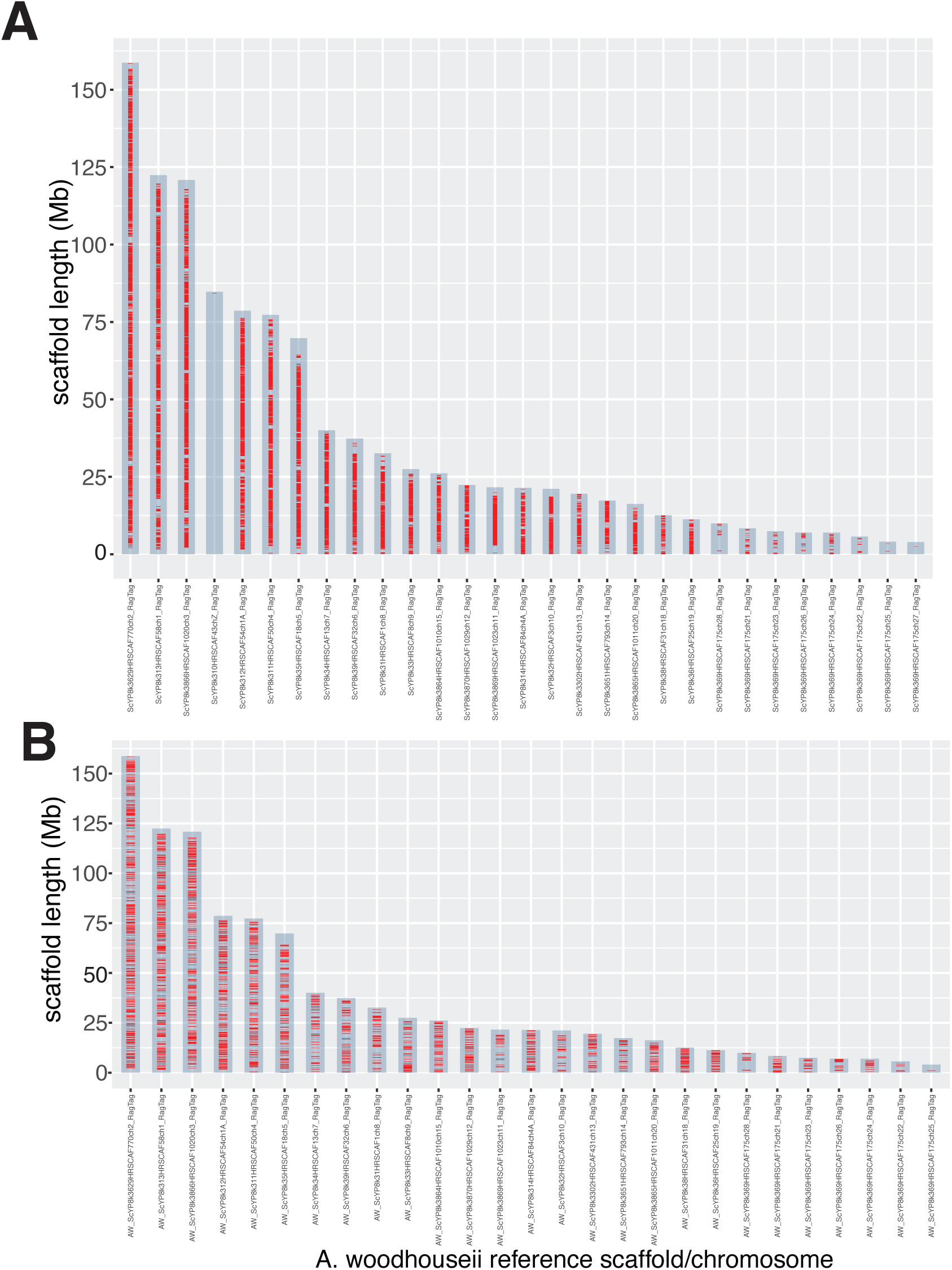
Dipcall results. Distribution of **A**) Dipcall regions and **B**) downsampled Dipcall regions among chromosomes. Each dipcall region is indicated by a red horizontal bar, but bar widths are not proportional to length of dipcall region. Downsampled regions (B) were used for bpp analysis.

**Fig. S4.**
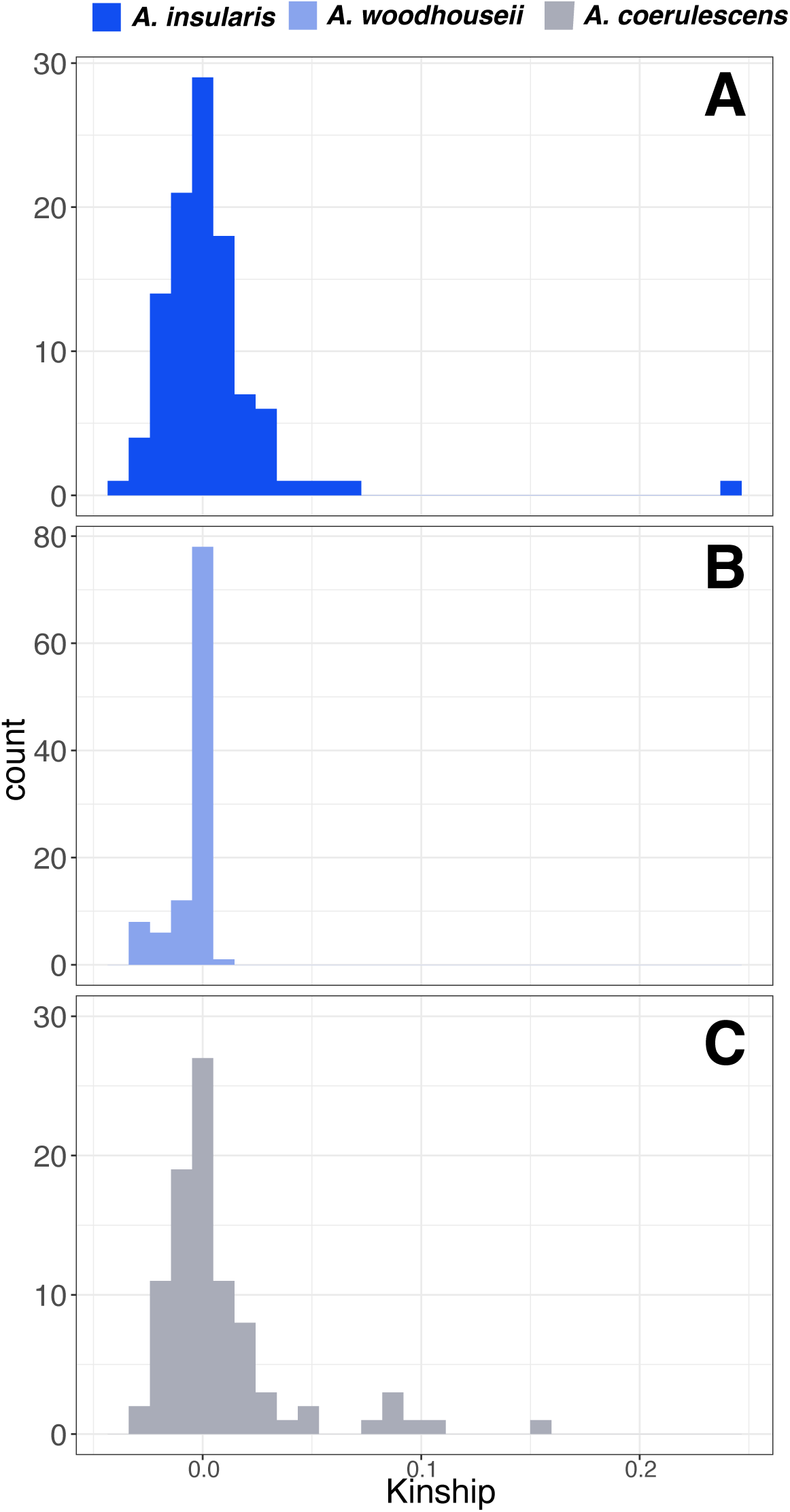
Estimates of kinship. Kinship was estimated with KING (Manichaikul et al. 2010) for all dyads within each of the three *Aphelocoma* species.

**Fig. S5.**
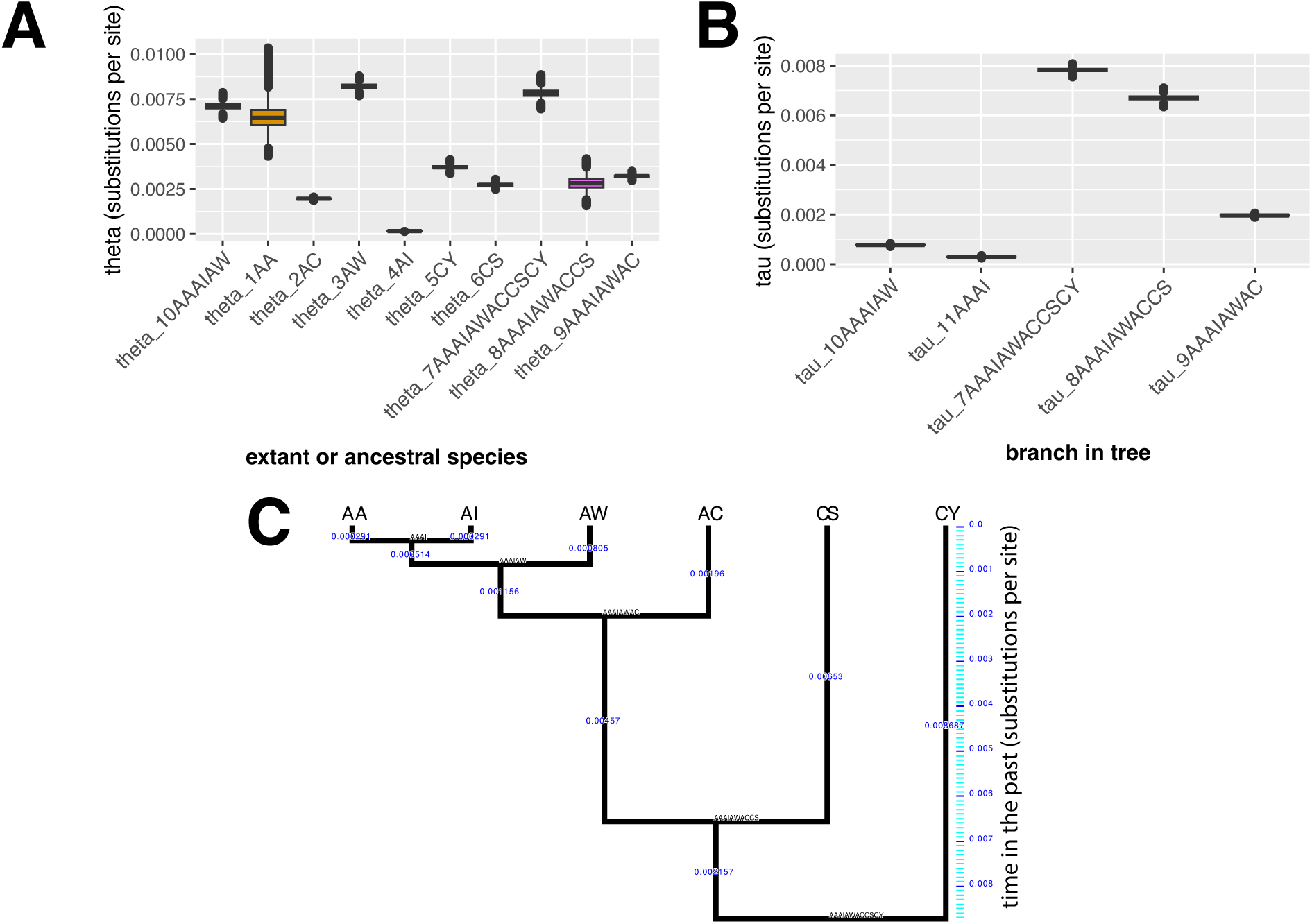
Results of bpp analysis. A, Posterior distributions of theta (*θ* = 4Nμ) for each extant or ancestral species in the species tree. B, posterior distributions of divergence times (*τ*) of the five branches leading from AA to the root of the tree. The tree is ultrametric so these five branches determine the shape of the entire tree. C, estimated branch lengths of species tree in units of substitutions per site.

**Fig. S6.**
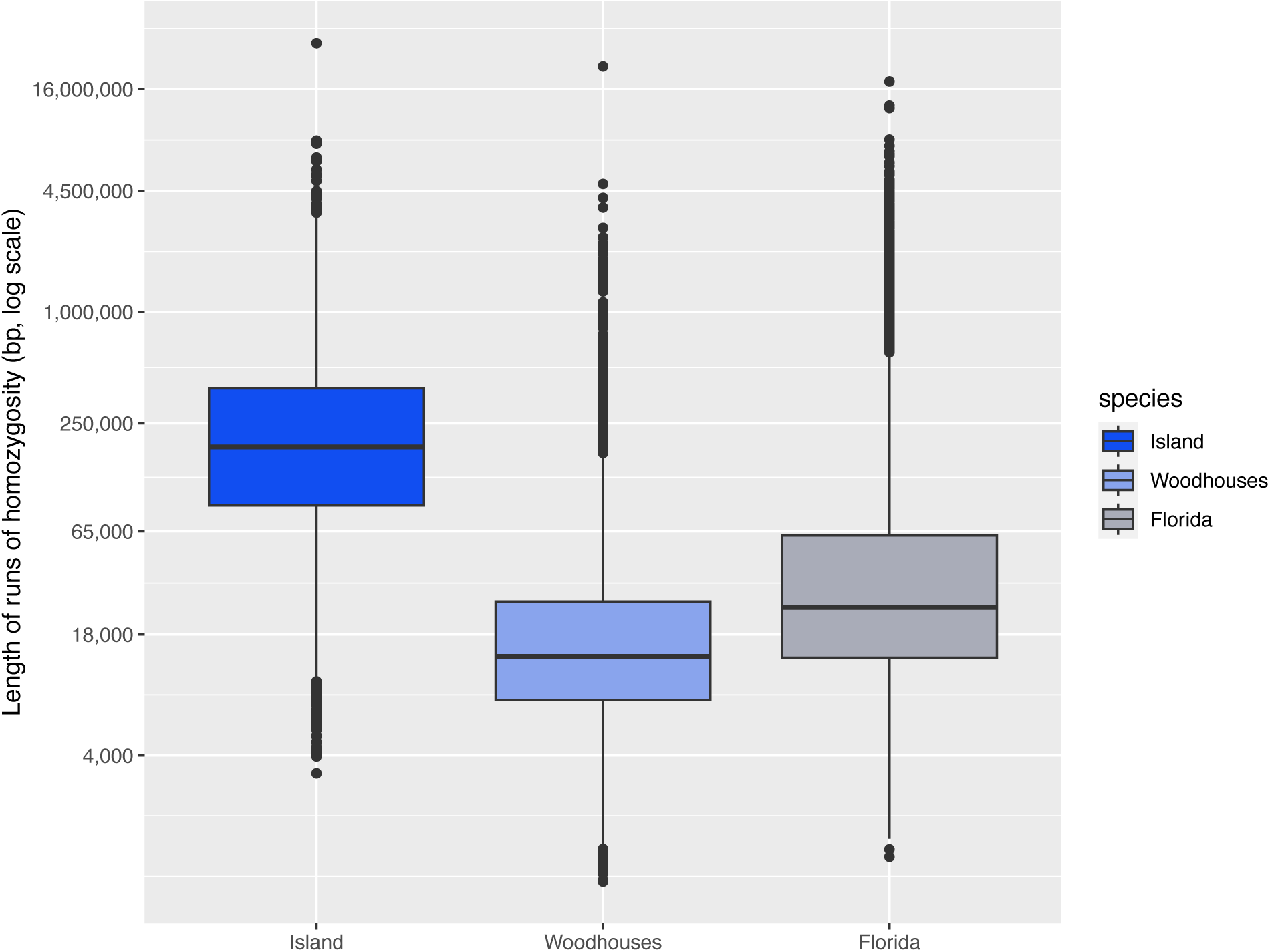
Runs of homozygosity. Distribution of runs of homozygosity in (left to right) *A. insularis*, *A. woodhouseii* and *A. coerulescens*.

**Fig. S7.**
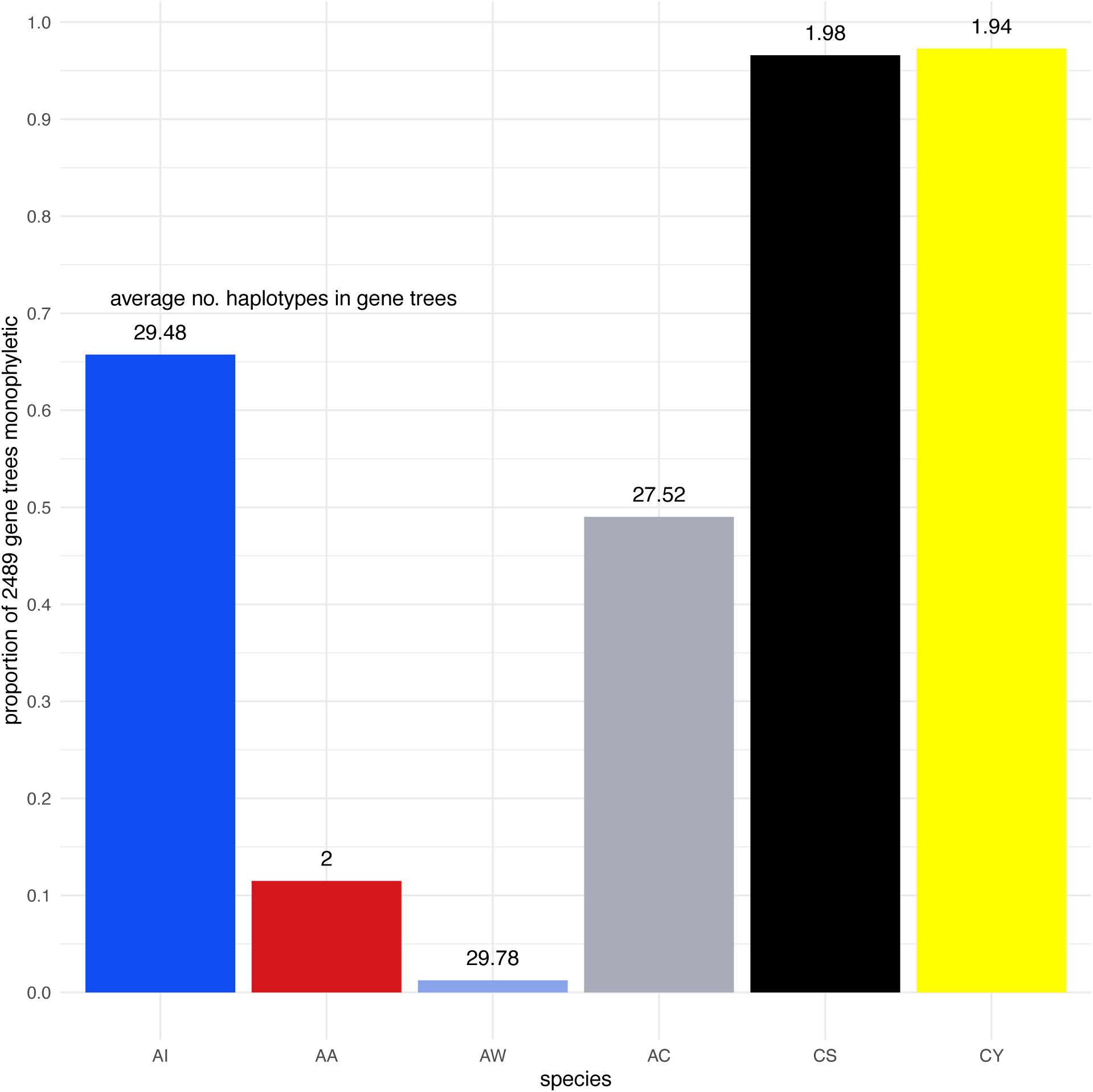
Proportion of gene trees monophyletic. Trees were made from 1-kb in dipcall regions and analyzed with IQ-Tree (Minh et al. 2020). The average number of haplotypes per gene tree is indicated above each bar.

**Fig. S8.**
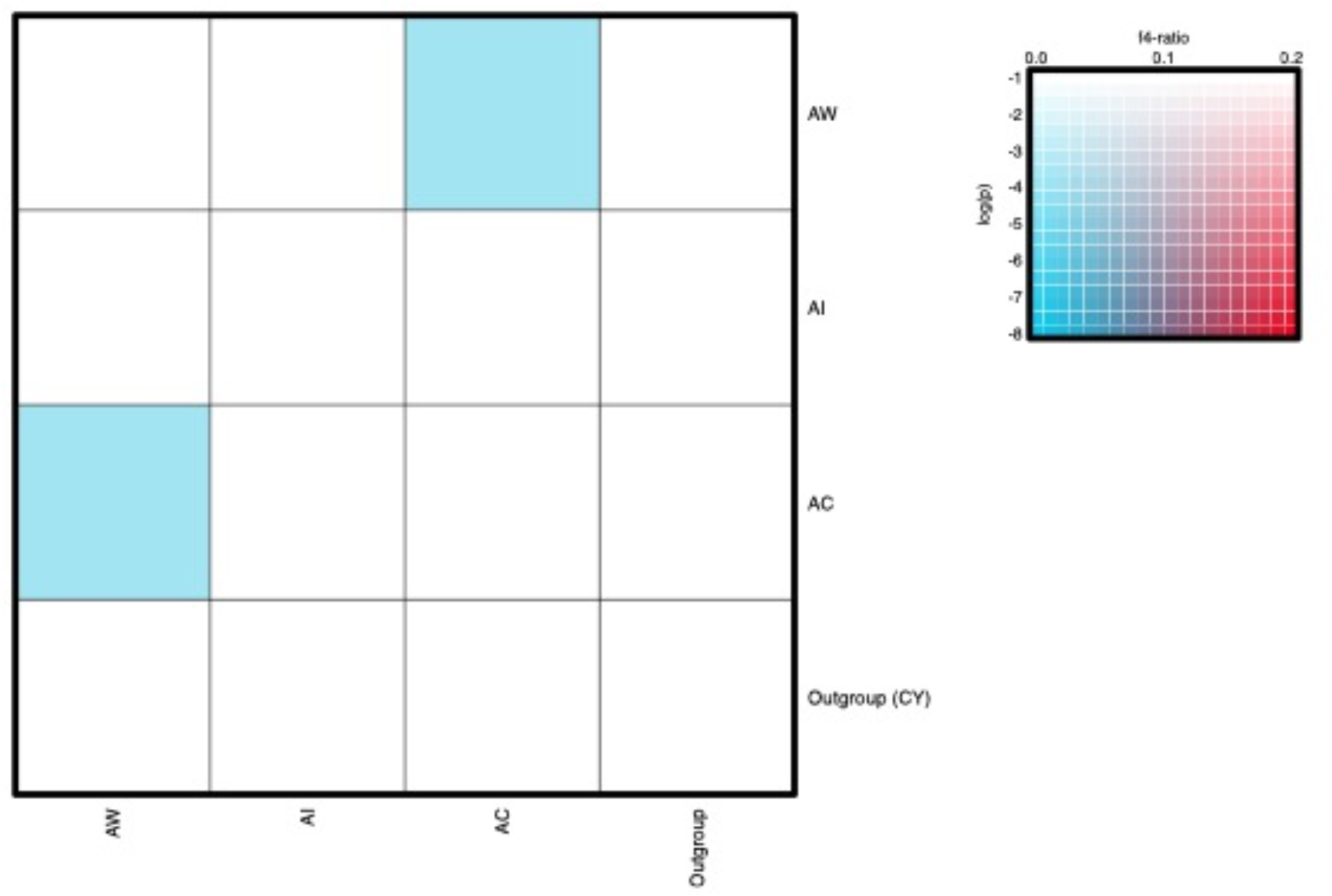
Tests of gene flow with f4 statistics. The above plots are for the unthinned VCF analysis (see Methods), which yielded a significant signal of gene flow between AW and AC. In the thinned analysis, this signature disappears (see Methods).

**Fig. S9.**
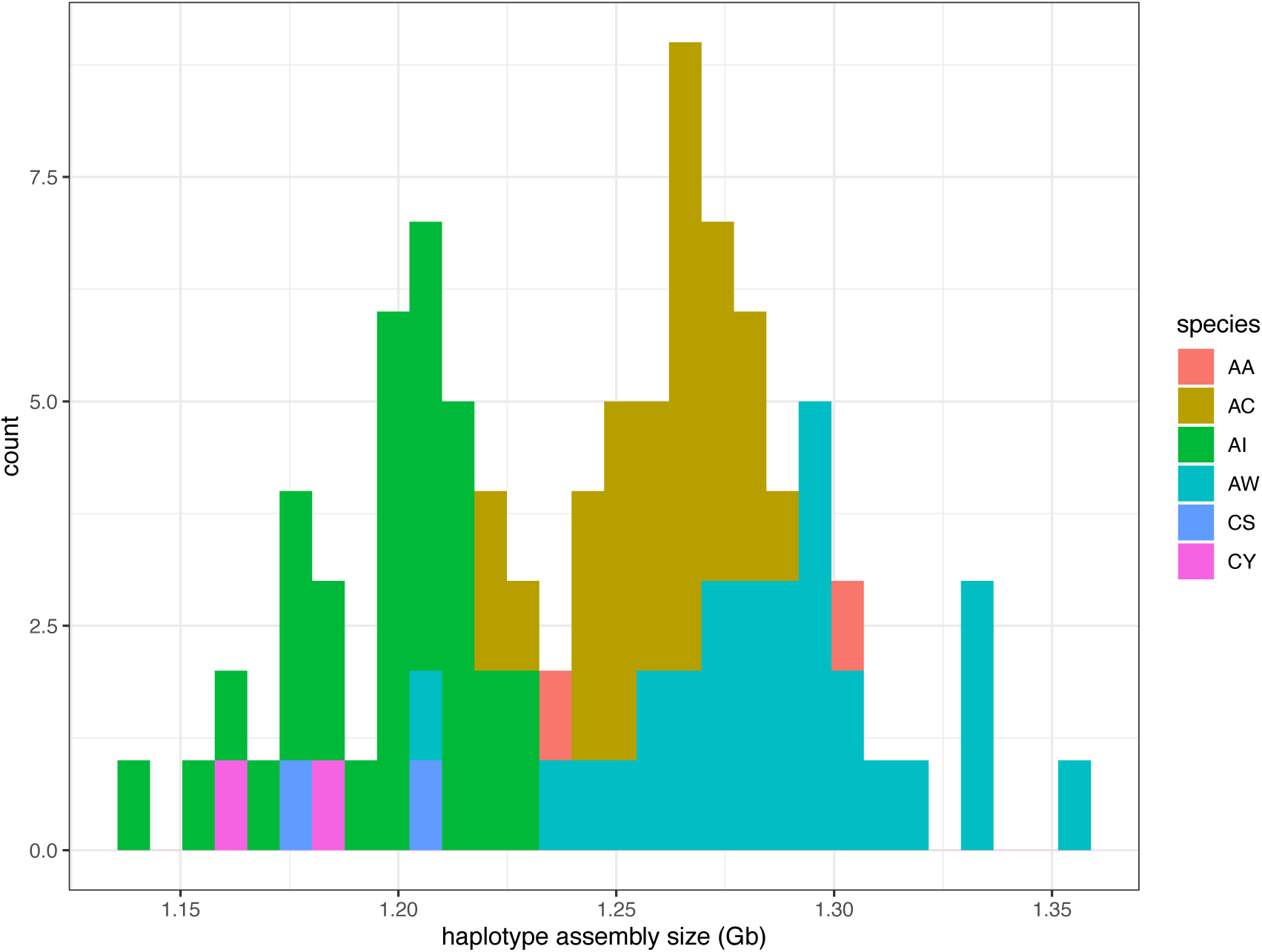
Histogram of assembly sizes (Gb) across 47 *Aphelocoma* individuals. AA=*A. californica*; AC=*A. coerulescens*; AI=*A. insularis*; AW=*A.woodhouseii*; CS=*Cyanocitta stelleri*; CY=*Cyanocorax yucatanicus*.

**Fig. S10.**
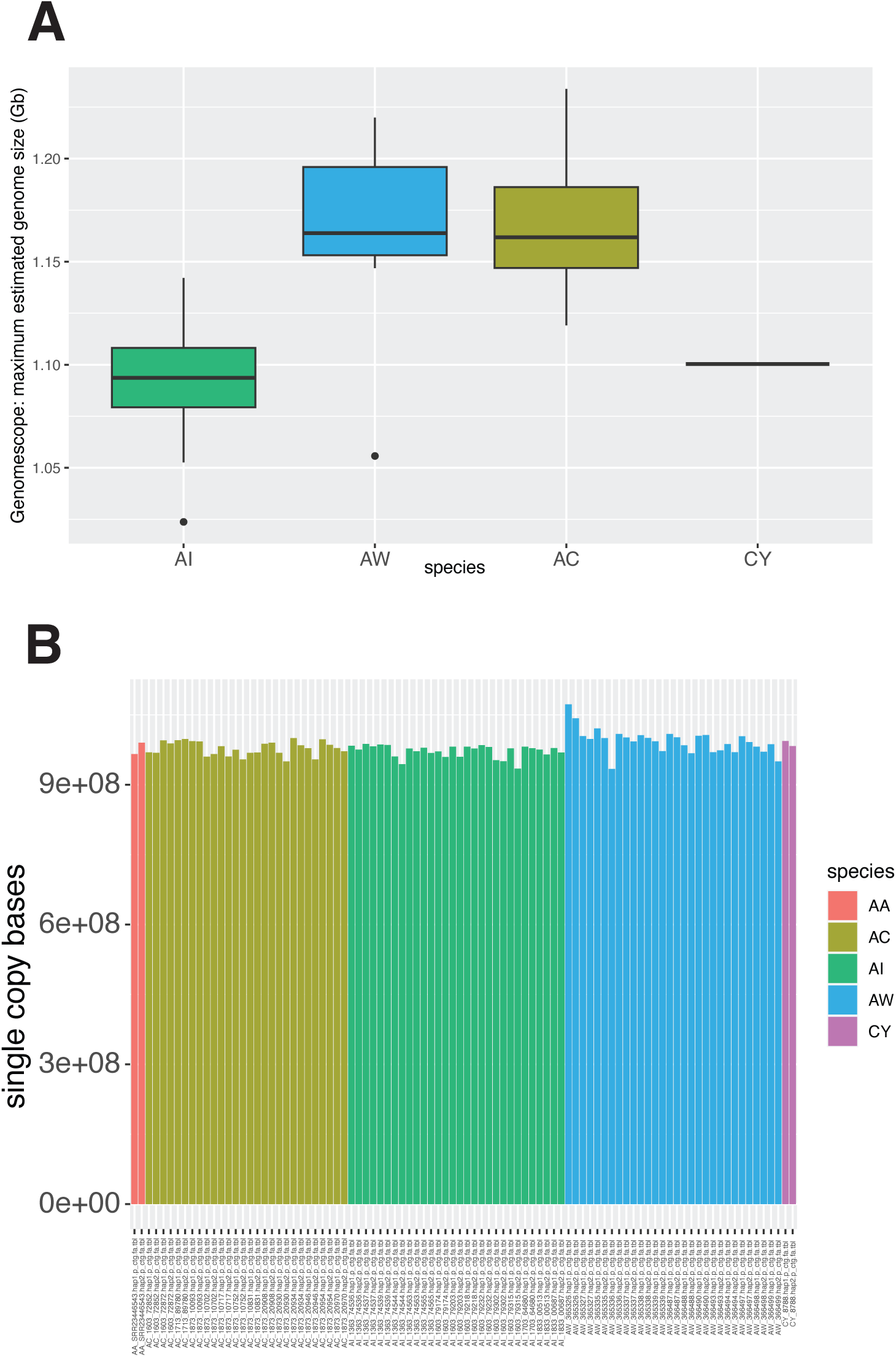
Further details of assembly sizes of focal species. **A,** Estimates of maximum genome size from Genomescope. **B,** number of single copy bases per haplotype from RepeatMasker.

**Fig. S11.**
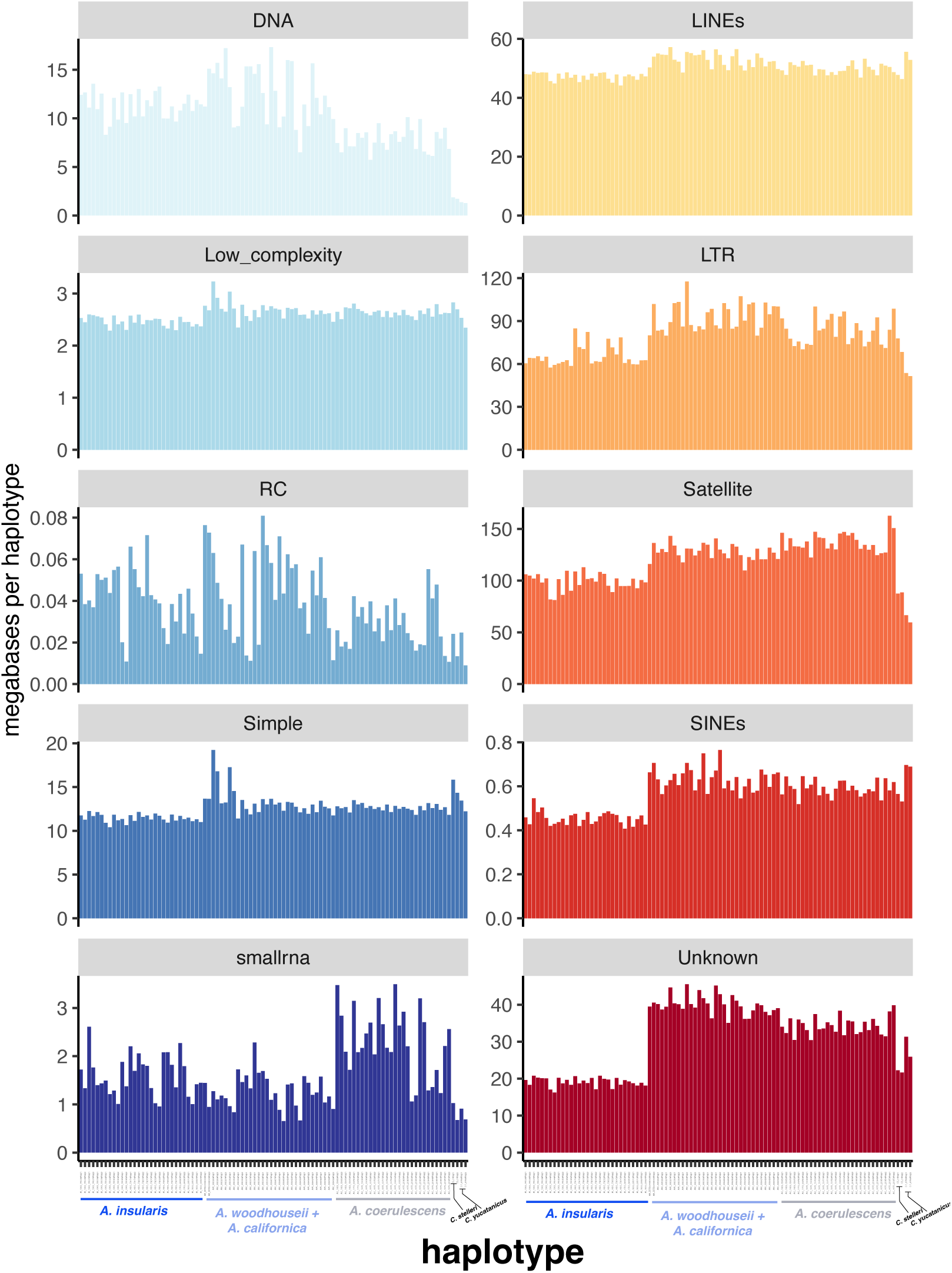
Abundance of different repeat types across haplotypes. Abundances are derived from RepeatMasker analysis of individual haplotypes. Species are indicated at the bottom.

**Fig. S12.**
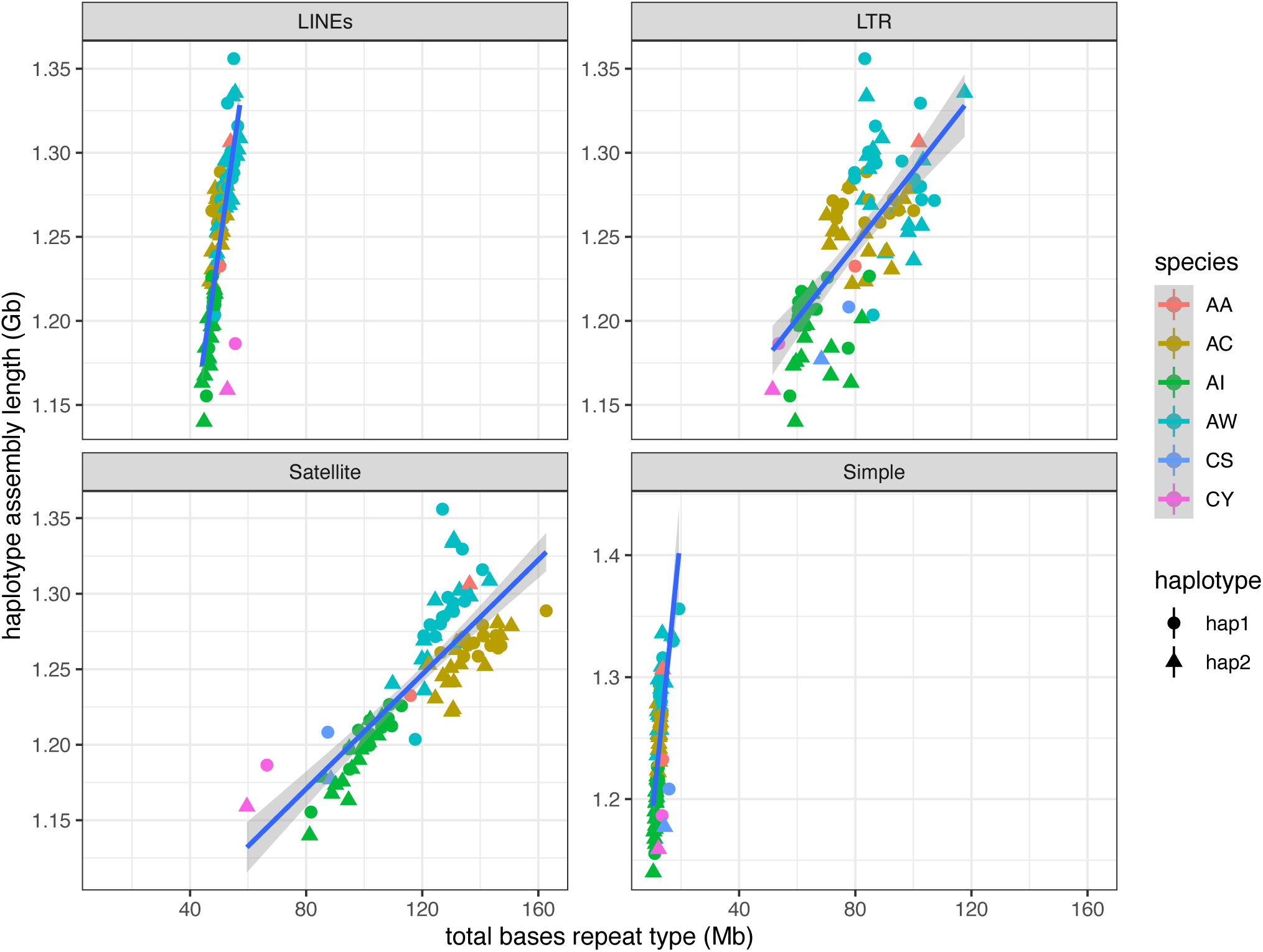
Repeat abundances and assembly sizes. Plots of the four repeat types that are significantly associated with differences in assembly size between haplotypes across species. A linear model was analyzed predicting assembly size from the abundance of 10 repeat types (including unknown; Fig. S11: F = 409, df= 15,78, R^2^ = 0.985, p < 2.2e-16) and species. Satellites (R^2^=0.86), LINEs (R^2^=0.86), LTRs (R^2^=0.71) and simple sequence repeats (R^2^=0.84) were significantly associated, as was Low complexity repeats (not shown, R^2^=0.88).

**Fig. S13.**
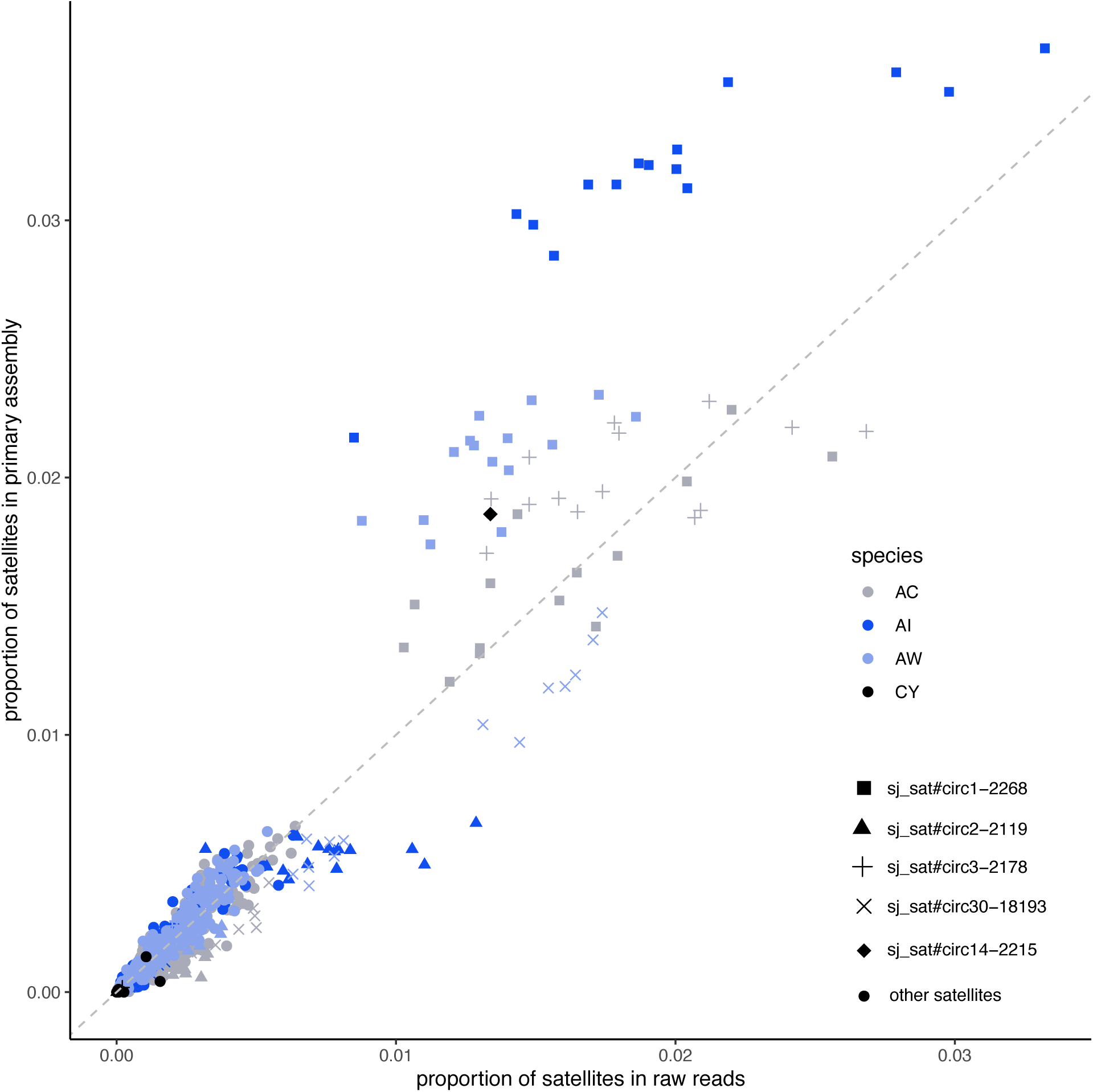
Correlation of proportion of satellites in reads and assemblies. The plot shows the relationship between proportional abundance of different satellites across four species in HiFi reads and in primary assemblies (one per individual). The gray dashed line indicates y=x. The five most common satellites appear in different symbols according to the key at right. The four core species are indicated by different colors. The correlation coefficient between proportions in reads and assemblies is 0.13 (95% c.i. =[0.073, 0.182], t = 4.5565, df = 1256, p-value = 5.707e-06).

**Fig. S14.**
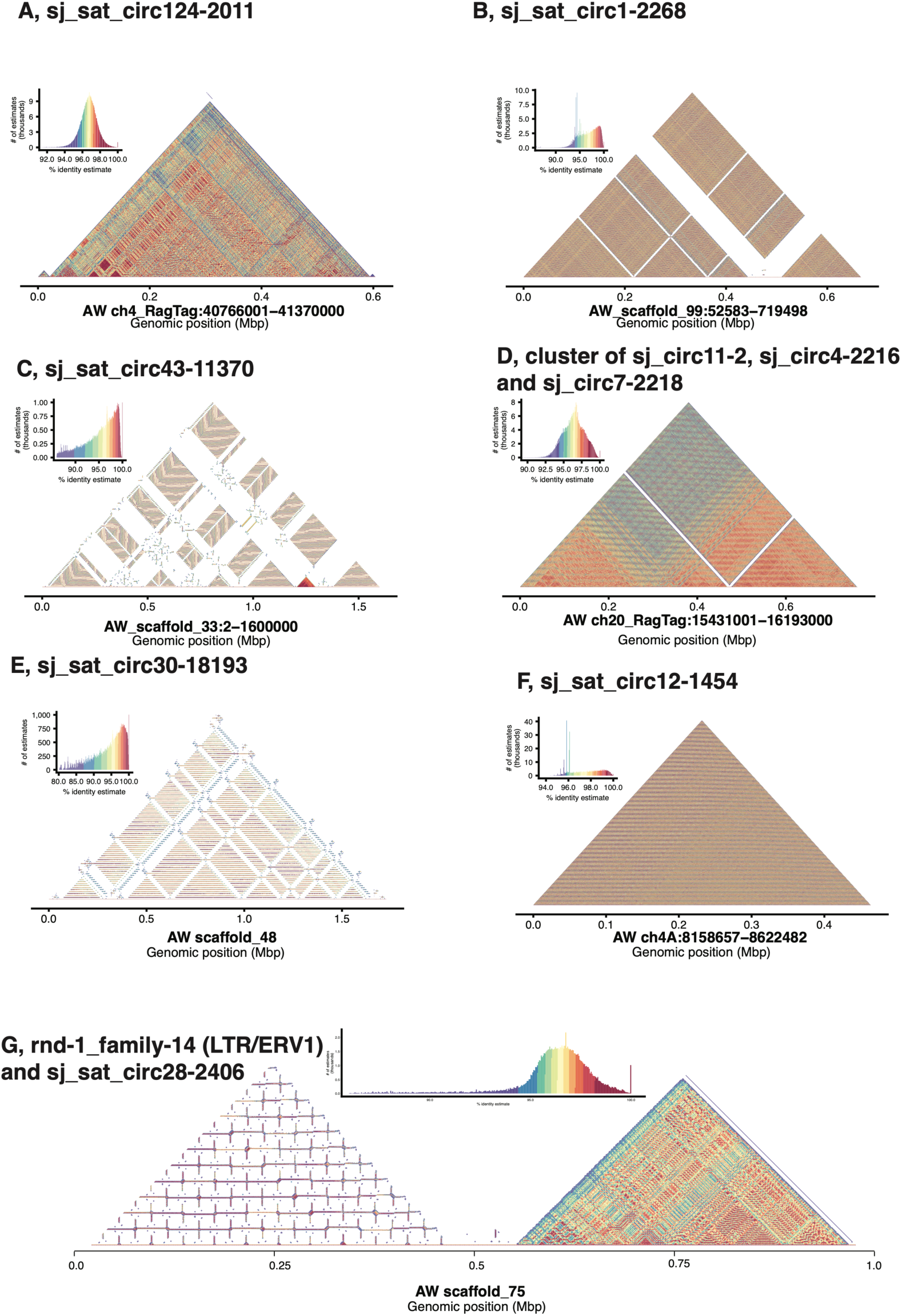
Moddotplots of major satellites. **A-G**, major satellites with abundance key for each plot at upper right. Coordinates in the AW reference and scale in Mbp are indicated below each plot. Plot G shows a SRF satellite as well as a nearby long terminal repeat array.

**Fig. S15.**
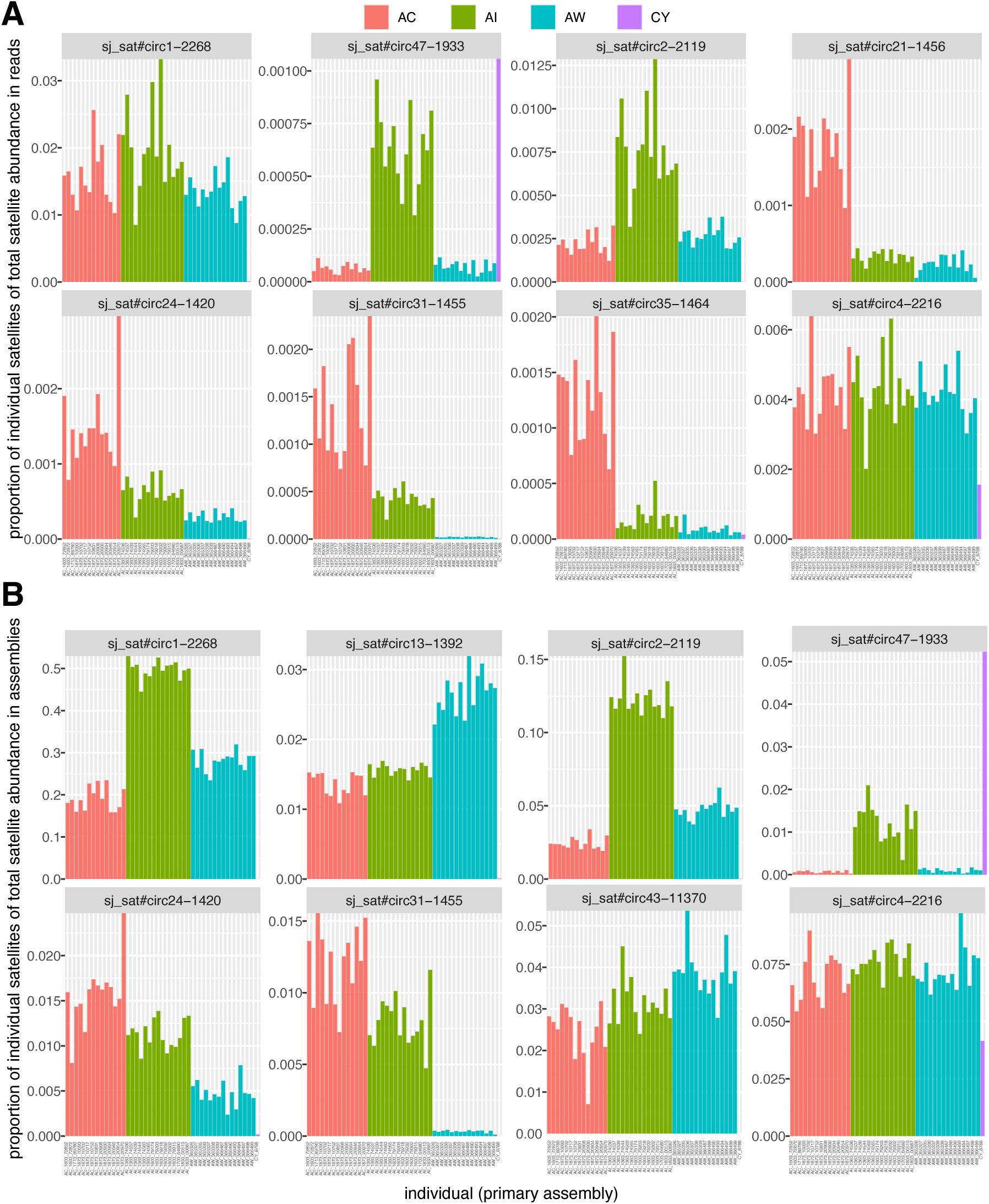
Selected high-abundance satellites in AI birds. These satellites are depicted because they are more common in AI birds than in either AW or AC or both in either HiFi reads (**A**) or in primary assemblies (**B**), despite AI birds having the smallest assemblies.

**Fig. S16.**
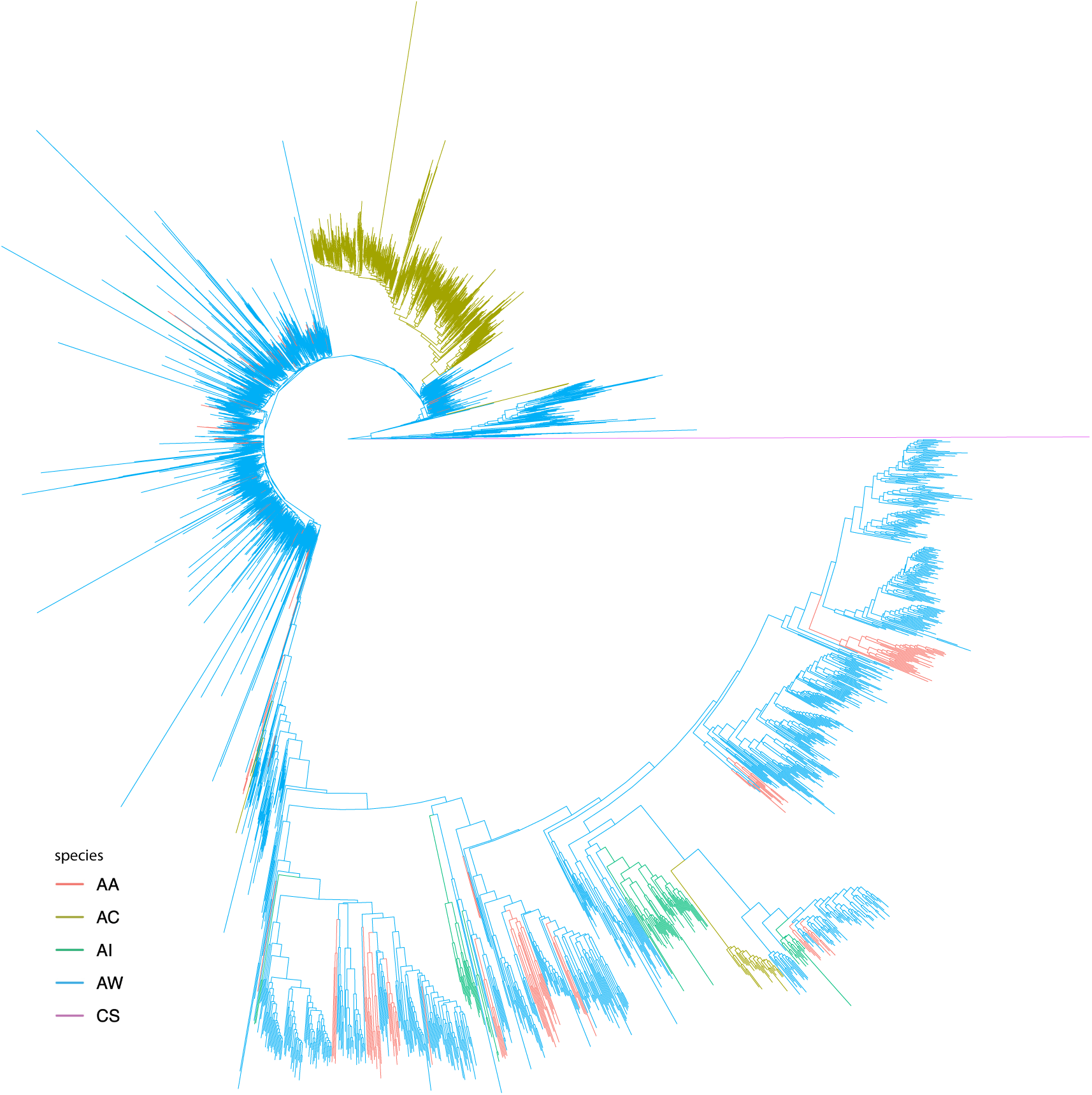
Phylogenetic relationships of ∼27,000 18-kb satellite units. The tree was made with iqtree (Nguyen et al. 2015) from 3,500 exemplars of the original 27,083 units that differed by a minimum of 0.001 substitutions per site, and visualized with ggtree (Yu et al. 2017). See Methods for further detail.

**Fig. S17.**
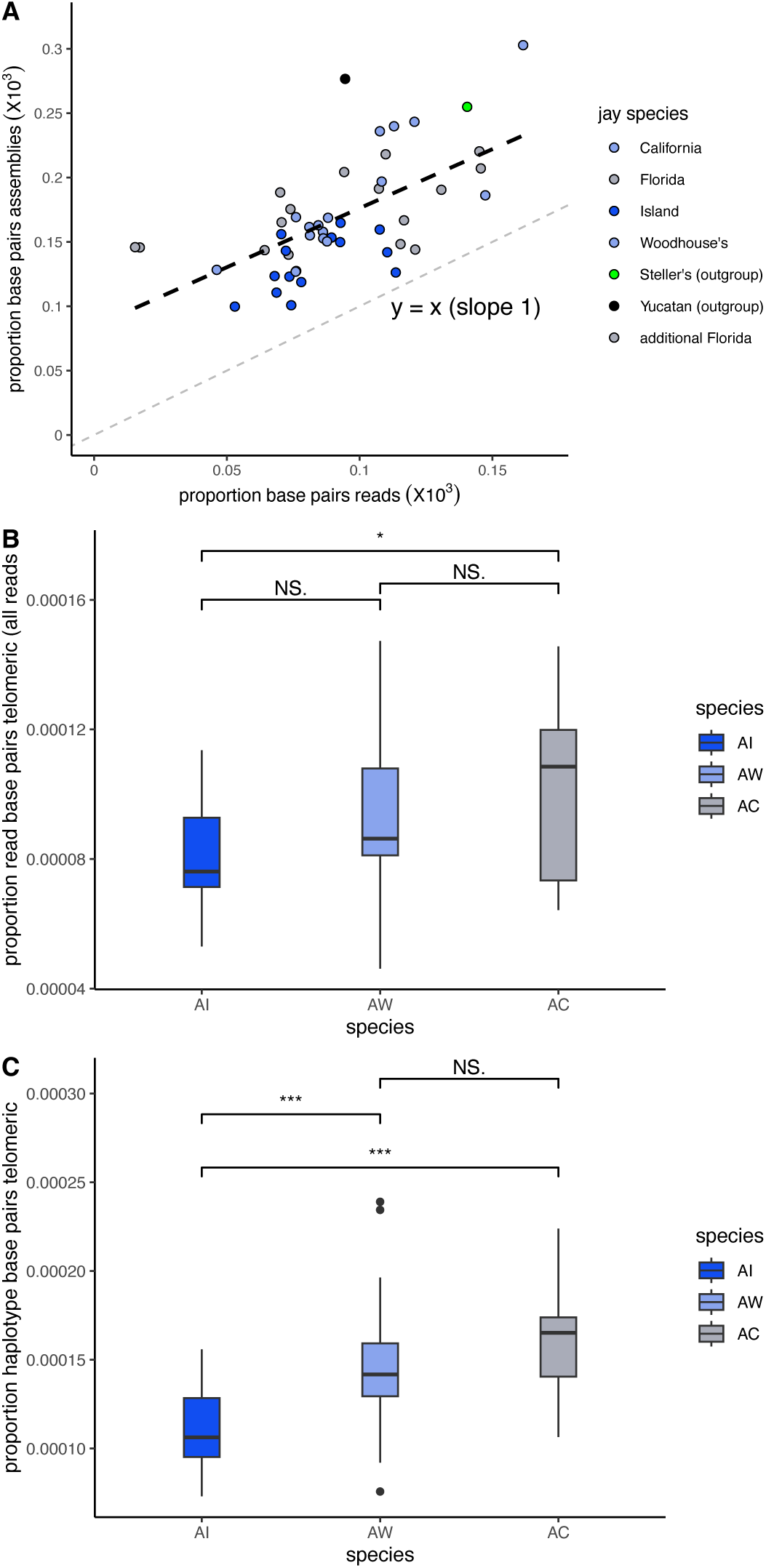
Abundances of telomeric sequences in scrub-jays and relatives. **A**, correlation between telomere abundances in HiFi reads and in individuals (sum of two haplotype assemblies per individual). The dashed line shows y=x. The data has a slope of 0.917 (F = 30.04, R^2^ = 0.3769, p = 1.628e-06). **B**, boxplot of proportion of telomeric sequence in HiFi reads among individuals of AI, AW and AC. **C**, boxplot of proportion of telomeric sequence in assembled haplotypes among individuals of AI, AW and AC

**Fig. S18.**
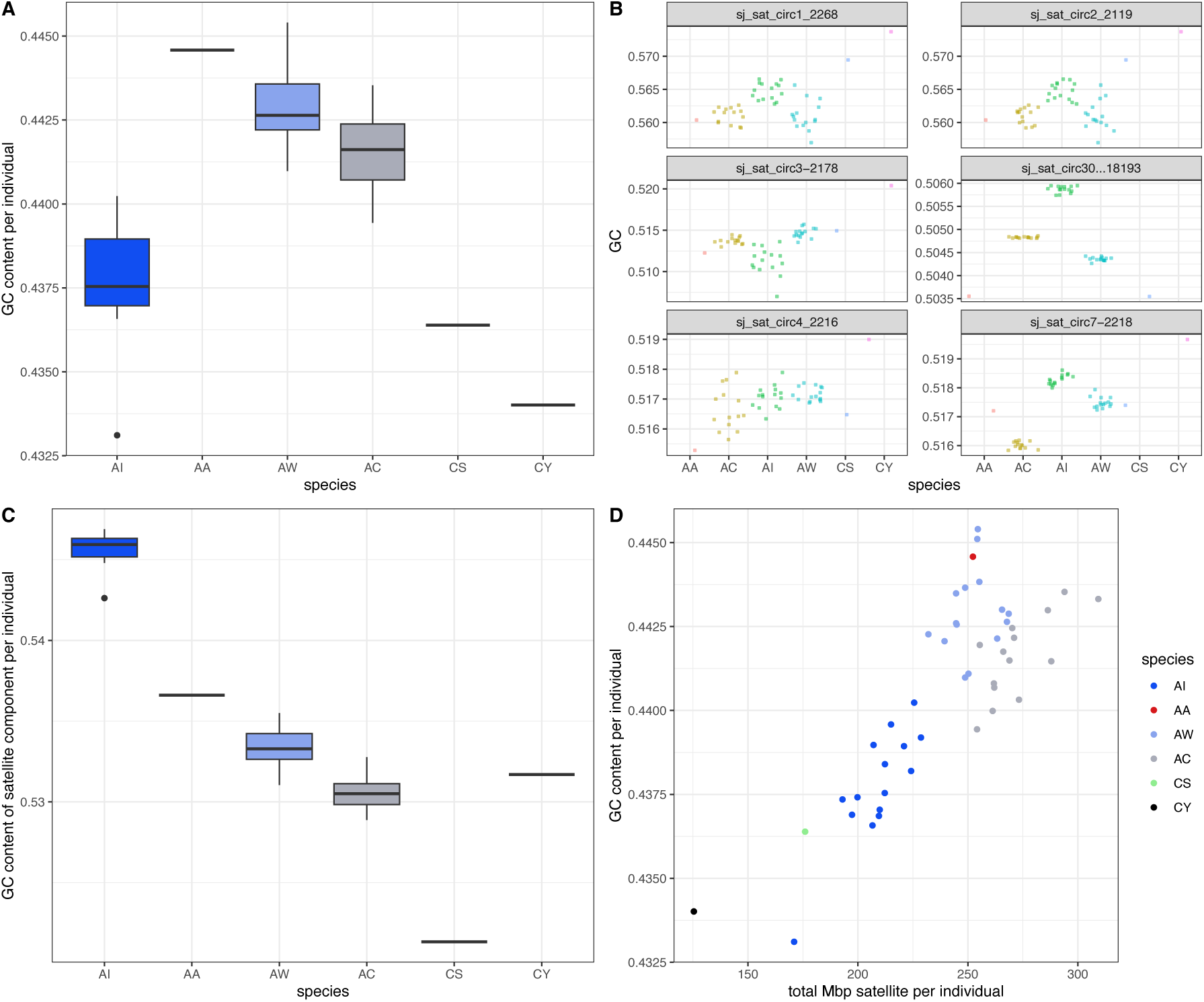
Effects of satellite GC content on genome-wide GC content. **A**, Genome-wide GC content for species in the study. Species codes are as in fig. S3. **B**, GC content of the six most common satellites identified by Satellite Repeat Finder (Zhang et al. 2023). **C**, GC content of entire satellite component of each individual. **D**, correlation of total satellite component per individual and genome-wide GC content. A linear model explaining genome-wide GC content and including five satellites (excluding sj_sat_circ2_2119, which was redundant) had R^2^ = 0.63, p = 1.54e-08, with significant or near effects of sj_sat_circ1_2268, p = 0.024, *β* = −4.08e-09; sj_sat_circ30_18193, p = 0.66, *β*= 1.29e-10; sj_sat_circ7_2218, p = 0.057, *β*= 2.03e−10; and the non-satellite genome component, p = 8.52e-06, *β* = 2.09e-11.

**Fig. S19.**
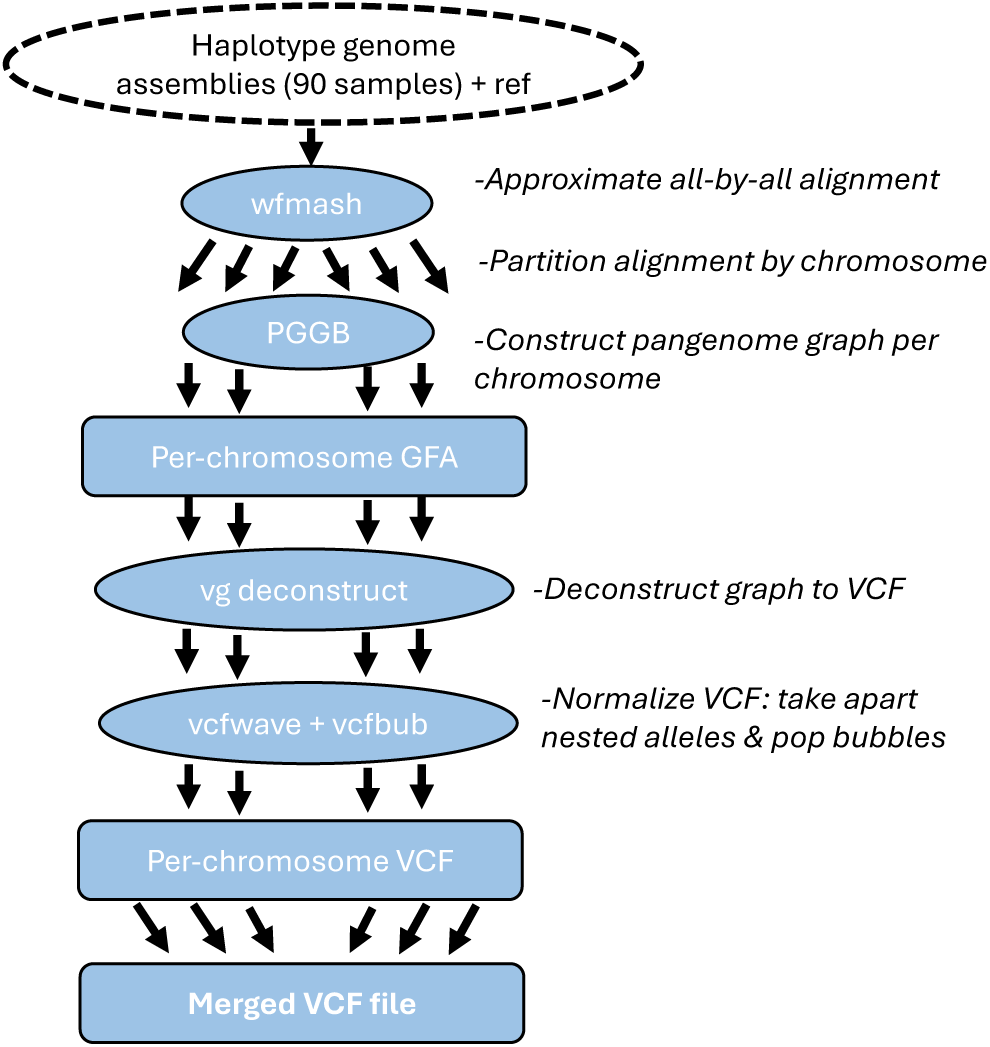
Pipeline for creating the merged VCF file from PGGB pangenome graph. The pipeline begins with the 90 phased haplotypes from AW, AC, AI and CY and uses wfmash, PGGB, vcfwave and vcfbub to generate per-chromosome VCFs, followed by merging these with bcftools (Danecek et al. 2021).

**Fig. S20.**
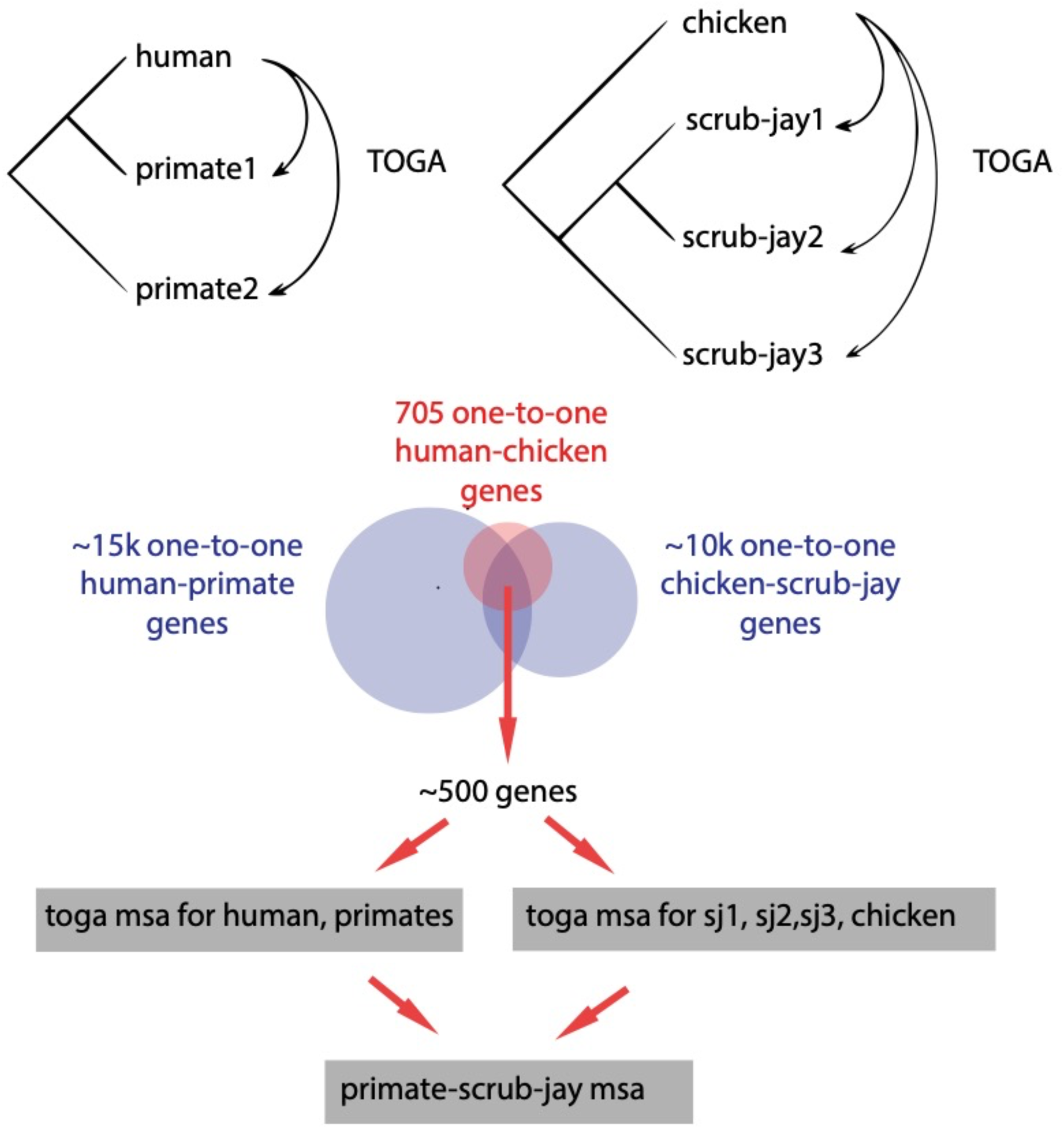
Pipeline for generating jay and primate 1:1 ortholog alignments. TOGA (Kirilenko et al. 2023) was used to generate human-primate alignments and jay-chicken alignments. An existing alignment of human-chicken allowed finding genes in common between the two groups and merging the multiple sequence alignments (MSA) for each group separately and to each other.

**Fig. S21.**
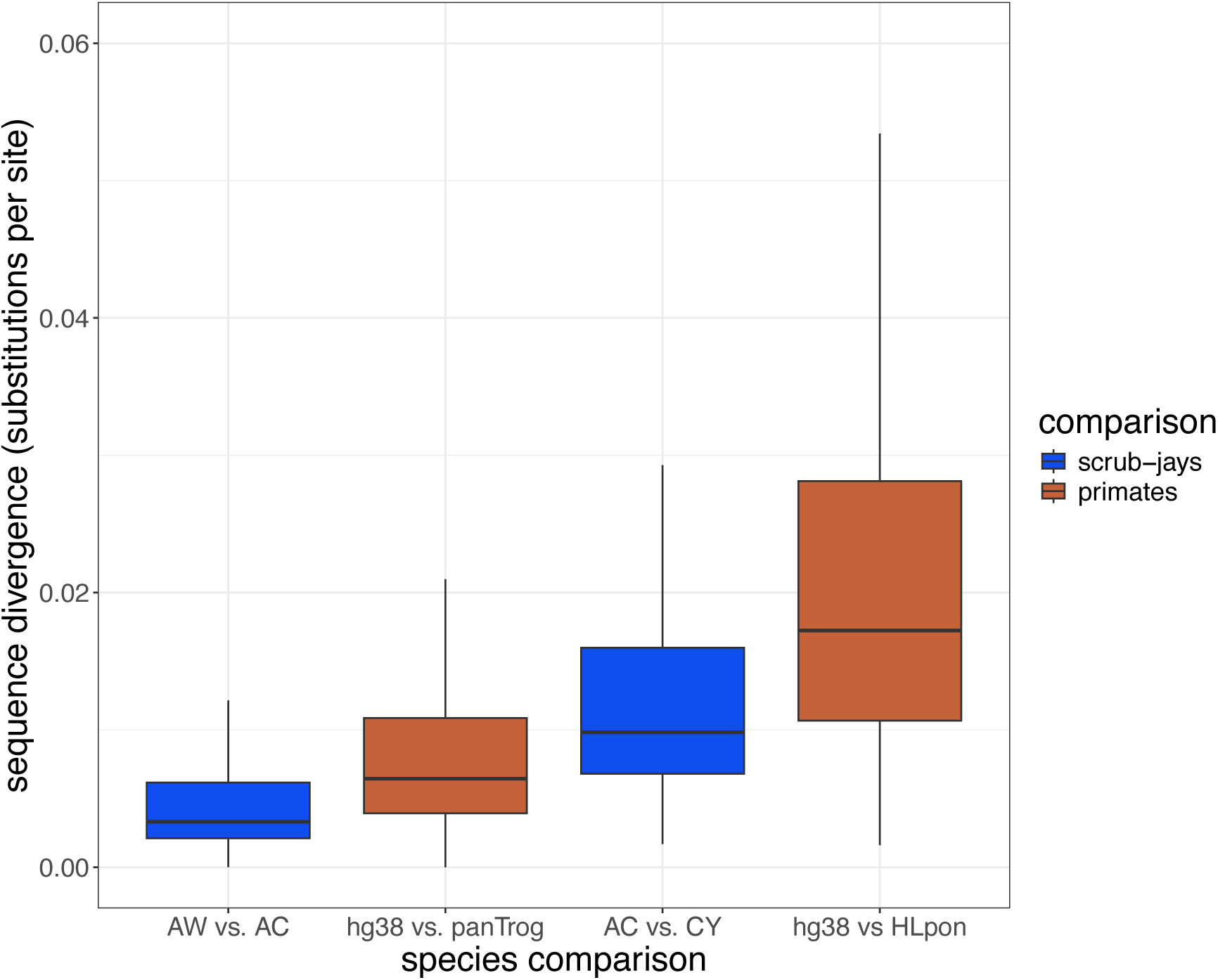
Comparison of sequence divergence between scrub-jays and primates. Alignments of ∼400 orthologous genes were analyzed with ape (Paradis et al. 2004) to estimate sequence divergence between species. Species codes are as in fig. S3. hg38 is the human genome assembly GRCh38 (NCBI GCF_000001405.40); panTrog is the Chimpanzee (*Pan troglodytes*, GCA_000001515.5); HLpon = Sumatran orangutan, Pongo abelii (GCA_015021835.1).

**Fig. S22.**
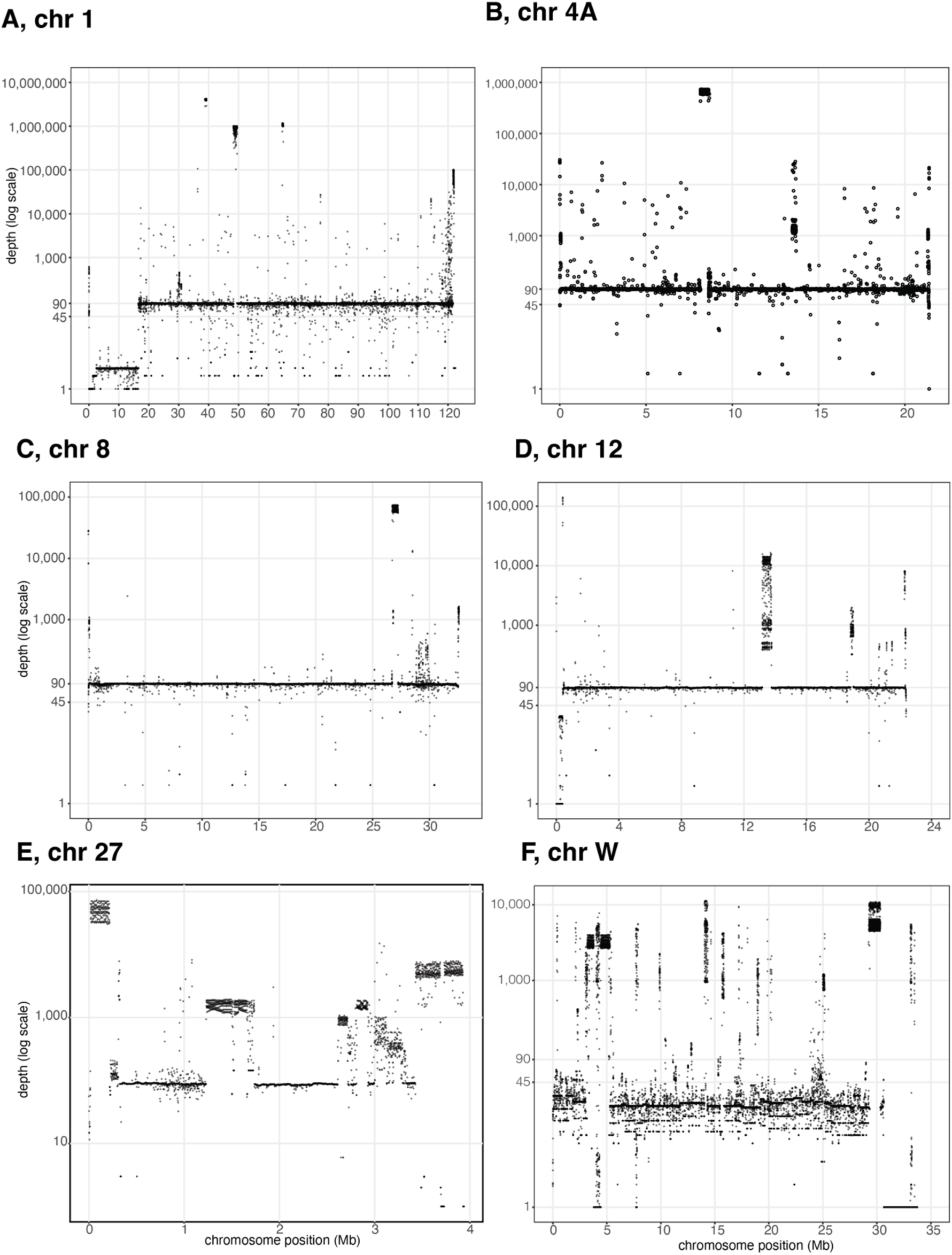
Examples of PGGB pangenome graph depth across whole chromosomes. **A-E**, chromosomes are indicated above each plot. Chromosome positions are indicated at the bottom of each plot. Y-axis is on a log-scale.

**Fig. S23.**
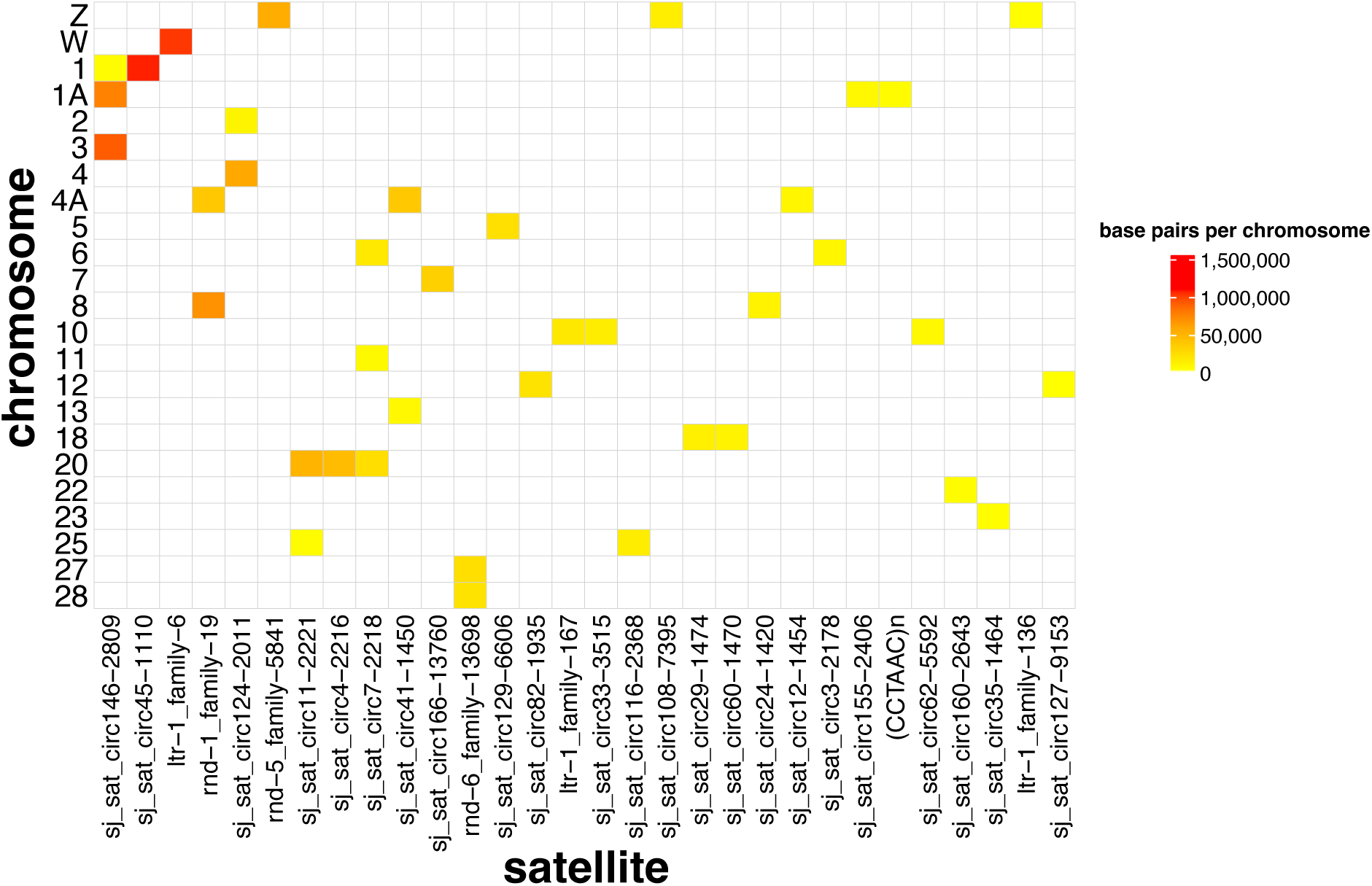
Distribution of satellites in highest-depth chromosomal regions. Bedfiles were produced of the highest-depth regions of each chromosome (generally depth > 1000; one region per chromosome). These bed files were intersected with the RepeatMasker annotation for the AW reference assembly and their abundance plotted using the R package ComplexHeatmap (Gu 2022).

**Fig. S24.**
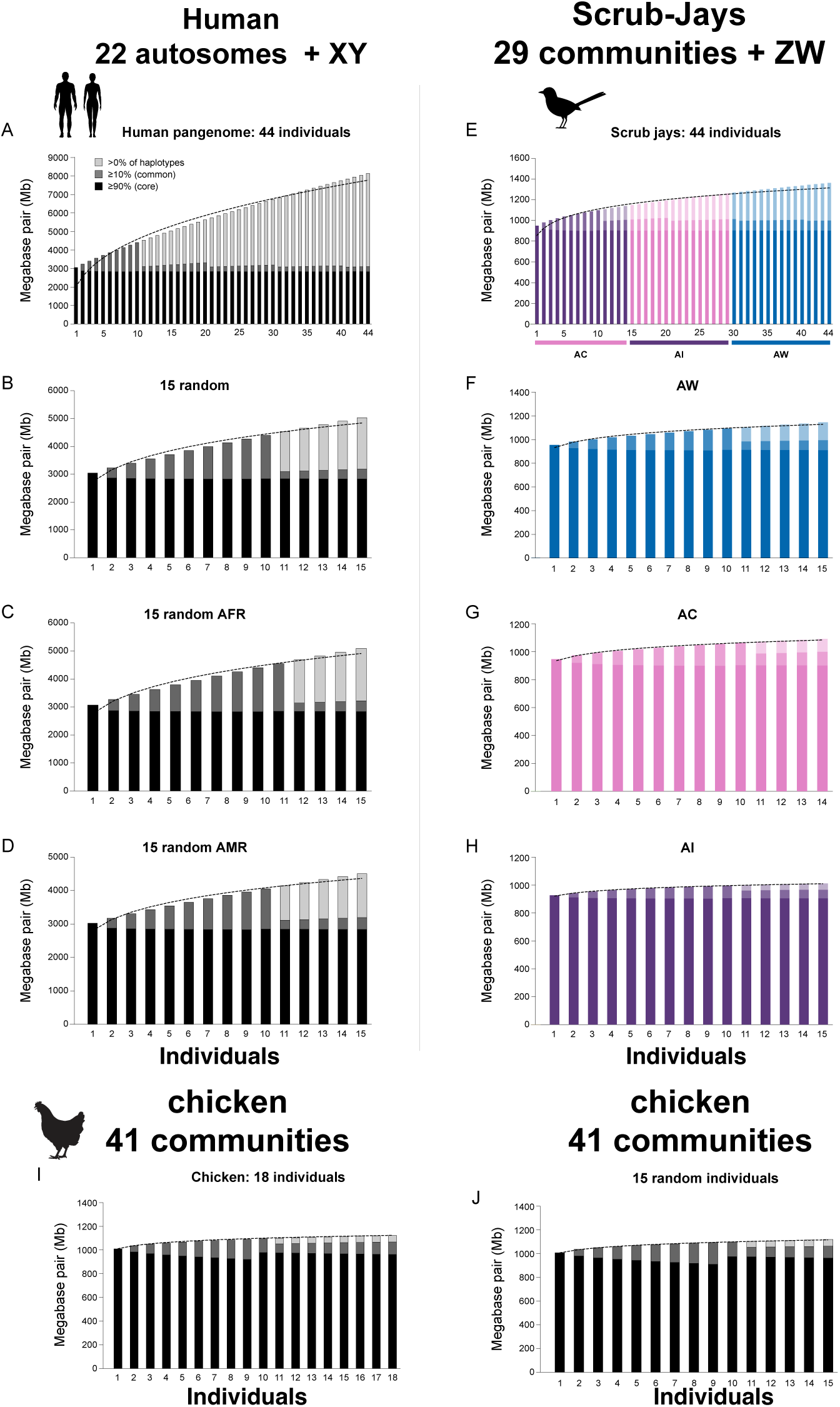
Comparison of pangenome Panacus plots of scrub-jay, human and chicken. PGGB pangenome graph growth curves for A, 44 individuals (88 haplotypes). B, 15 random human individuals. C, 15 random African individuals. D, 15 random American individuals. E, 44 Scrub-Jay individuals (88 haplotypes across AC, AI and AW). F, 15 AW individuals. G, 14 AC individuals. H, 15 AI individuals. I, 18 chickens. J, 15 chickens. Human data from (Liao et al. 2023) and chicken data from (Rice et al. 2023).

**Fig. S25.**
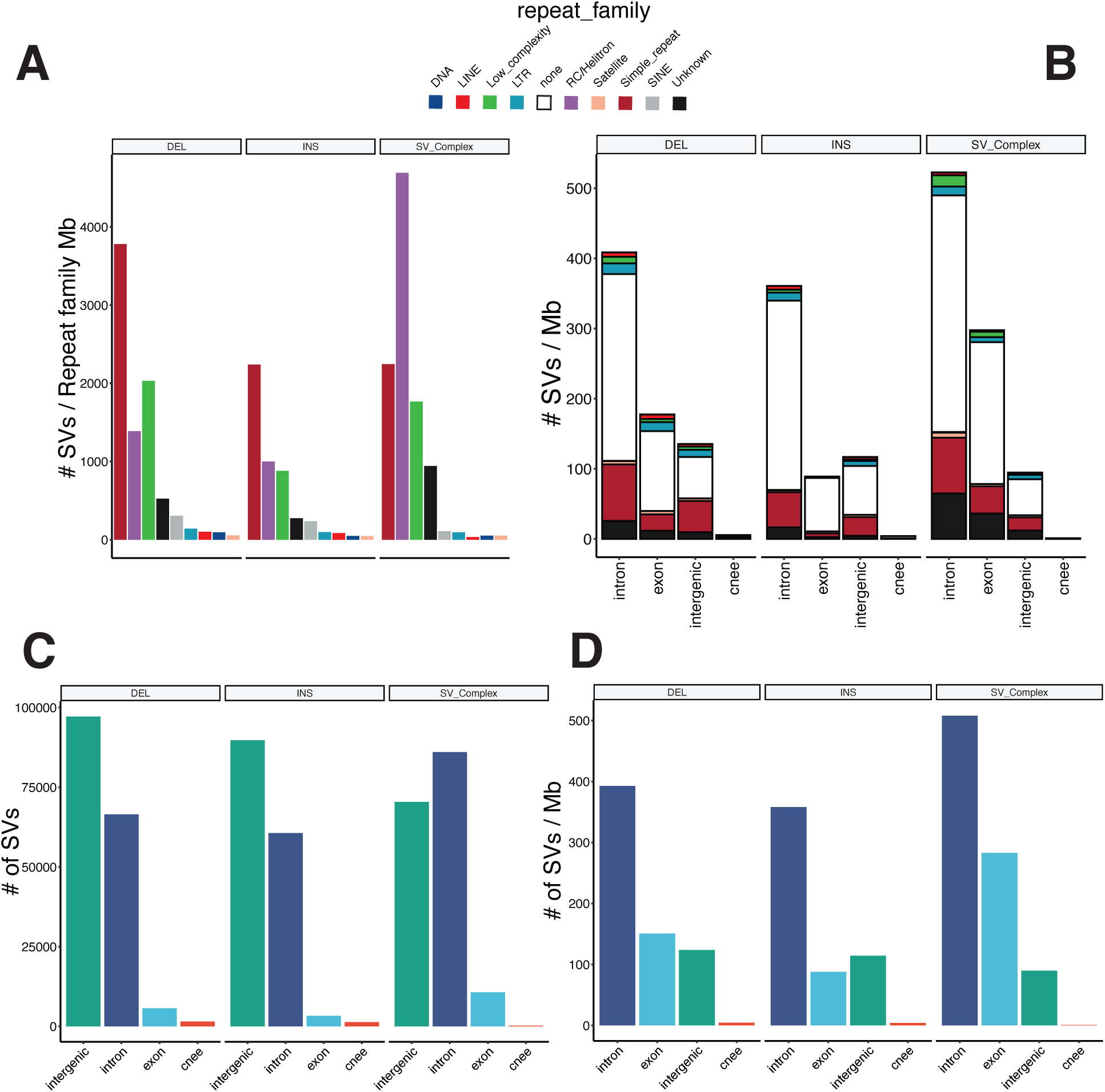
Overlap of structural variants, repeat types and annotations. **A**, Number of SVs (deletions, insertions and complex SVs) per repeat family as annotated by RepeatMasker. **B**, density (SVs per Mb) of deletions, insertions and complex SVs among repeat types. **C**, Number of SVs in different genomic annotations. **D**, Density of SVs in different genomic annotations.

**Fig. S26.**
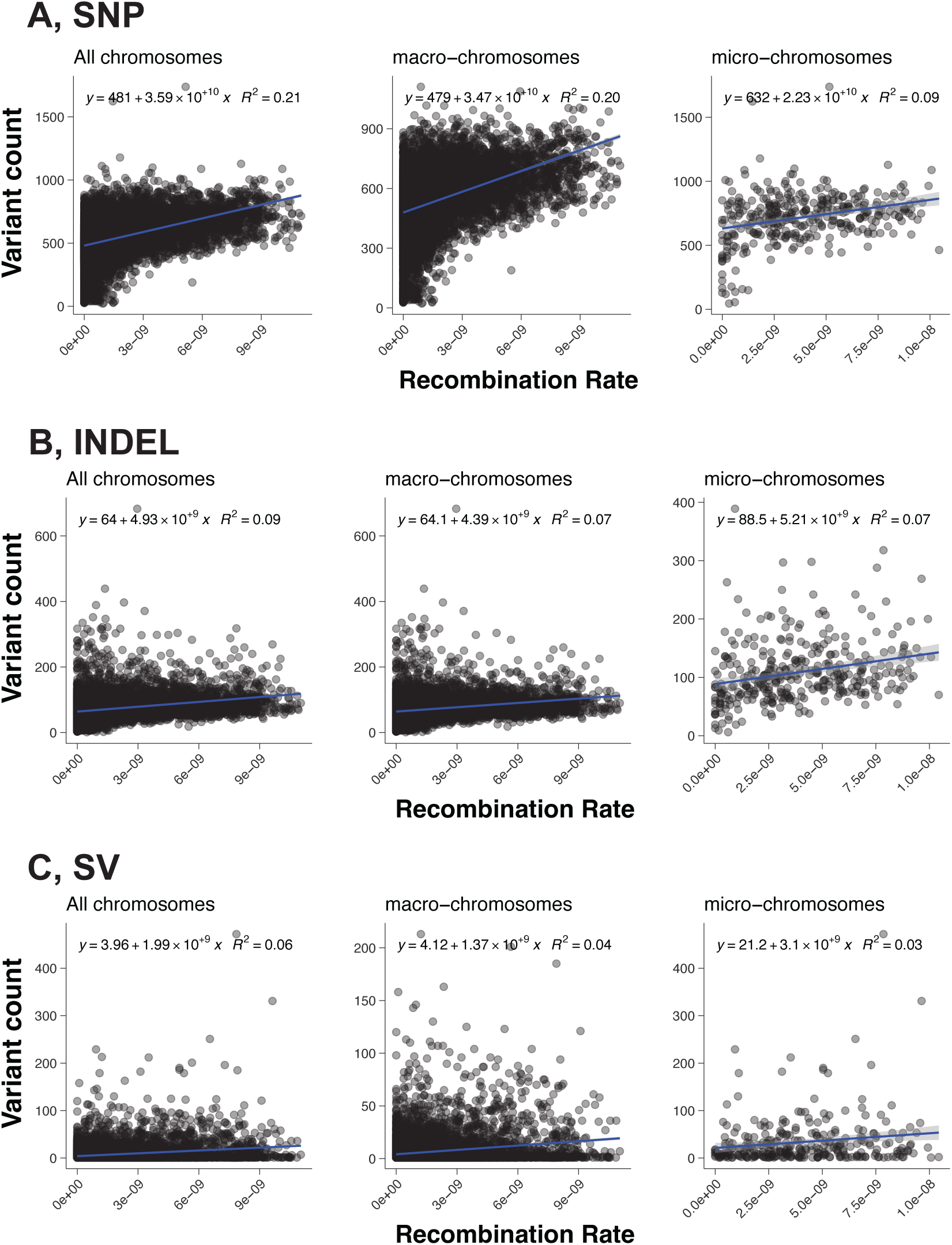
Association of SNP, indel and SV density with recombination rate. **A**, SNPs. **B**, indels. **C**, SVs. On each plot the y-axis is the count of variants in 1 Mb windows (from PGGB VCF). The x-axis is the average recombination rate in 1 Mb windows as estimated by RELERNN (Adrion et al. 2020). Left to right is all chromosomes, macro-chromosomes and micro-chromosomes. Equations relating variables are indicated on each plot.

**Fig. S27.**
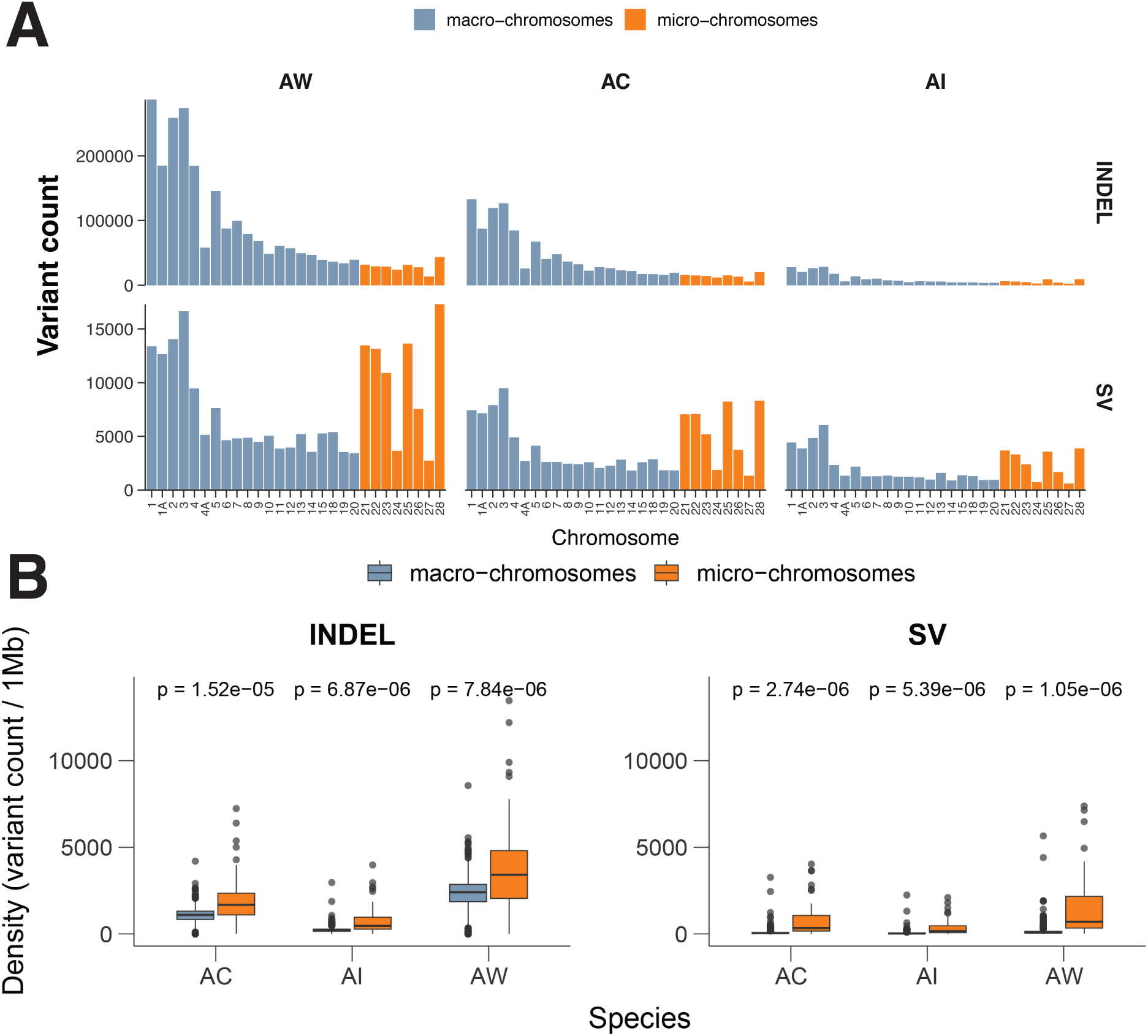
Number and density of SVs on micro- and macrochromosomes. A, number of indels and SVs across individuals within AC, AI and AW. B, Density of indels and SVs for each species between micro- and macro-chromosomes.

**Fig. S28.**
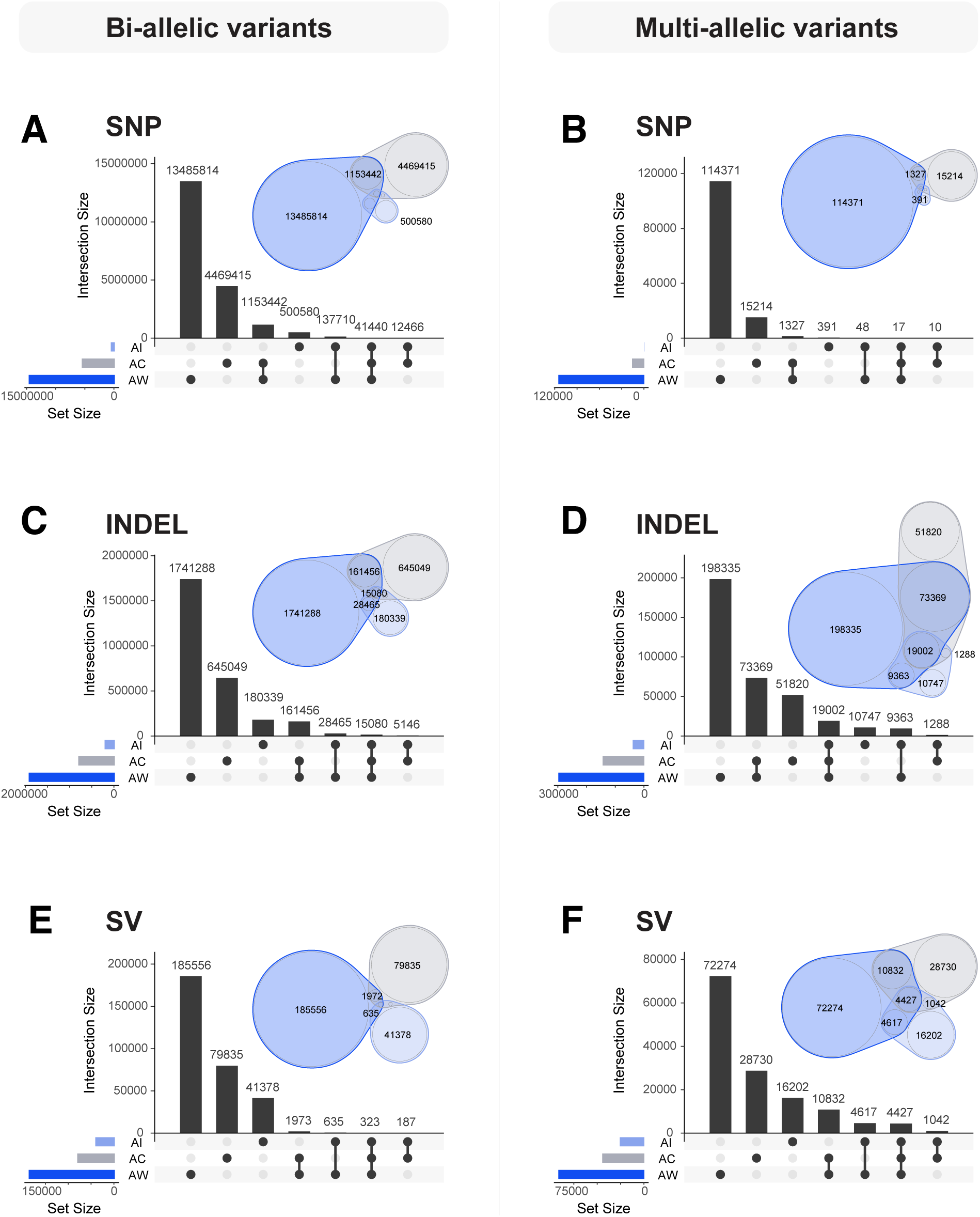
Patterns of sharing and species-specificity of SNPs, indels, and SVs. **A**, **C**, and **E** - biallelic SNPs, indels, and SVs, respectively. **B**, **D** and **F** - multiallelic SNPs, indels, and SVs. Each panel shows a plot of pairwise sharing of variants between species, as well as a Venn diagram (Quesada 2018).

**Fig. S29.**
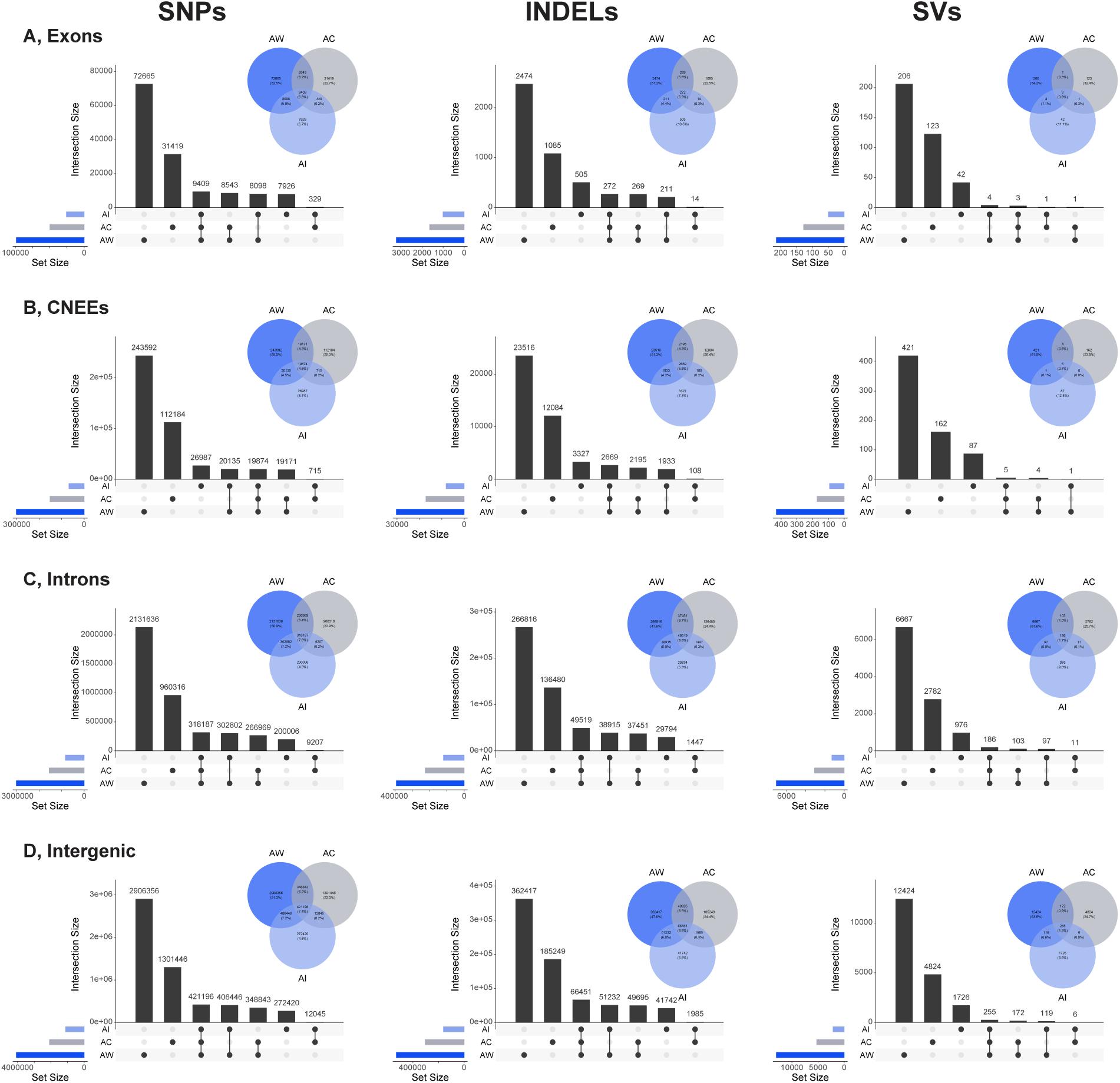
Patterns of interspecific sharing of SVs across genome compartments. Columns, left to right, indicate shared patterns between species in SNPs, indels and SVs. A, sharing in exonic variants. B, sharing in CNEEs. C, sharing in introns. D, sharing in intergenic regions. Each panel shows a Venn diagram as well as barplots of intersection sizes of lineage-specific variants, and variants shared among pairs and trios of species.

**Fig. S30.**
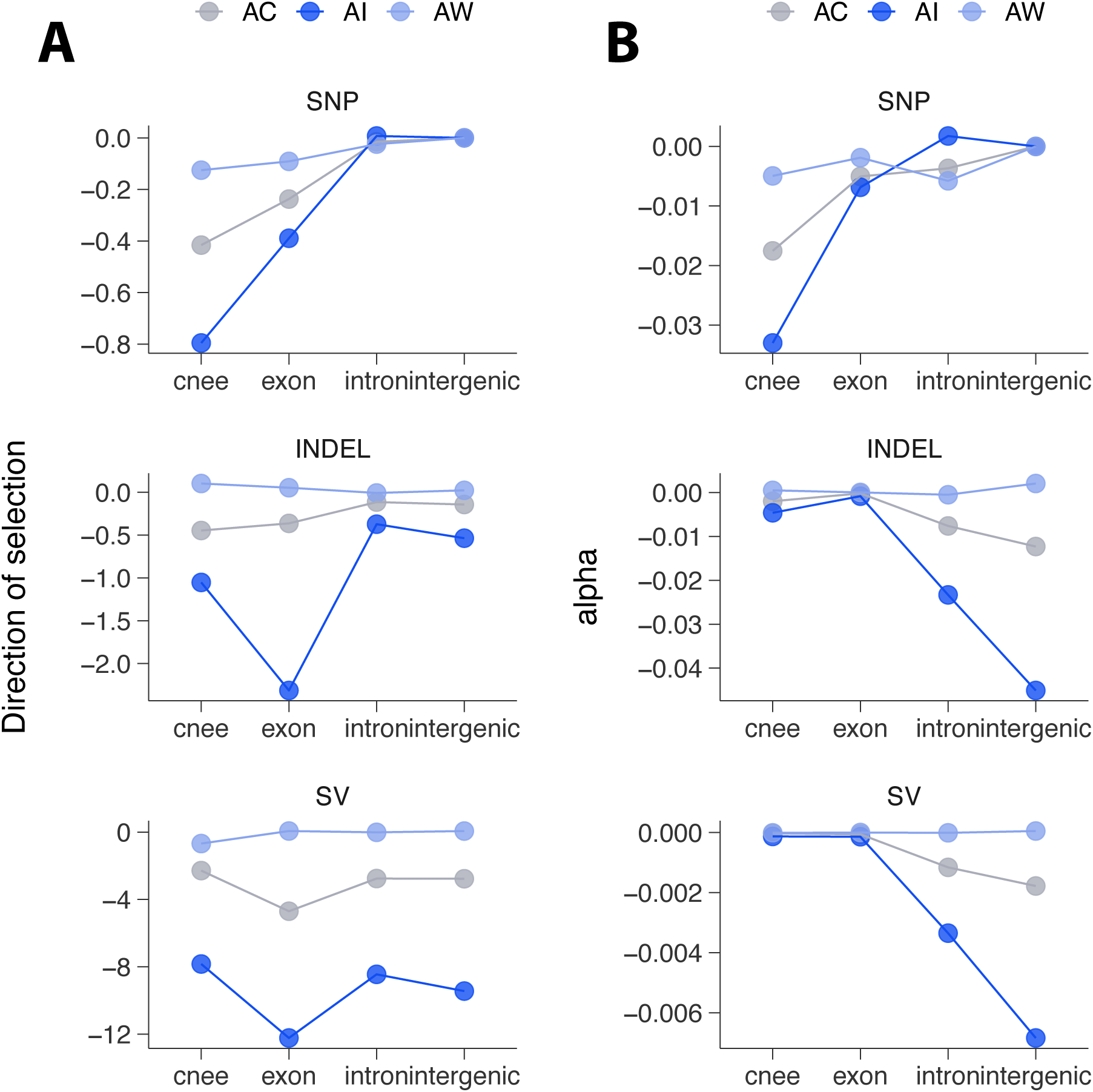
Estimates of the DoS and alpha in different genomic compartments. **A**, Direction of selection (DoS) estimated from the site frequency spectrum of SVs from the PGGB pangenome VCF. **B**, estimates of *α*, the fraction of SVs fixed by selection.

**Fig. S31.**
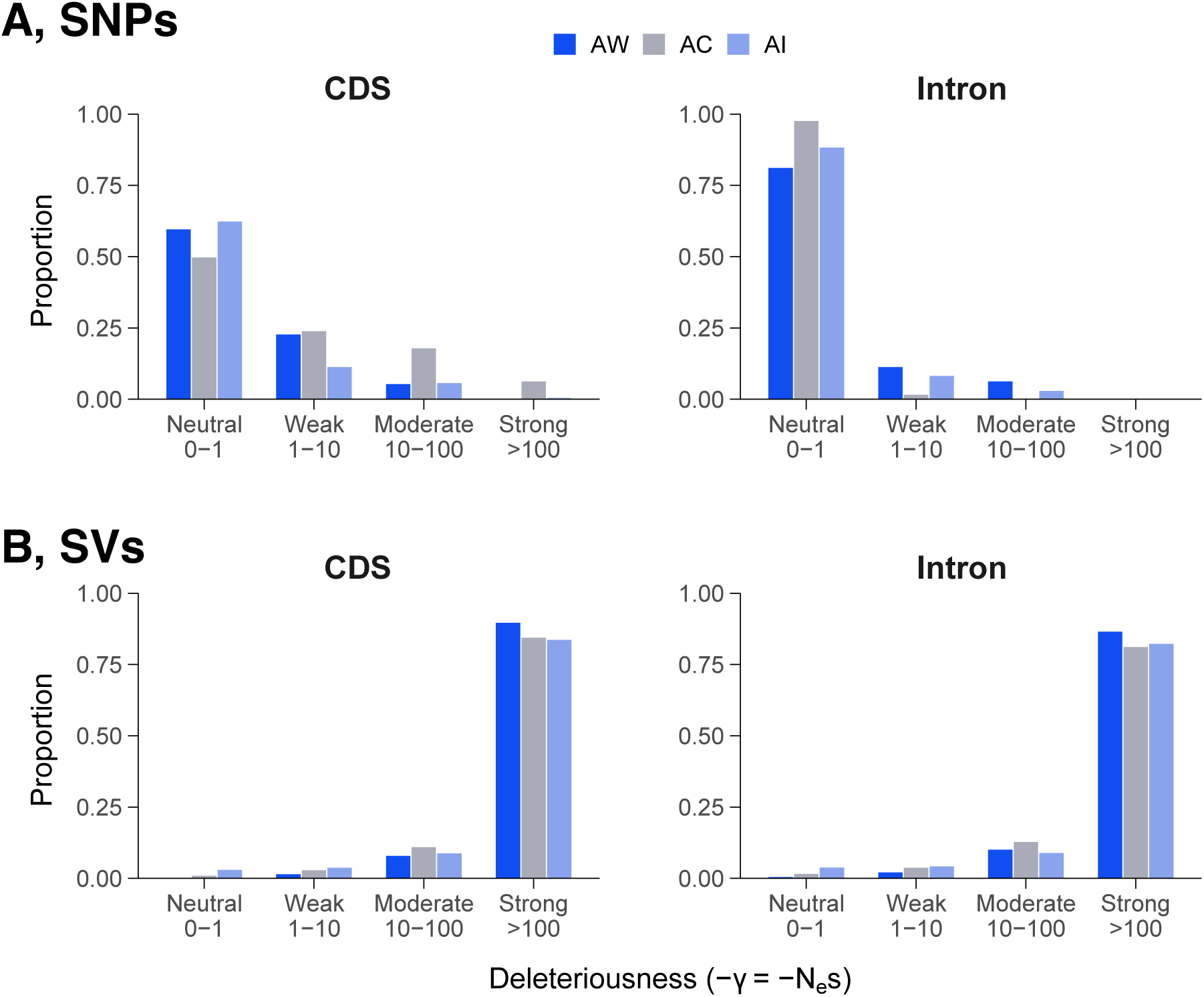
Estimates of the DFE from anavar. **A**, Binned DFE of SNPs from reach species from anavar (Barton and Zeng 2018). **B**, binned DFE of SVs. Coding regions and introns are in the left and right columns, respectively. Intergenic regions are not included because the putatively neutral category of SNPs comes from this subgenome.

**Fig. S32.**
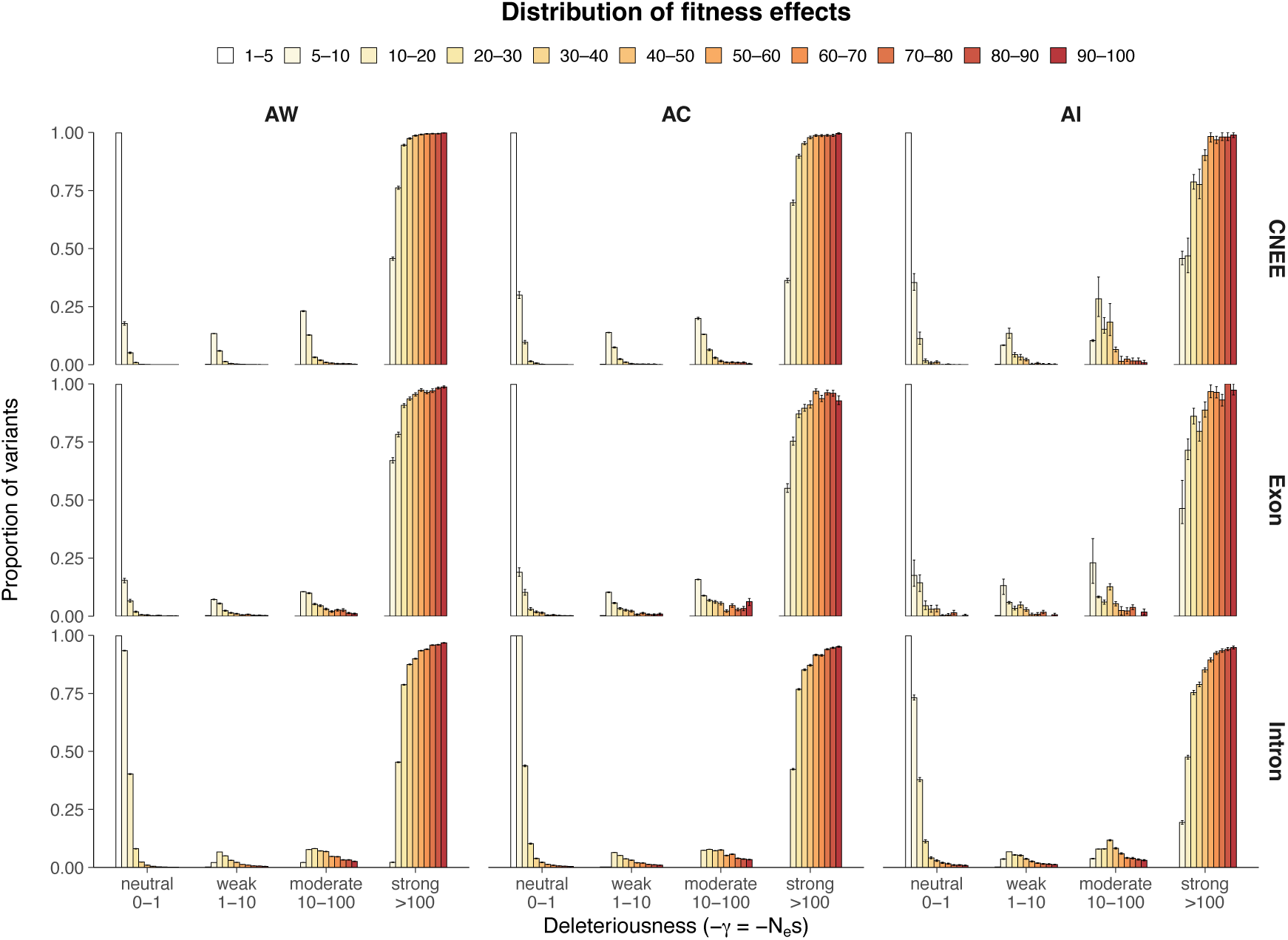
Changes in the distribution of fitness effects with SV length. The plot shows different distributions of the DFE when estimated with different subpopulations of SVs differing in length. The key for SV length in bp is at the top. Estimates of the DFE come from fastDFE (Sendrowski and Bataillon 2024).

**Fig. S33.**
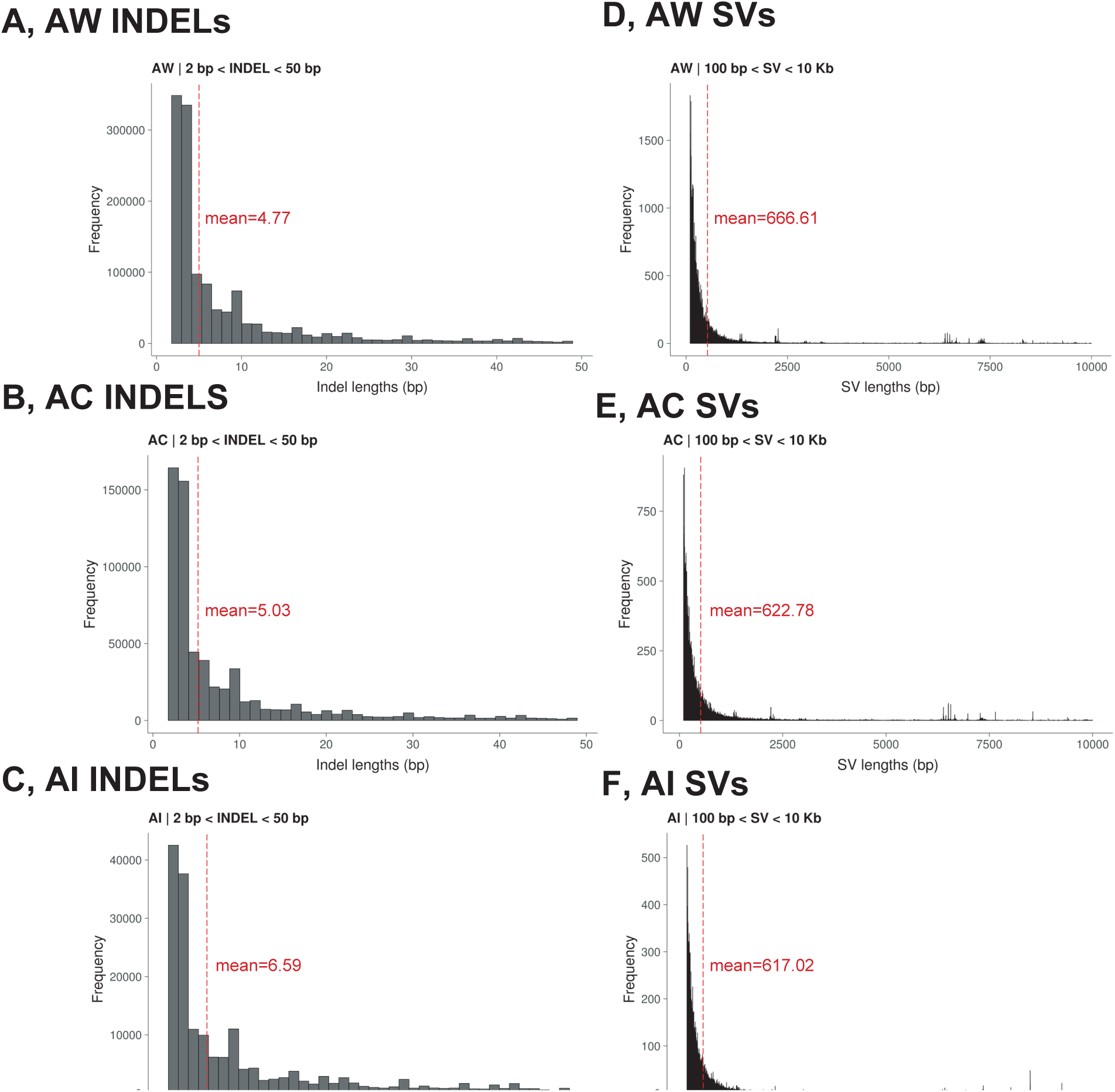
Distribution of lengths of indels and SVs among species. A-C, indel length distribution in AW, AC, and AI, respectively, from the PGGB VCF. D, E, F, SV length distribution in AW, AC, and AI, respectively. Data includes all SVs except for SV_complex.

**Fig. S34.**
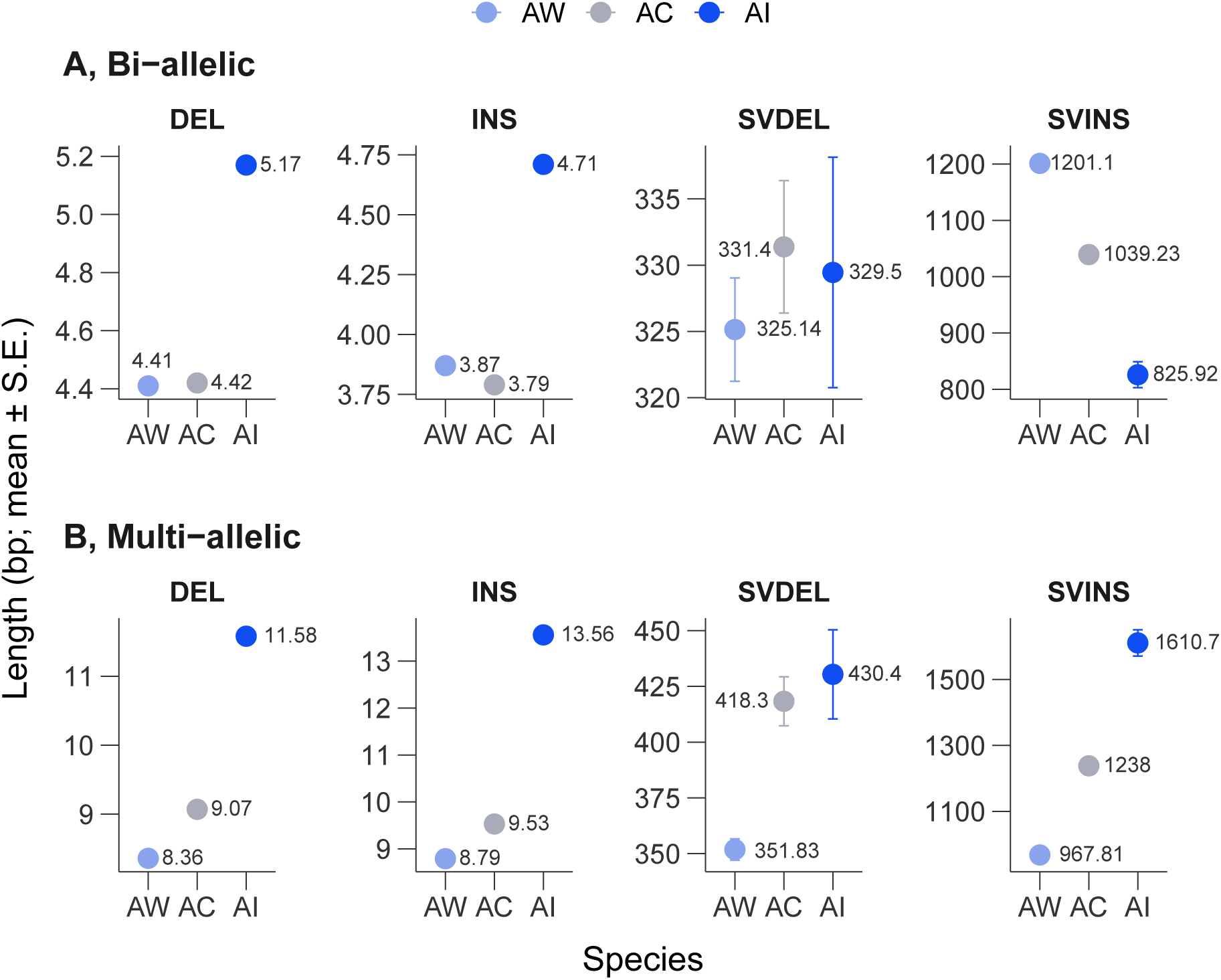
Mean lengths of biallelic and multiallelic indels and SVs between species. **A**, biallelic SVs, from the PGGB VCF. **B**, multiallelic SVs. Indels and SVs were polarized with the CY outgroup from the main PGGB pangenome graph.

**Fig. S35.**
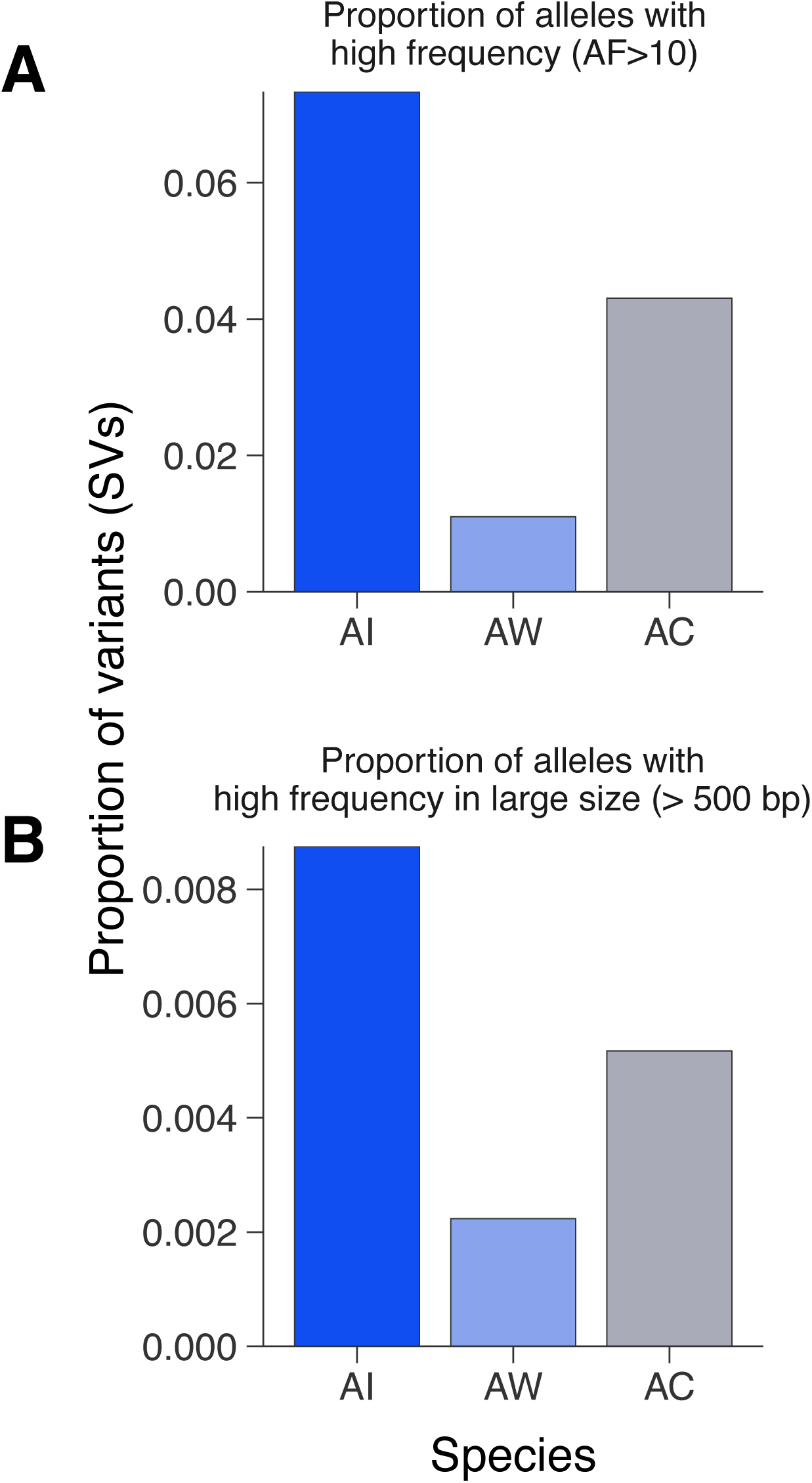
Frequencies of SVs in AI, AW and AC birds. **A**, proportion of derived SV alleles with allele count > 10 across species. **B**, proportion of large (> 500 bp) SV alleles with allele count > 10 across species.

**Fig. S36.**
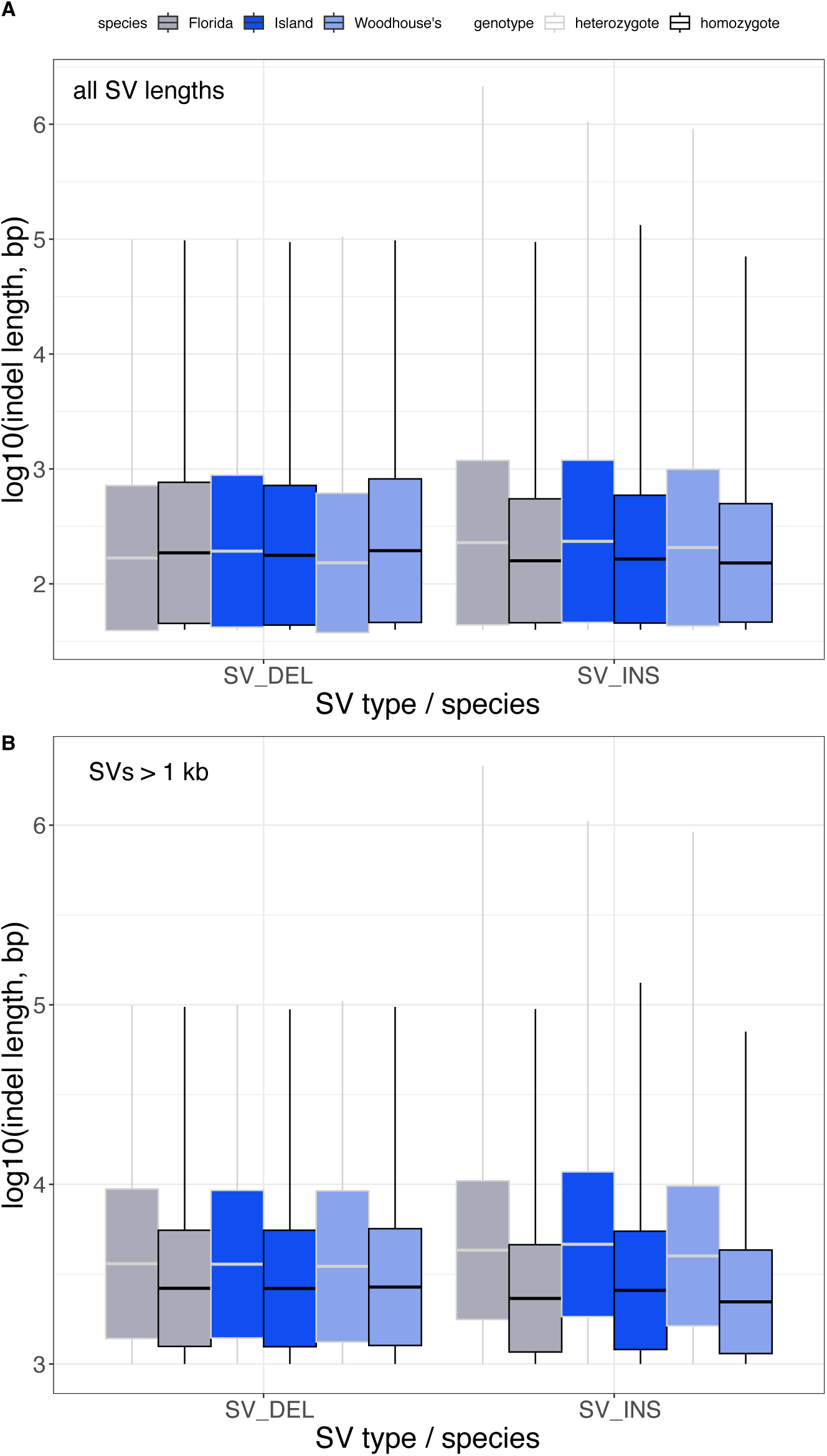
Distribution of indel lengths among homozygotes and heterozygotes. **A**, SVs of all lengths (> 50 bp). **B**, SVs > 1 kb. All birds were genotypes with svim-asm in diploid mode (Heller and Vingron 2021). All pairwise comparisons of genotypes within species are significant by a t-test (p < 0.05).

**Fig. S37.**
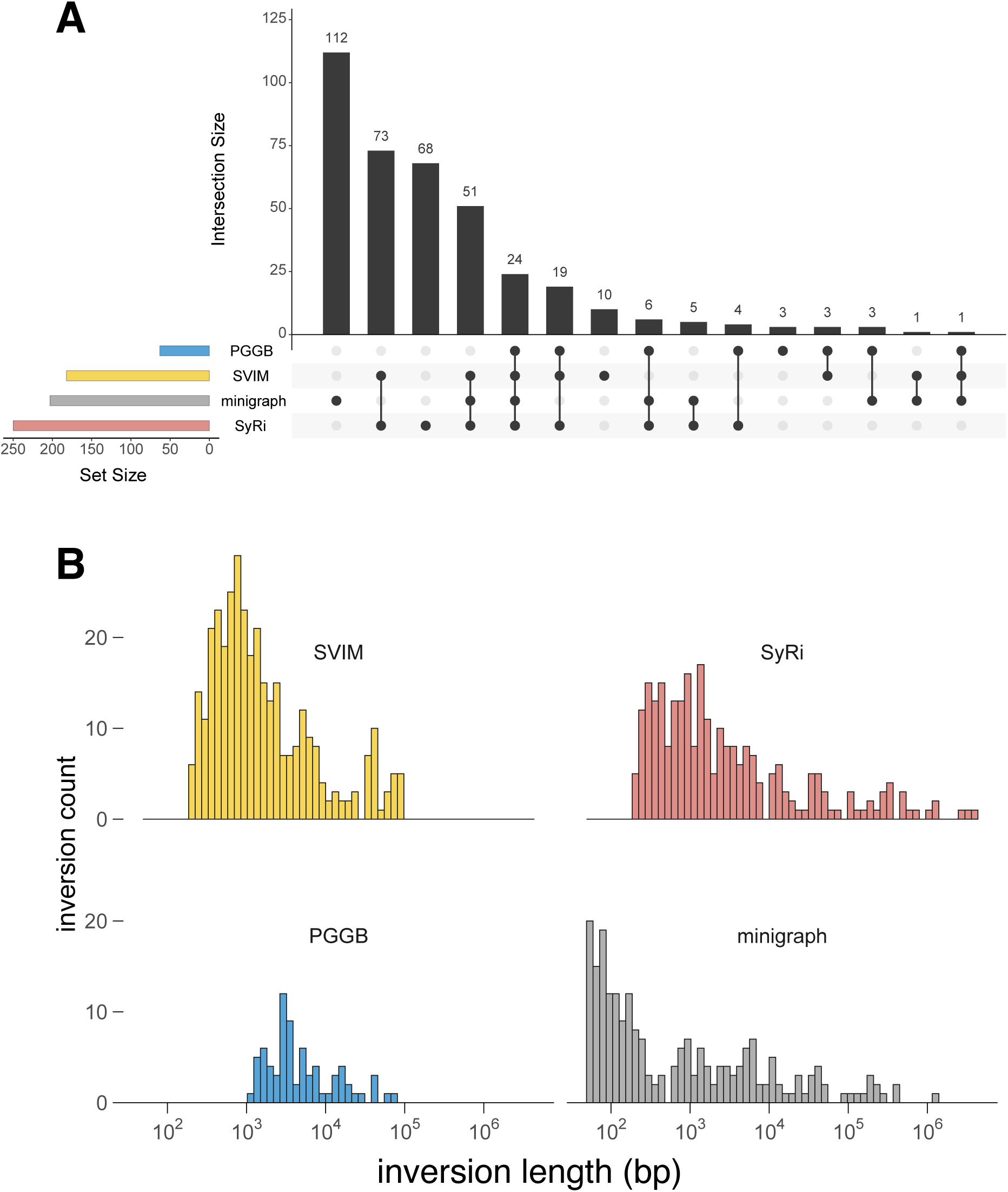
Sharing and distribution of inversion lengths detected by four different programs. **A**, sharing of inversions detected by PGGB, svim-asm, Syri and minigraph. **B**, distribution of inversion lengths detected by the four programs.

**Fig. S38.**
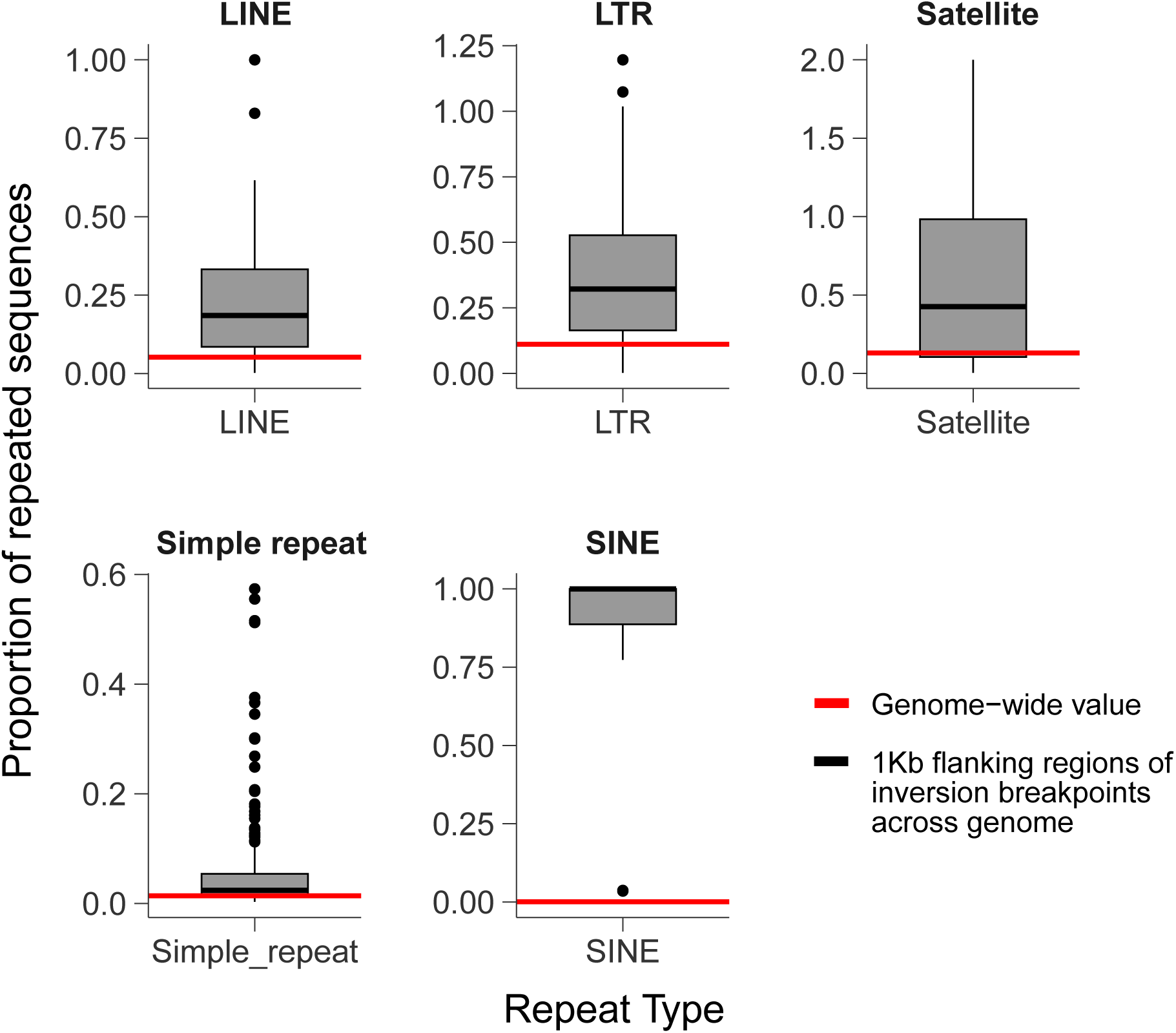
Enrichment of repeats in inversion flanking regions. Each boxplot depicts the proportion of repeat sequences in the 1-kb regions flanking the 95 inversions common to three software programs (see Methods). The red line indicates the genome-wide averages.

**Fig. S39.**
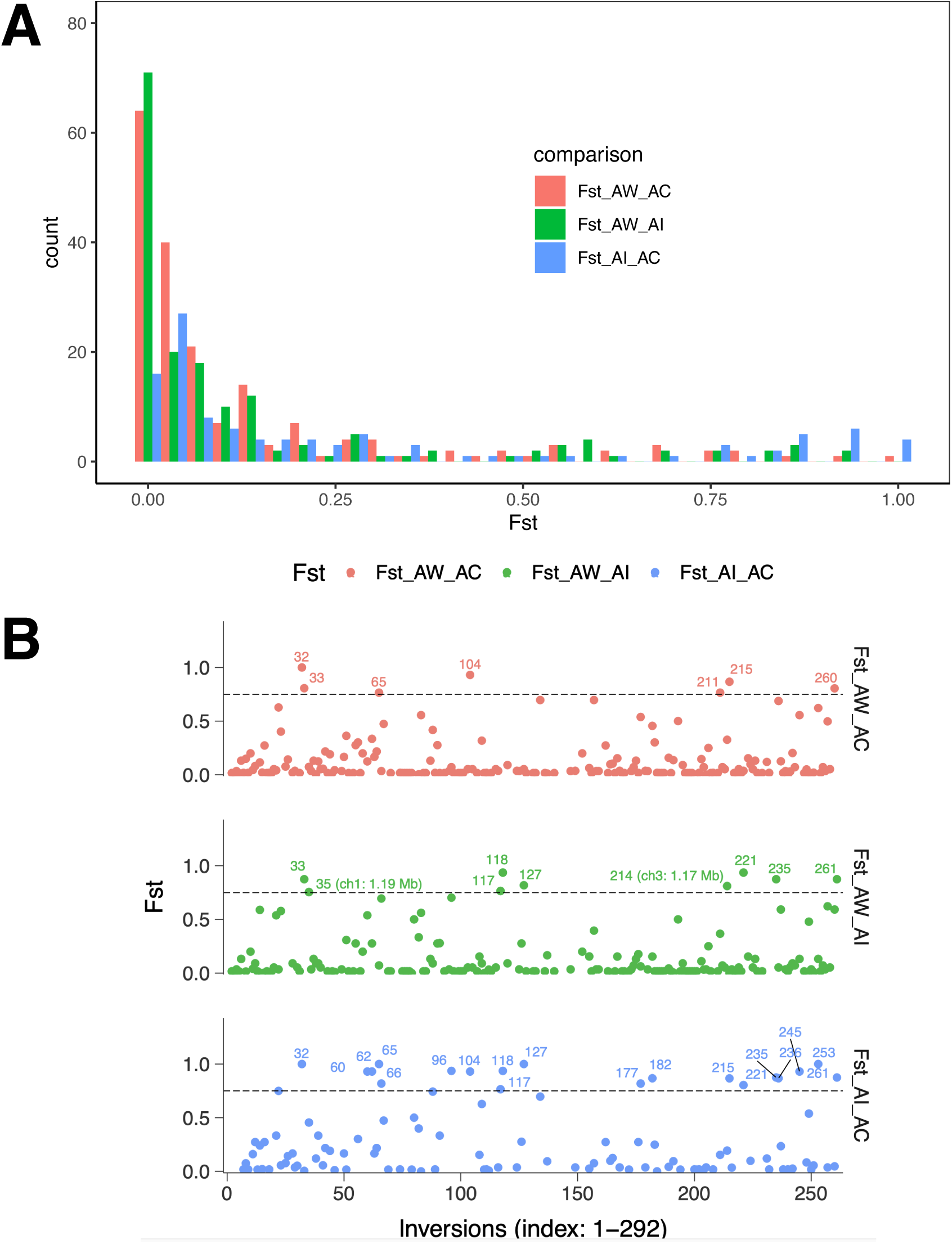
Inversion sharing and differentiation between species. **A**, histogram of inversion frequency differences (Fst) between pairs of species. **B**, same data as in A, the distribution of Fst among inversions for the three pairs of species. Specific inversions achieving high differentiation (> 0.75) are indicated with their index number and length for those > 1 Mb.

**Fig. S40.**
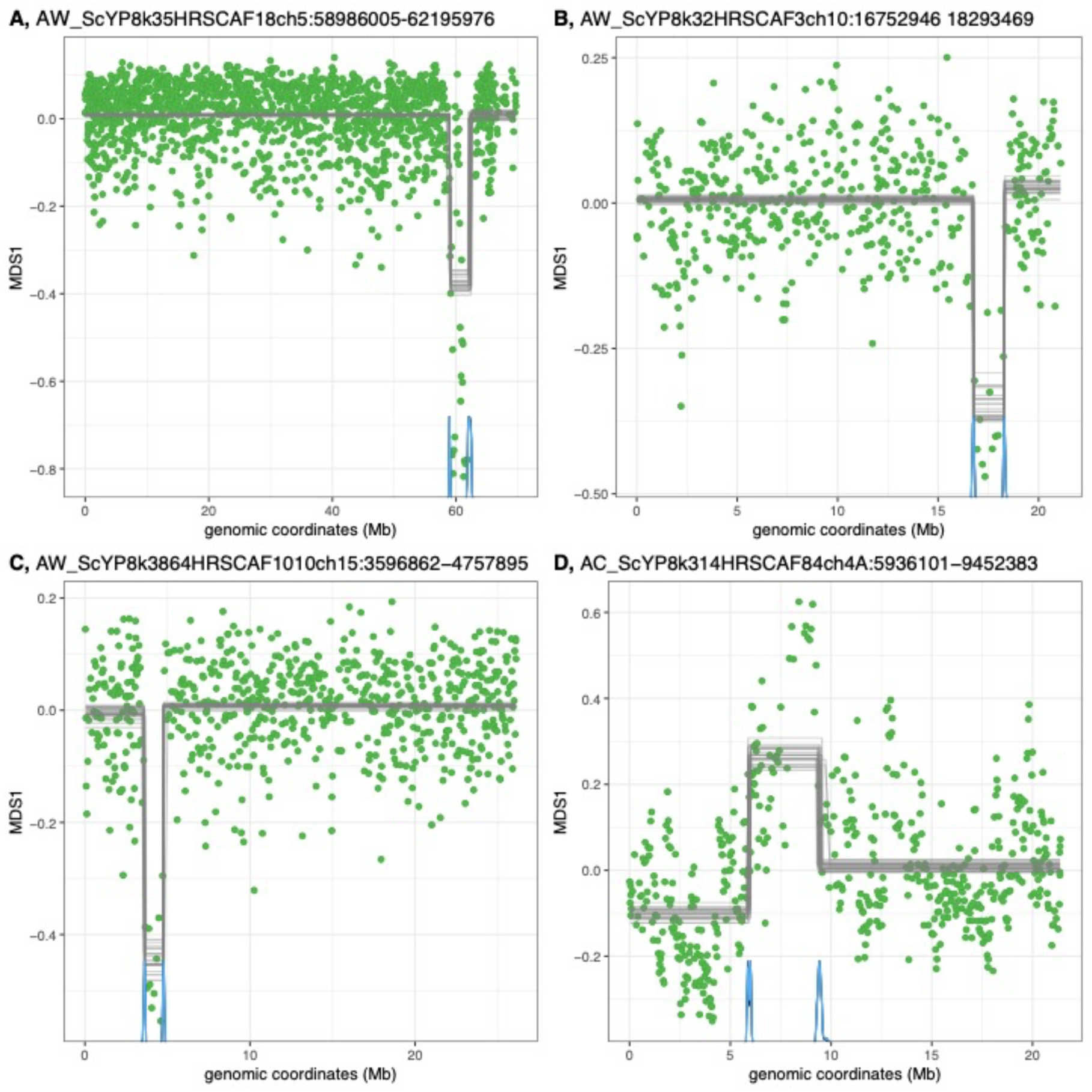
Four large inversions detected with lostruct. A, 3.2 Mb inversion on AW chr 5. B, 1.5 Mb inversion on AW chr 10. C, 1.2 Mb inversion on AW chr 15. D, 3.5 Mb inversion on AC chr 4A. On each plot, the green dots indicate the MDS1 value for the 5 kb or 50 kb window generated by lostruct (Li and Ralph 2019). The gray lines indicate Bayesian posterior distributions of breakpoints estimated by mcp (Lindeløv 2020). The blue densities at the bottom of each plot show the 95% c.i.s of each breakpoint.

**Fig. S41.**
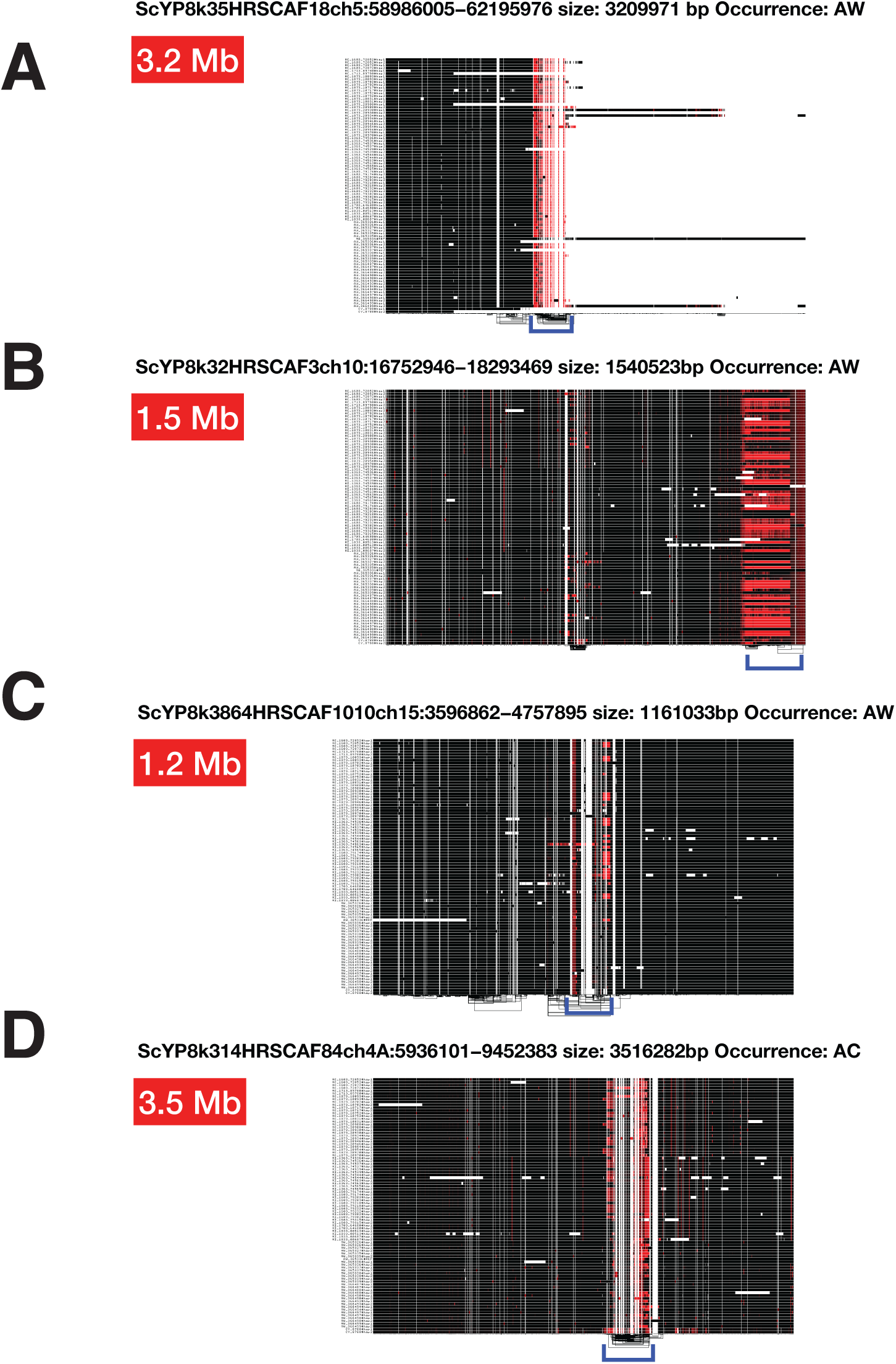
Visualizations of four large inversions using odgi viz. A-D, four putative inversions. Odgi (Guarracino et al. 2021) was used to visualize segments of the PGGB pangenome graph implicated in putative inversions detected by lostruct, using the - **S**flag. In each panel, black indicates alignment on the + strand and red on the - strand. The blue bracket below each plot indicates the putative inversion, visualized as changes from black to red (and back) in the 1D plots.

**Fig. S42.**
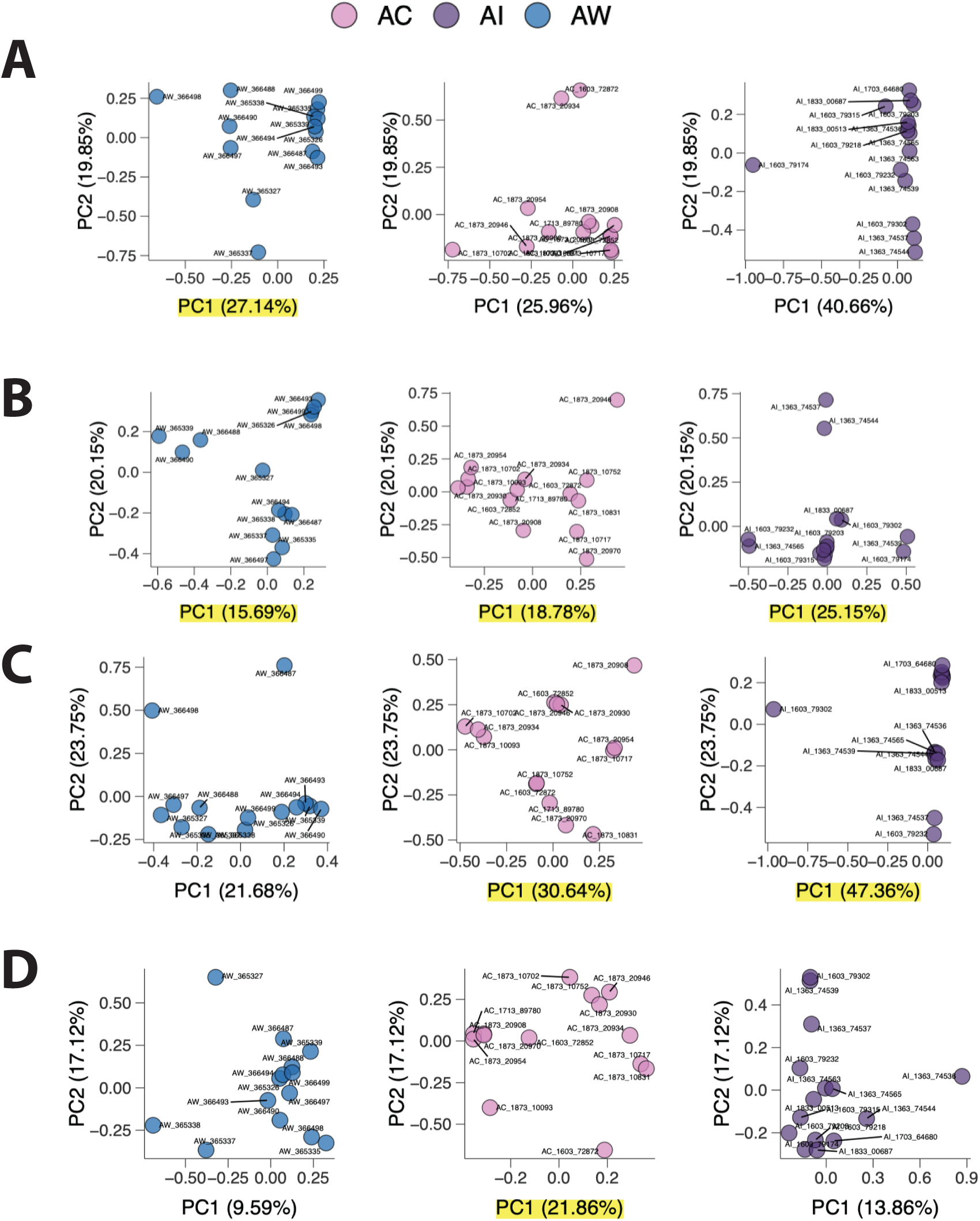
PCA analysis of genotypes across four large inversions. A-D, PCA plots of individuals of AW, AC, and AI for each of four putative inversions. PC1 and 2 scores were generated from each individual using the PGGB VCF and a bedfile of the inversion in question. Individual identities are indicated to demonstrate that it is not the same set of individuals that are achieving high PC1 scores for different inversions.

**Fig. S43.**
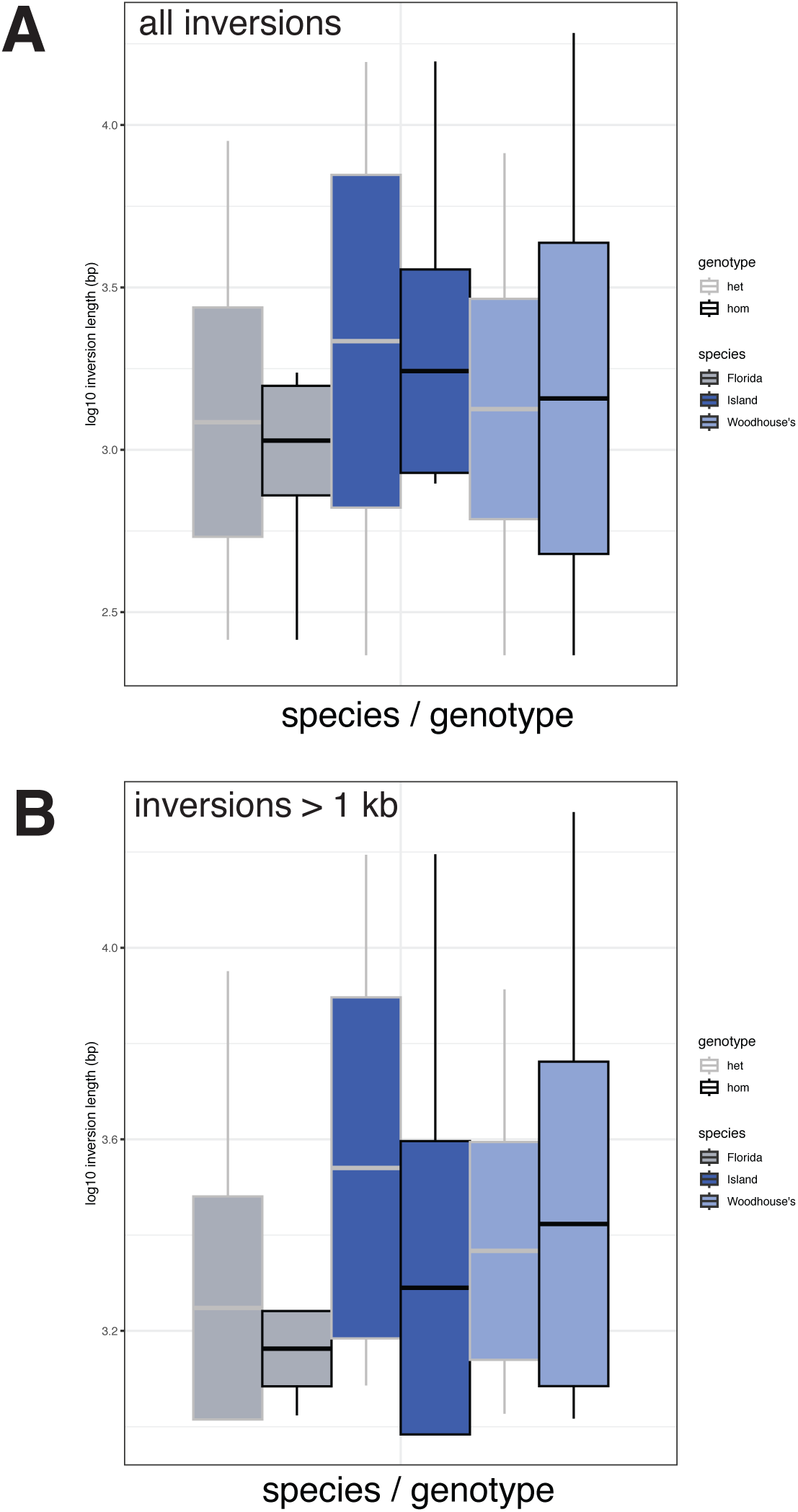
Distribution of inversion lengths among genotypes and species. **A**, boxplot of average log10(lengths) of inversions of all lengths in homozygotes and heterozygotes, as detected by svim-asm (Heller and Vingron 2021). No pairwise tests of mean log10 inversion length by genotype are significant by t.test. **B**, boxplot of average log10 lengths of inversions greater than 1 kb in homozygotes and heterozygotes. Only in AI were pairwise tests of mean log10 inversion length by genotype significant by t.test (p = 0.008).

**Fig. S44.**
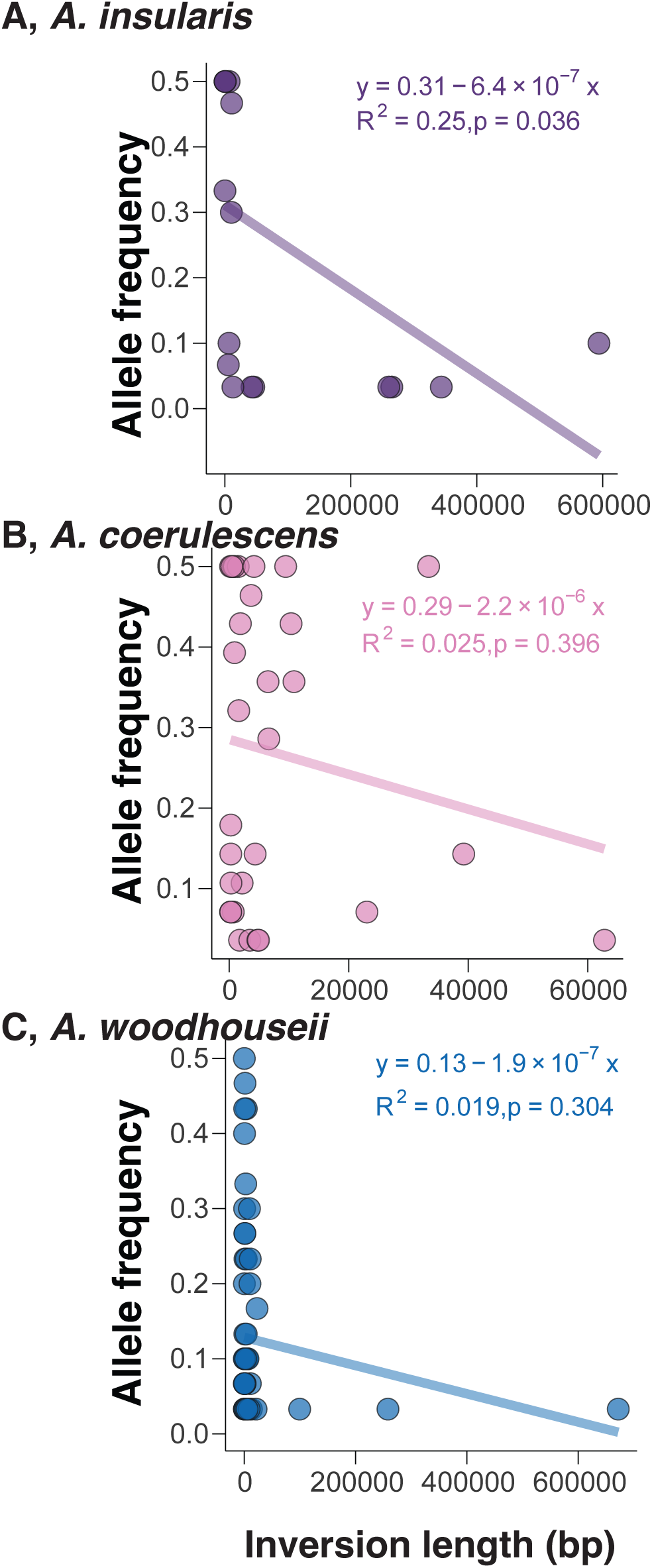
Inversion frequency within species as a function of inversion length. A, AI. B, AC. C, AW. Equations relating inversion length to frequency are provided for each plot.

**Fig. S45.**
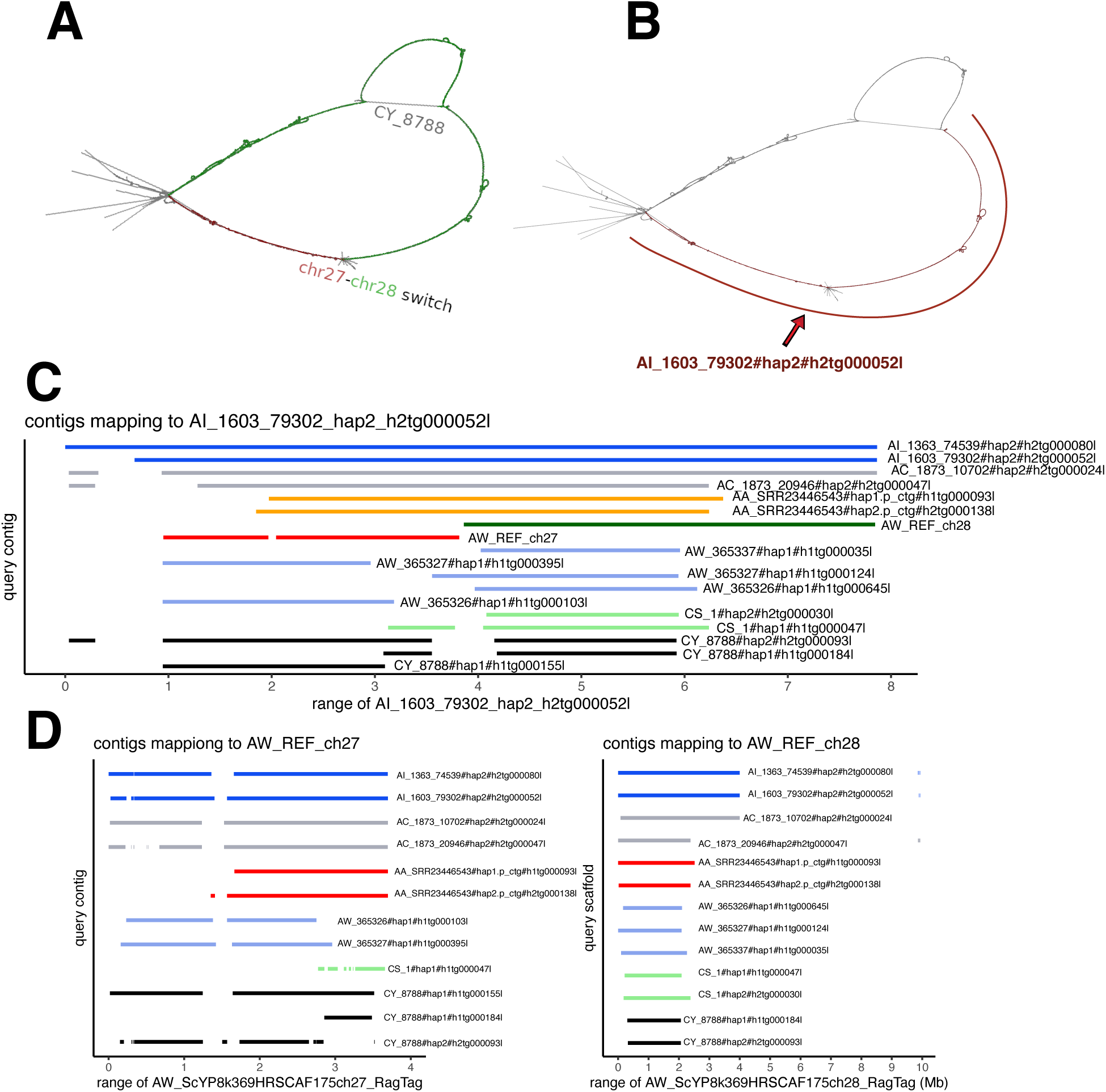
Characterization of a chromosomal fission in *A. woodhouseii*. A, bird-like 2D layout of the AW reference chr 27 / chr 28 region (PGGB community 14). The layout is colored red for AW ch 27 and green for AW ch28. B, 2D layout of PGGB community 14 with a long AI contig (contig AI_1603_79302#hap2#h2tg000052l) indicated, showing that it spans the AW ch27 / ch28 junction. C, minigraph mapping of diverse contigs to the long AI contig spanning the AW ch27 / ch28 junction (AI_1603_79302#hap2#h2tg000052l). Each horizontal line represents a single hifiasm contig mapping to the same same strand of the AI target for at least 250 kb. All map qualities scores are 60. D, minigraph mapping of diverse contigs to the AW reference chr 27 / chr 28 chromosomes. Each horizontal line represents a single hifiasm contig mapping to the same same strand of the AI target for at least 250 kb. All map qualities scores are 60. Note how the AW contigs mapping to ch27 and ch28 are different, whereas those from other species span the two AW chromosomes.

**Fig. S46.**
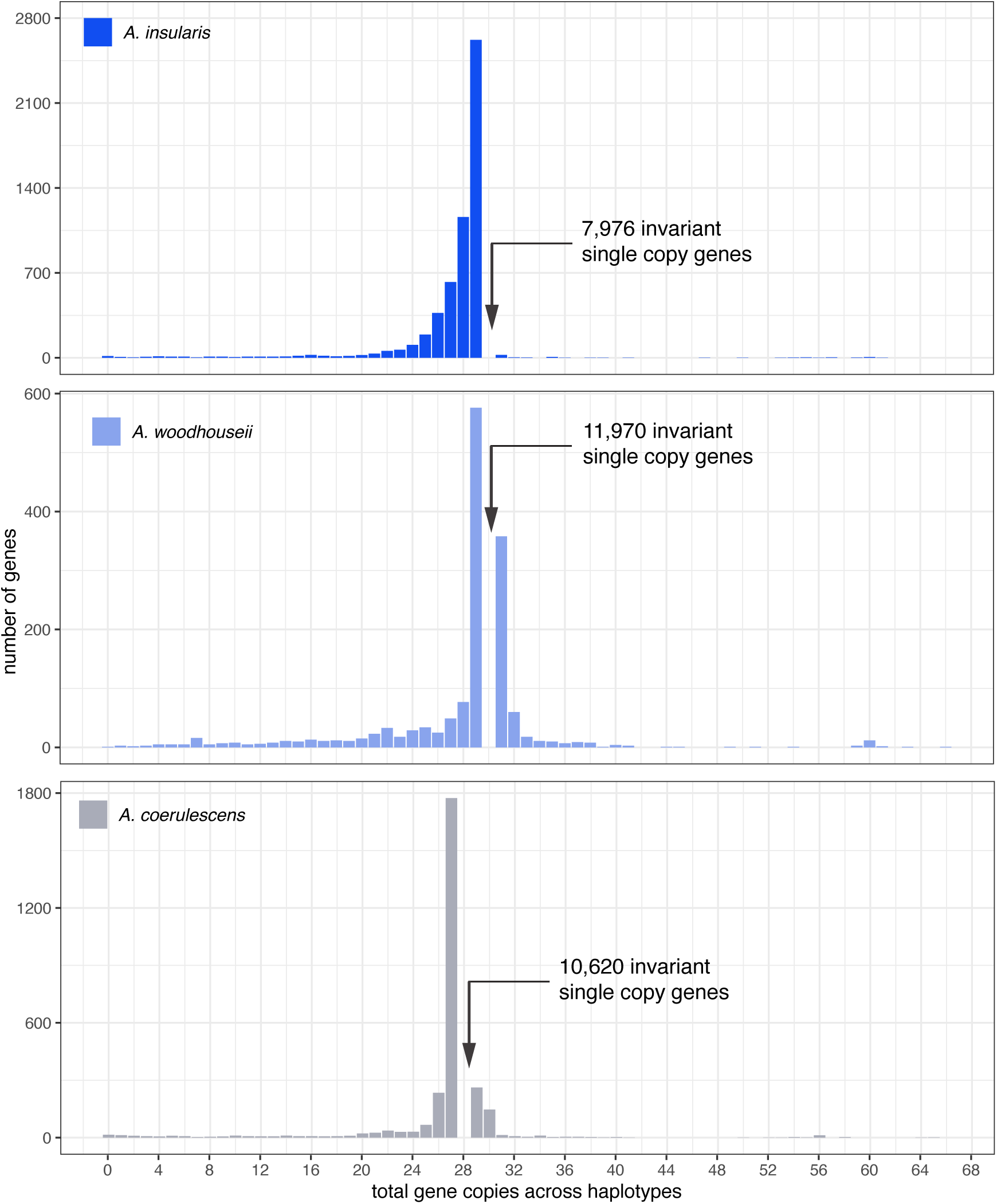
Distribution of total gene copies per haplotype by pangene. The large number of single copy genes are not plotted to allow better visualization of the non-single copy categories. The numbers of invariant single-copy genes are indicated for each species. Note the larger number of genes exhibiting deletions in AI, and the smaller numbers of multiplications of genes in AI versus AW and AC. All numbers reflect no filtering of pangene results (see table S14).

**Fig. S47.**
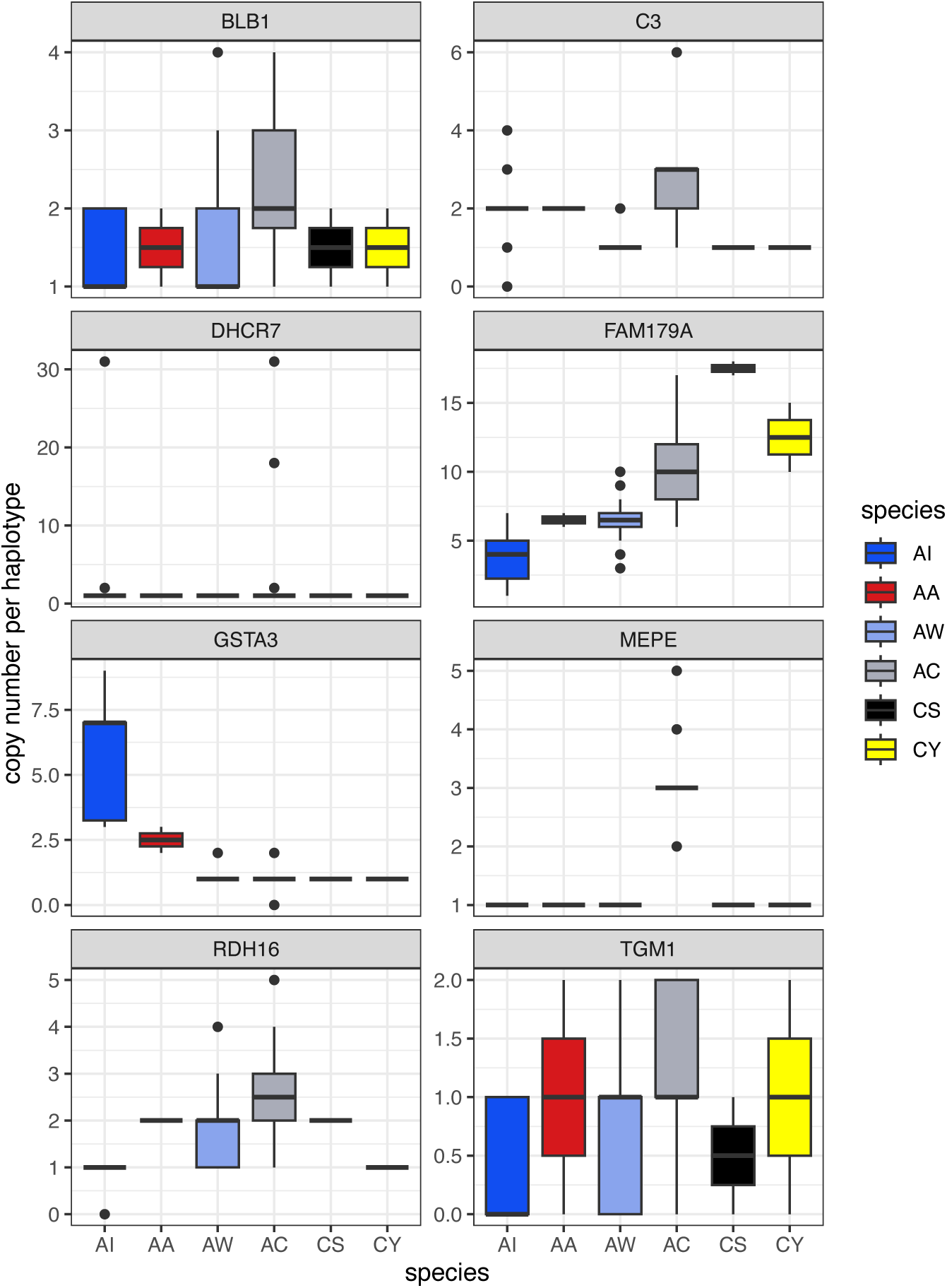
Examples of interspecific variation in gene copy number using pangene. Eight genes exhibiting significant or noteworthy variation in gene copy numbers between species. Species codes are indicated at the bottom of each plot. BLB1 corresponds to the major histocompatibility complex class II B genes.

## Notes

https://www.ncbi.nlm.nih.gov/bioproject/1206191

### Competing Interest Statement

The authors have declared no competing interest.

